# PATZ1 Reinstates a Growth-Permissive Chromatin Landscape in Adult Corticospinal Neurons After Injury

**DOI:** 10.1101/2025.05.14.654166

**Authors:** Anisha S. Menon, Manojkumar Kumaran, Deepta Susan Beji, Dhruva Kumar Kesireddy, Netra Krishna, Shringika Soni, Yogesh Sahu, Sneha Manjunath, Katha Sanyal, Meghana Konda, Ishwariya Venkatesh

## Abstract

**Background:** The failure of axon regeneration in the adult central nervous system represents a major barrier to recovery from spinal cord injury and neurodegenerative disease. Pro-growth transcription factors can promote regenerative responses, but their effects remain partial, suggesting that additional restraints must be relieved for these factors to achieve their full potential. The chromatin landscape of adult neurons has emerged as a candidate mechanism, yet we lack a developmental map of when and how this epigenetic restriction occurs, whether injury can reverse it, and how to therapeutically target it.

**Results:** We assembled a comprehensive chromatin accessibility atlas spanning mouse forebrain development from embryonic day 11 through adulthood using bulk and single-nucleus ATAC-seq. This revealed progressive restriction of growth-gene promoters and enhancers across postnatal development, leaving over 95% of growth-associated regulatory elements substantially inaccessible in mature neurons. We found that the distance between injury site and neuronal cell body determines the magnitude of chromatin reopening: intracortical lesions proximal to motor cortex soma triggered ten-fold greater enhancer reactivation compared to distal thoracic spinal cord crush. Motif analysis identified PATZ1, a chromatin-remodeling transcription factor, as correlated with this proximity effect. Viral delivery of PATZ1 to adult cortex converted the limited epigenomic response to distal injury into a profile approaching that of proximal injury, selectively reopening enhancers at growth-associated loci and depositing active H3K27ac marks. Hi-C analysis demonstrated that PATZ1 additionally reorganizes higher-order chromatin architecture, inducing compartment switching at growth loci and remodeling topologically associating domain boundaries. Integration with single-nucleus transcriptomics revealed that while PATZ1 selectively opens chromatin at growth genes, transcriptional output and axon regeneration remain modest, indicating that combinatorial approaches pairing epigenetic priming with pro-growth transcription factors may be required for functional repair.

**Conclusions:** This study provides a developmental timeline of chromatin closure at regeneration-associated genes and identifies PATZ1 as a molecular tool capable of reversing this epigenetic barrier in adult neurons. Our findings indicate that chromatin accessibility functions as a gatekeeping mechanism that must be addressed before transcription factor-based therapies can achieve their full effect, establishing epigenetic priming as a targetable component of CNS repair strategies.

## Introduction

The adult central nervous system (CNS) is largely incapable of self-repair, leaving traumatic brain and spinal injuries, as well as many neurodegenerative disorders, with few restorative options [1–3]. This regenerative failure reflects not only an inhibitory extracellular environment but also intrinsic brakes that tighten as neurons mature [1,2,4–12]. One of the most stringent intrinsic barriers is a progressive closure of chromatin around growth-associated loci, which sequesters regenerative programmes from the transcriptional machinery [13,14]. Broad histone-deacetylase inhibitors offer only limited benefit, presumably because they act indiscriminately on the neuronal epigenome [14]. A precise means of re-opening chromatin specifically at pro-growth regions, therefore, remains a major unmet need.

Although decades of work have illuminated transcription-factor networks that promote CNS axon outgrowth [4,5,7,15–27], more recent studies identify epigenetic regulation as an additional and decisive layer of control [28–35]. In zebrafish retinal ganglion cells and in mammalian peripheral neurons, axotomy triggers selective chromatin relaxation at growth-associated loci, permitting binding of pro-regenerative transcription factors [19,22,31,36,37]. By contrast, when central axons are injured, their parent somata mount little or no comparable epigenomic response. Conversely, artificially disrupting chromatin accessibility in otherwise regeneration-competent neurons is sufficient to block repair, together implying that the epigenetic barrier, not the injury signal alone, is rate-limiting for CNS regeneration repair [36,37]. Yet we still lack a developmental map of chromatin closure in CNS projection neurons, an understanding of whether injury distance influences their epigenetic plasticity, and a strategy to target chromatin opening without broad collateral effects.

Here, we chart chromatin accessibility in mouse corticospinal neurons from embryogenesis through adulthood and after spinal cord injury. We find that growth-gene loci close in two discrete post-natal waves (P0–P4 and P7–adult) and that injury delivered near the soma elicits a ten-fold stronger chromatin response than an equivalent lesion in the thoracic cord. Motif and footprint analyses identify PATZ1, a GC-binding zinc-finger protein of the stripe factor family[38], which coordinates broad transcriptional programmes during neural development[39], as abundant during the growth-competent period, lost in adults, and selectively redeployed following proximal injury. Viral delivery of PATZ1 to adult cortex after thoracic crush re-opens thousands of enhancers, installs H3K27ac at growth genes and reshapes 3-D genome architecture, thereby converting a muted distal-injury response into a developmental-like growth state. These findings position PATZ1 as a tractable lever for targeted chromatin remodeling and suggest that pairing stripe-factor delivery with existing pro-regenerative interventions could advance functional repair of the adult CNS.

## Results

### Two discrete postnatal contractions of chromatin accessibility at axon-growth genes

We first asked when forebrain neurons surrender the open chromatin landscape that supports developmental elongation. Public ENCODE bulk ATAC-seq datasets span embryonic day 11 (E11) to birth, revealing a progressive rise in chromatin accessibility peaking around E18, with a notable reduction by adulthood; however, these datasets are sparse in the postnatal window. We performed additional bulk ATAC-seq on forebrain tissue at P0, P4, P7, and adult timepoints to extend the developmental series (Figure 2A; Additional File 1-Table S2). Both ENCODE and in-house datasets were generated from forebrain tissue. To enable comparison between datasets, we applied batch correction using a negative binomial regression framework, with successful correction confirmed by PCA (Additional File 2-Figure S1-A; Additional File 1-Table S1; see Methods for details). Because ENCODE and in-house data were generated under different conditions, we present them in separate panels and interpret them as complementary rather than directly comparable (Figure 1A-B). We acknowledge that bulk ATAC-seq on heterogeneous forebrain tissue captures accessibility across multiple cell types, and that cell-type-specific changes may be diluted or masked by other populations. However, bulk accessibility profiles can still reveal broad developmental trends in chromatin architecture, and harnessing publicly available ENCODE data allowed us to maximise temporal coverage without duplicating existing resources.

**Figure 1.**
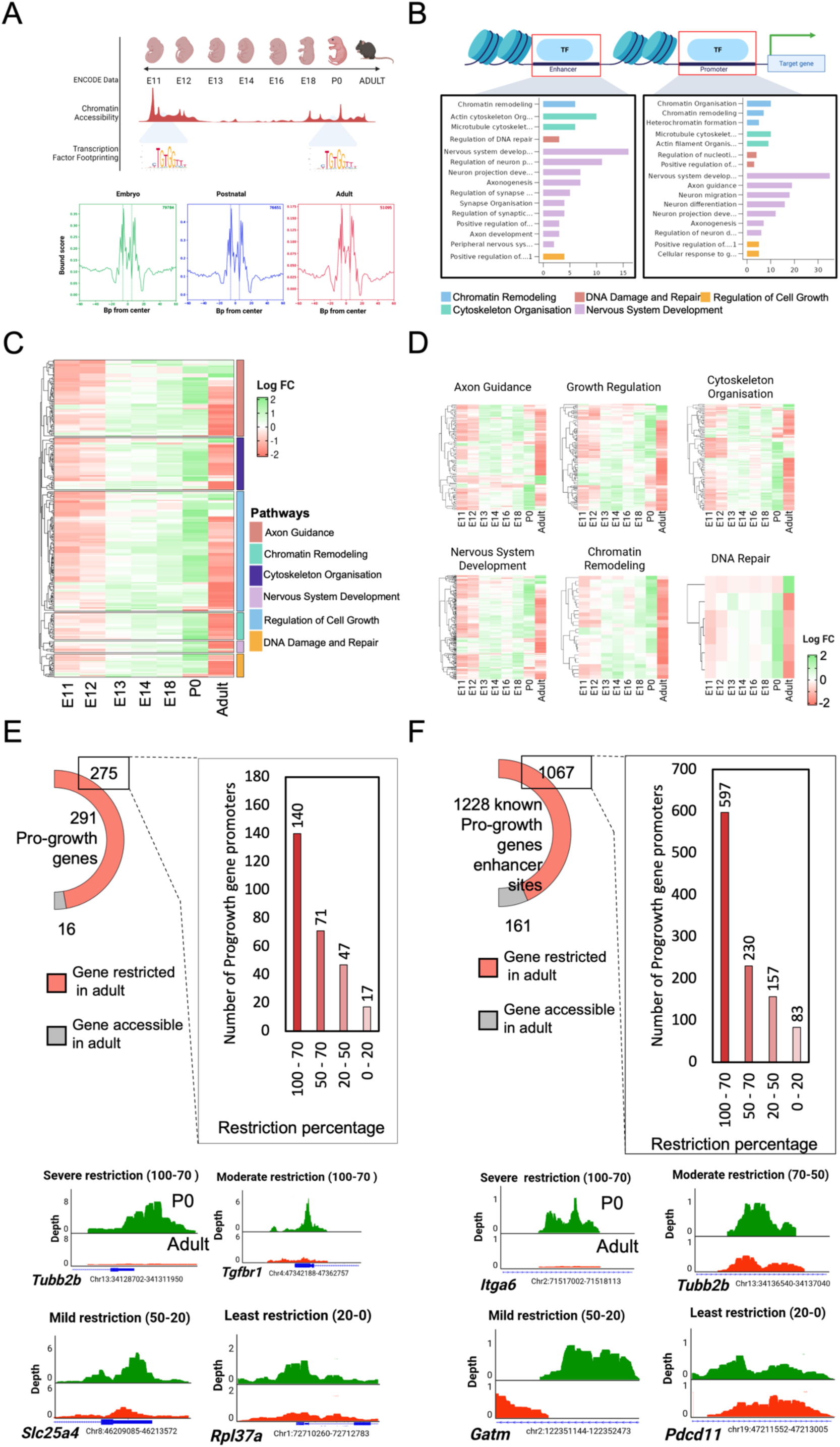
ATAC-seq reveals severe restriction of promoter and enhancer accessibility in pro-growth genes across age. (A) Top panel: Schematic diagram of ENCODE ATAC-seq datasets isolated from the motor cortex of mice from early embryonic ages, Postnatal Day 0 (P0), and adult mouse cortex were analysed for genome-wide patterns in chromatin accessibility. Bottom panel: Representative traces of transcription factor footprinting across age. (B) Pro-growth gene-specific promoters and enhancers were isolated for further analysis based on Gene Ontology (GO) Analysis. (p-value <0.05). (C) Heatmap of promoter accessibility across ages, showing high accessibility at P0 (green) neurons and reduced accessibility in adults (red) (p-value <0.05). (D) Heatmap of enhancer accessibility across age for growth-related GO terms (p-value <0.05). (E) Quantification and representative examples of age-dependent chromatin accessibility restriction at pro-growth gene promoters. (Left) Donut chart showing that of 291 pro-growth genes analysed, 275 have promoters that become restricted in the adult motor cortex (pink), while 16 retain accessibility in the adult (grey). (Centre) Bar graph categorising the 275 restricted promoters by the degree of accessibility loss relative to P0, across four restriction bins: Severe (100–70%, n=140), Moderate (70–50%, n=71), Mild (50–20%, n=47), and Least (20–0%, n=17). (Right) Representative UCSC Genome Browser tracks showing ATAC-seq chromatin accessibility signal (y-axis: read depth) at pro-growth gene promoters across developmental timepoints, with the P0 condition shown in green (open chromatin) and the adult condition shown in red (restricted chromatin). (F) Quantification and representative examples of chromatin accessibility restriction at pro-growth gene enhancer sites. (Left) Donut chart showing that of 1,228 known pro-growth gene enhancer sites analysed, 1,067 become restricted in the adult motor cortex (pink), while 161 remain accessible in the adult (grey). (Centre) Bar graph categorising the 1,067 restricted enhancers by the degree of accessibility loss relative to P0, across four restriction bins: Severe (100–70%, n=597), Moderate (70–50%, n=230), Mild (50–20%, n=157), and Least (20–0%, n=83). (Right) Representative UCSC Genome Browser tracks showing ATAC-seq chromatin accessibility signal (y-axis: read depth) at pro-growth gene enhancer loci, with P0 shown in green and adult in red.

**Figure 2:**
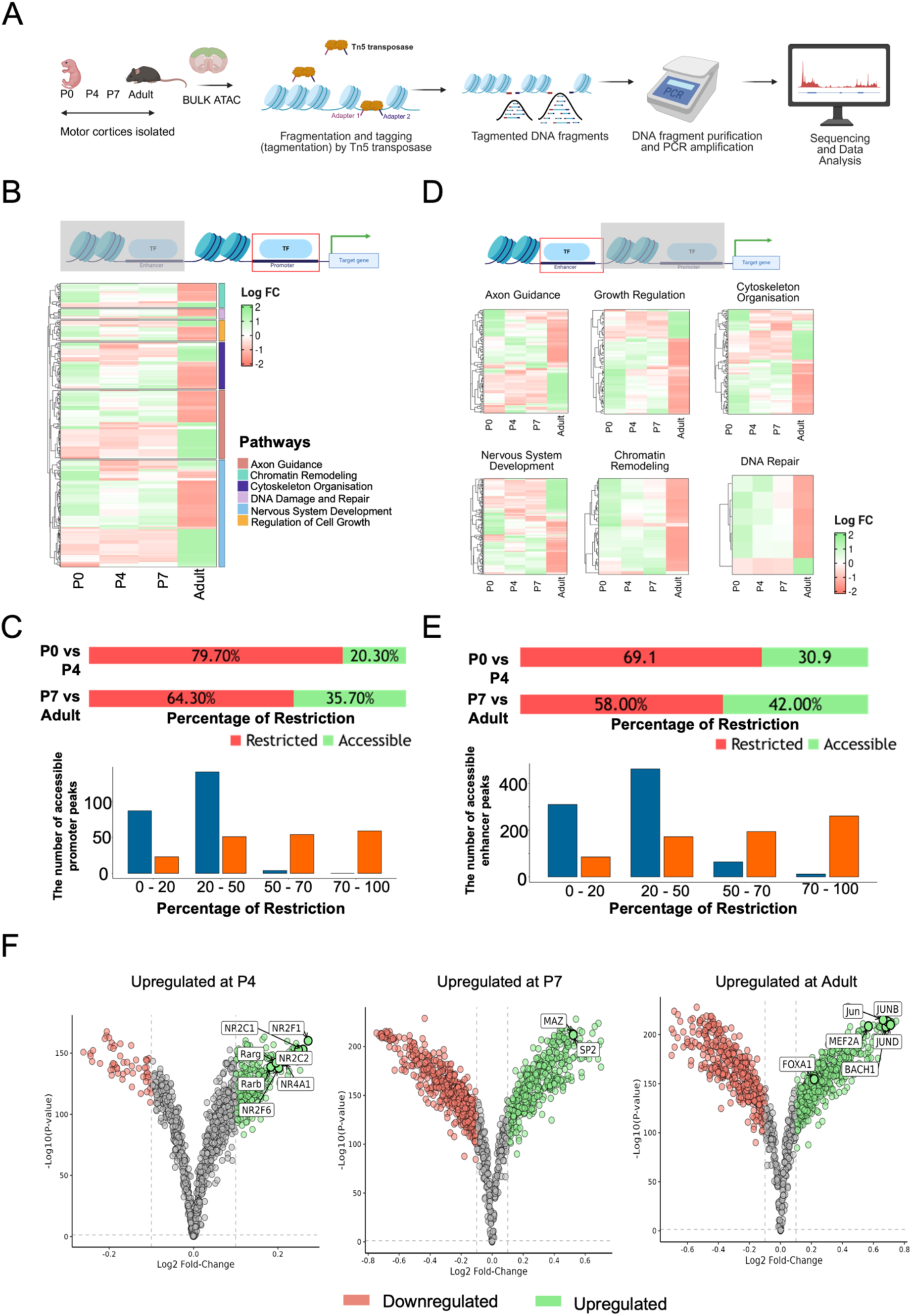
ATAC-seq reveals restriction of promoter and enhancer accessibility in pro-growth genes across post-natal stages. (A) Schematic of the bulk ATAC-seq experimental workflow. Motor cortices were isolated from mice at postnatal day 0 (P0), P4, P7, and adult stages. Chromatin was fragmented and tagged by Tn5 transposase, which simultaneously inserts sequencing adapters at accessible genomic regions. Fragmented DNA was purified and amplified by PCR, followed by sequencing and bioinformatic analysis. (B) (Top) Schematic illustrating that promoter regions were isolated and analysed for chromatin accessibility changes. (Bottom) Hierarchically clustered heatmap showing scaled fold-change (Scaled FC) in promoter chromatin accessibility across developmental timepoints (P0, P4, P7, Adult) for pro-growth genes grouped by biological pathway (colour-coded sidebar: Axon Guidance — blue; Chromatin Remodeling — purple; Cytoskeleton Organisation — teal; DNA Damage and Repair — dark purple; Nervous System Development — green; Regulation of Cell Growth — orange). Colour scale indicates Scaled FC (green = increased accessibility; red = decreased accessibility; range –2 to +2). Promoter accessibility is highest at P0 and shows progressive restriction from P4 onwards (p < 0.05). (C) Quantification of pro-growth gene promoter accessibility restriction across postnatal development. (Top) Horizontal stacked bar charts comparing the percentage of promoters that are restricted (red/dark) versus accessible (green) in two pairwise comparisons: P0 vs P4 (79.7% restricted, 20.3% accessible) and P7 vs Adult (64.3% restricted, 35.7% accessible). (Bottom) Bar graph showing the number of accessible promoter peaks (y-axis) distributed across four bins of restriction percentage (x-axis: 0–20%, 20–50%, 50–70%, 70–100%), comparing P0 vs P4 (blue bars) and P7 vs Adult (orange bars). In the P0 vs P4 comparison, peaks are predominantly concentrated in the lower restriction bins (0–50%), whereas in the P7 vs Adult comparison, a greater proportion of peaks shift into higher restriction bins (50–100%), indicating progressive promoter closing with age. (D) (Top) Schematic illustrating that enhancer regions were isolated and independently analysed for accessibility changes. (Bottom) Pathway-specific heatmaps showing Scaled FC in enhancer chromatin accessibility across developmental timepoints (P0,P4,P7,Adult) for six GO-term-defined pathways: Axon Guidance, Growth Regulation, Cytoskeleton Organisation, Nervous System Development, Chromatin Remodeling, and DNA Repair. Colour scale as in (B) (green = open; red = closed). Enhancer chromatin accessibility is highest at P0 and undergoes progressive restriction through postnatal development into adulthood (p < 0.05). (E) Quantification of pro-growth gene enhancer accessibility restriction across postnatal development. (Top) Horizontal stacked bar charts comparing the percentage of enhancers that are restricted (red/dark) versus accessible (green) in two pairwise comparisons: P0 vs P4 (69.1% restricted, 30.9% accessible) and P7 vs Adult (58.0% restricted, 42.0% accessible). (Bottom) Bar graph showing the number of accessible enhancer peaks (y-axis) distributed across four restriction percentage bins (x-axis: 0–20%, 20–50%, 50–70%, 70–100%), comparing P0 vs P4 (blue bars) and P7 vs Adult (orange bars). Consistent with the promoter data in (D), the P7 vs Adult comparison shows a relative increase in peaks at higher restriction levels compared to the P0 vs P4 comparison, reflecting ongoing enhancer silencing at pro-growth loci through postnatal maturation. (F) Volcano plots showing stage-specific transcription factor (TF) footprinting analysis, identifying TFs with significantly enriched occupancy (upregulated, green) or reduced occupancy (downregulated, red) at accessible chromatin regions relative to the preceding stage. Each dot represents one TF motif; x-axis shows Log2 fold-change in footprint score and y-axis shows –log10(adjusted p-value). Dashed vertical lines indicate fold-change thresholds. Labelled TFs with significantly increased occupancy include: at P4 — NR2C1, NR2F1, NR2C2, NR4A1, RARG, RARB, and NR2F6; at P7 — MAZ and SP2; and at Adult — Jun, JUNB, JUND, MEF2A, FOXA1, and BACH1.

To examine chromatin dynamics at genes relevant to axon growth, we employed two complementary approaches. First, we curated 291 promoters and 1,228 enhancers flanking genes implicated in axon guidance, cytoskeleton dynamics, cell-growth regulation, chromatin remodeling, DNA-damage repair, and nervous-system development (Figure 1C-D; Additional File 1-Table S1). These categories were selected because axon guidance and cytoskeleton genes directly mediate growth cone navigation and extension; cell-growth regulators control the metabolic and signalling programmes required for elongation; chromatin remodellers may enable or constrain access to growth programmes; DNA-damage repair genes were included because they are upregulated upon treatment with pro-regenerative transcription factors such as KLF6 and Nr5a2, and because highly interacting chromatin regions, including enhancers and super-enhancers that form long-range loops, are enriched in DNA repair factors, suggesting that protection from double-strand breaks at these sites is essential during active transcription[40]; and nervous-system development genes encompass broad neurodevelopmental programmes.

We note two important methodological considerations regarding this enhancer set. First, the enhancer–gene associations were derived computationally using the Activity-By-Contact (ABC) algorithm[20,41], which integrates chromatin accessibility, H3K27ac signal, and Hi-C contact data to link distal regulatory elements to their predicted target genes. While the ABC model has been extensively benchmarked against perturbation-based enhancer–gene maps and shows high precision for active regulatory elements, these associations represent probabilistic predictions rather than experimentally validated links, and a subset may not reflect functional regulatory relationships in all cellular contexts. Second, the promoters and enhancers in this set were not independently characterised by reporter assay or CRISPR-based perturbation in the current study; their inclusion is based on genomic proximity to genes with established roles in axon growth and their classification as candidate regulatory elements by ENCODE annotation and ABC modelling.

Accessibility at these growth-gene promoters was not static. In early embryos (E11-E12), the vast majority of loci were only modestly accessible. Accessibility then rose monotonically, peaking between E18 and P0 in concert with the period of brisk CST outgrowth (Figure 1C; Additional File 1-Table S1). Instead of plateauing, accessibility fell during the postnatal ages. Quantitative modelling revealed two kinetically distinct contractions: a first wave between P0 and P4 that imposed mild-to-moderate restriction (20-50% loss of signal at 48.5% of promoters [141 of 291] and 37.7% of enhancers [463 of 1,228]), and a second wave between P7 and adulthood that produced more extensive closure, with at least 50% loss at 38.8% of promoters [113 of 291] and 37.1% of enhancers [455 of 1,228]. By maturity, 38.8% of promoters (113 of 291) and 37.1% of linked enhancers (455 of 1,228) had lost at least half of their peak accessibility, leaving the majority (61.2% of promoters [178 of 291] and 62.9% of enhancers [773 of 1,228]) partly open (Figure 2 B-E; Additional File 1-Table S2).

To illustrate the heterogeneity of chromatin closure at individual loci, we categorized promoters and enhancers by their degree of restriction in adulthood relative to peak accessibility at P0. Of the 291 curated growth-gene promoters, 140 showed severe restriction (100-70% loss), 71 showed moderate restriction (70-50% loss), 47 showed mild restriction (50-20% loss), and only 17 having the least restrictive promoter regions(less than 20% loss) in adults (Figure 1E). Representative genome browser tracks spanning these four restriction categories demonstrate the range of developmental closure patterns: genes such as *Tubb2b* exhibited near-complete loss of promoter accessibility by adulthood (severe restriction), whereas *Tgfbr1* and *Slc25a4*showed progressively milder degrees of closure, with *Rpl37a* maintaining substantial accessibility into maturity (Figure 1E, right panels, Additional File 1-Table S1).

Enhancers displayed similar stratification but with greater absolute numbers in each category. Of 1,228 growth-associated enhancers, 597 were severely restricted, 230 moderately restricted, 157 mildly restricted, and 83 remained least restricted in adults (Figure 1F). Genome browser examples at enhancer regions linked to *Itga6*, *Tubb2b, Gatm*, and *Pdcd1* illustrate these distinct restriction profiles (Figure 1F, right panels). Even within the same functional pathway, individual regulatory elements exhibited divergent closure kinetics, suggesting locus-specific regulation rather than uniform pathway-level control. Notably, axon-guidance enhancers displayed bidirectional accessibility patterns in adulthood, with adult-inhibitory loci (*Robo2*, *Ephb2*, *Cntn1*) retaining open chromatin while developmental pathfinding genes underwent progressive restriction, consistent with functional partitioning of the pathway (Additional File 2 – Figure S22).

We note that the P7-to-adult interval spans several weeks and likely encompasses gradual changes rather than a discrete transition; the exact timing of closure within this window remains undefined by our sampling. We therefore interpret P4 and P7 as informative boundaries rather than as precise developmental checkpoints. By P4, many growth loci have begun to restrict, and by P7, the majority show substantial closure, though residual accessibility persists into adulthood at some sites.

When we clustered accessibility trajectories, promoters behaved largely synchronously, whereas enhancers displayed pathway-specific variation. Between P0 and P4, 79.7% of promoters showed restriction compared to 69.1% of enhancers; by the P7 to adult transition, 64.3% of promoters and 58.0% of enhancers were restricted (Figure 2 C-E; Additional File 1-Table S2). Despite this broad closure, enhancers for nervous-system-development genes retained measurable openness into adulthood, with roughly one-quarter remaining above the 50% accessibility threshold, suggesting a need for ongoing plasticity in that pathway. Enhancers governing DNA-repair genes were even more refractory: nearly 30% retained embryonic-level accessibility in the adult. Beyond potentially supporting rapid lesion-induced repair, this persistent openness may preserve architectural flexibility required for long-range enhancer-promoter contacts that activate genome-maintenance pathways, though this hypothesis requires direct testing (Figure 1C-D; Additional File 1-Table S1).

TF-footprinting across the same series exposed changes in binding landscapes. The embryonic and early-postnatal period was dominated by orphan nuclear receptors (NR2C1/2, NR2F1/6) and RAR-related factors that coordinate early development (Figure 2F left; Additional File 1-Table S2). Between P4 and P7, these footprints diminished, replaced by strong occupancy of SP2, and MAZ (Figure 2F middle; Additional File 1-Table S2). By adulthood, JUN, JUNB, and MEF2A became the prevailing binders (Figure 2F right; Additional File 1-Table S2). The timing of these TF exchanges correlates with the two waves of chromatin contraction, which hints at active regulation of promoter-enhancer closure.

We extend the approach holistically to address curation bias through two mechanisms. First, to extend our study beyond growth loci, we examined chromatin accessibility across the entire mouse genome (hereafter referred to as the global gene set). Global gene promoters showed progressive restriction from embryonic stages through adulthood, mirrored by a proportional shift from accessible to restricted chromatin at the population level (Additional File 2 – Figure S19A–B). In contrast to the selective two-wave contraction observed at pro-growth loci, global genes showed a more gradual and less pronounced reduction, confirming that the restriction we describe is not a genome-wide phenomenon but is disproportionately concentrated at growth-associated regulatory elements. Second, to comprehensively profile chromatin accessibility without bias from manual curation, we retrieved complete gene sets associated with GO terms relevant to neuronal growth and plasticity [Signal transduction, Chromatin remodeling, Actin cytoskeleton organisation, Axon Guidance, Axonogenesis, Neurogenesis, central nervous system development, Neuron development, Axon extension, Regeneration and Neuron projection guidance]. Using their GO IDs, we compiled all genes annotated to these categories and generated heatmaps (Additional File 2 – Figure S1-B). The patterns observed from the unbiased GO-based gene sets were consistent with those from the curated list, supporting the robustness of our findings across both analytical frameworks. Finally, to establish that the observed chromatin restriction is functionally consequential rather than epigenomically inert, we integrated ATAC-seq accessibility with matched RNA-seq expression data across developmental stages. Confusion matrix analysis confirmed a strong genome-wide correspondence between open chromatin and active transcription (Additional File 2 – Figure S19C). Critically, this concordance was progressively lost specifically at pro-growth loci during postnatal development: embryonic pro-growth genes showed high chromatin–transcription agreement that diminished markedly in the adult (Additional File 2 – Figure S19D), establishing that developmental chromatin closure at these loci is accompanied by functional transcriptional silencing, not merely epigenomic reorganisation.

### A distal thoracic crush activates a modest, broadly distributed epigenomic response

Seven days after complete T8 crush, an interval when axonal degeneration has stabilised but spontaneous sprouts are still scarce, we isolated nuclei from motor cortex and generated 3’ single-nucleus ATAC-seq (snATAC-seq) libraries (Figure 3A). Representative histology confirming the crush is shown in Additional File 2 – Figure S5.To identify corticospinal-enriched populations, we performed anterograde AAV tracing one week before injury, injecting AAV-CAG-GFP into layer V motor cortex to label descending projection neurons. We then used fluorescence-activated nuclear sorting (FANS) to enrich for GFP-positive nuclei prior to library preparation (Additional File 2-Figure S2-A-D). Within the snATAC-seq data, we identified layer 5 extratelencephalic (L5 ET) neurons based on chromatin accessibility at marker genes including *Etv1*, *Crym*, and *Mrpl11* [42–44] (Figure 3B, Additional File 2-Figure S3-E-F). L5 ET neurons include corticospinal neurons but also other subcortical projection neurons; we therefore refer to this population as L5 ET throughout rather than as CST specifically[43].

**Figure 3.**
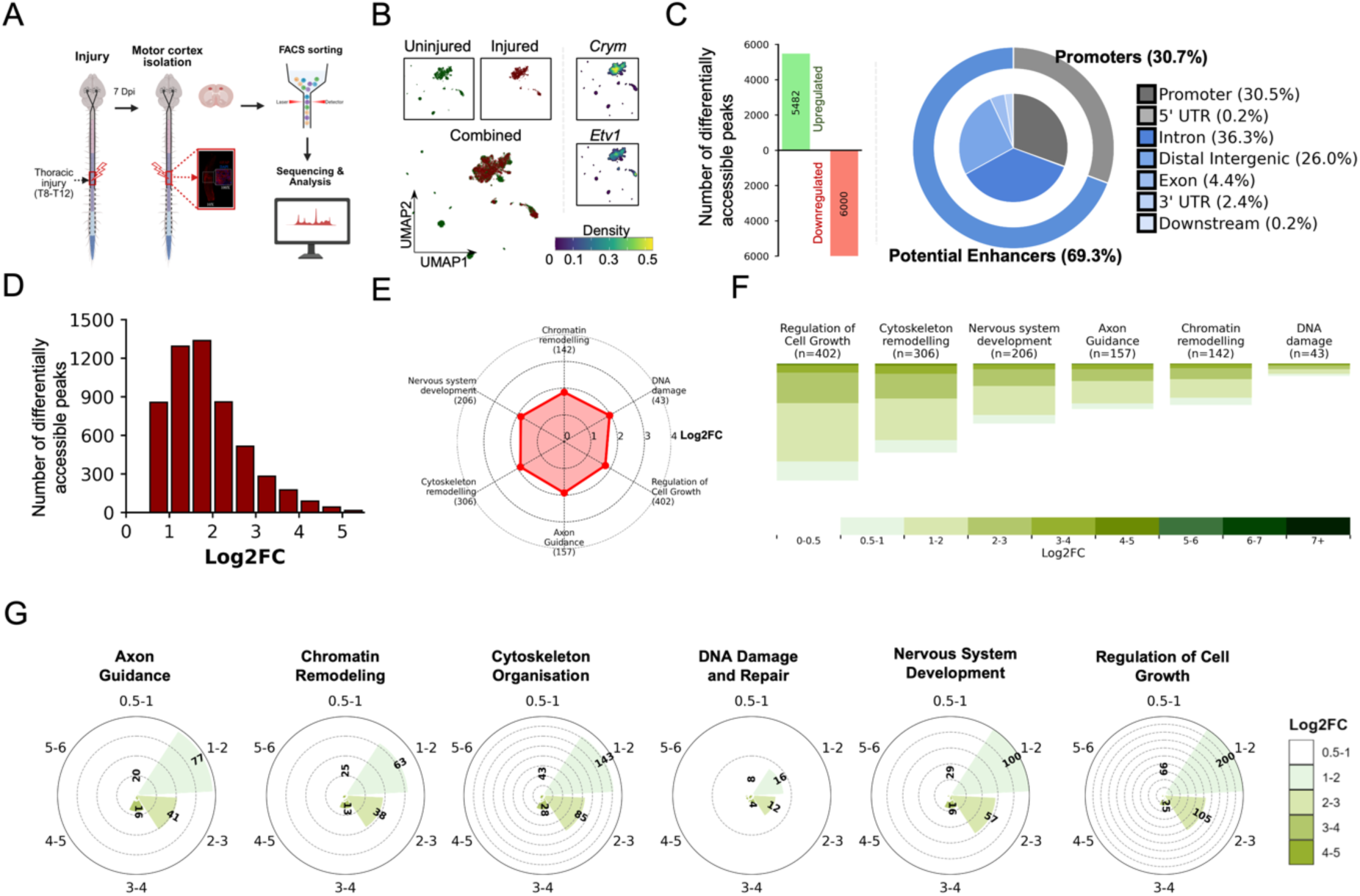
Single-nuclear (sn)ATAC-seq reveals that CNS neurons mildly open the chromatin post-injury at key loci. (A) Schematic of the snATAC-seq experimental workflow. Following thoracic spinal cord injury (7 days post-injury, dpi), motor cortex tissue was isolated and dissociated. Nuclei were sorted by FACS and processed for library preparation, sequencing, and downstream bioinformatic analysis. (B) UMAP plots showing snATAC-seq data from uninjured (green) and injured (red) conditions plotted separately and combined (bottom left). Dot density is indicated by the colour scale (0–0.5). Feature activity for the corticospinal neuron markers Crym and Etv1 (top right) was used to identify Layer 5 extratelencephalic (L5 ET) neuronal clusters for downstream analysis. (C) (Left) Bar graph showing the number of differentially accessible chromatin peaks in L5 ET neurons following injury: 5,482 upregulated (green, increased accessibility) and ∼6,000 downregulated (red, decreased accessibility). (Right) Donut chart showing the genomic distribution of the 5,482 upregulated peaks. The outer ring indicates that 30.7% of peaks fall within promoter regions; the inner pie chart provides a full breakdown: Promoter (30.5%), Intron (36.3%), Distal Intergenic (26.0%), Exon (4.4%), 3’ UTR (2.4%), 5’ UTR (0.2%), and Downstream (0.2%). Together, intronic, distal intergenic, and other non-promoter regions account for 69.3% of upregulated peaks, consistent with their classification as putative enhancer elements. (D) Histogram showing the Log2 fold-change (Log2FC) distribution of the 5,482 upregulated peaks. The majority of peaks display moderate increases in accessibility, with most falling in the Log2FC range of 1–3. (E) Radar (spider) chart illustrating the mean chromatin accessibility Log2FC across six GO-term-defined biological pathways in injured neurons: Chromatin Remodeling (n=142), DNA Damage (n=43), Regulation of Cell Growth (n=402), Axon Guidance (n=157), Cytoskeleton Remodeling (n=306), and Nervous System Development (n=206). The red shaded polygon represents the injured condition; concentric rings represent Log2FC values from 0 to 4. The chart enables comparison of the relative magnitude of chromatin opening across pathways. (F) Heatmap displaying the distribution of chromatin accessibility Log2FC values for genes within each of the six biological pathways (GO term analysis). Each column represents one pathway (gene count, n, indicated); each row represents an individual gene, ranked by Log2FC. Colour intensity reflects Log2FC magnitude according to the scale bar (right), with darker green indicating greater accessibility. Only peaks reaching statistical significance are shown (p < 0.05, Bonferroni correction). (G) Circular bar plots showing the number of differentially accessible peaks binned by Log2FC magnitude for each of the six biological pathways. Angular position indicates the Log2FC bin (0.5–1, 1–2, 2–3, 3–4, 4–5, 5–6), and bar length (radial extent) indicates the count of peaks in each bin. Shading intensity (light to dark green) corresponds to increasing Log2FC, as indicated in the shared legend (right). These plots highlight that the majority of peaks cluster in lower Log2FC bins, while a smaller subset shows high-magnitude accessibility changes.

After quality control, 3,351 L5 ET nuclei were retained (median of approximately 8,400 unique fragments per cell). snATAC-seq library quality was confirmed by TSS enrichment and fragment size distribution analysis (Additional File 2-Figure S3-A-B). Figure 3B shows the UMAP embedding of snATAC-seq data with L5 ET clusters highlighted (Additional File 2-Figure S3-C-D), and Figure 3C displays accessibility at L5 ET marker loci.

Comparison with age-matched uninjured cortex uncovered 5,482 regions that increased and 6,000 that decreased in accessibility (Figure 3C; Additional File 1-Table S3). Enhancers dominated the gains: 69.3% lay more than 2 kb from an annotated transcription start site. GO term analysis in semantic space confirmed enrichment for transcription, signal transduction, and chromatin remodeling as the largest clusters (Figure 3D; Additional File 2 – Figure S4A).

Analysis of the 5,482 gained accessibility peaks revealed broad but modest chromatin opening across growth-associated pathways. The majority of up-peaks (62%) fell within a log2FC of 1.5–2.5, with only 3.4% exceeding log2FC 3 (Figure 3E–G, Additional File 2 – Figure S4-B). Despite this limited amplitude, GO term analysis identified enrichment across regulation of cell growth (402 peaks), cytoskeleton organisation (306), nervous-system development (206), and axon guidance (157), with DNA-damage and repair contributing 43 peaks. Median fold-changes were highest in cell growth and cytoskeleton modules (log2FC ∼1.9) and lower in axon-guidance loci (log2FC ∼1.2) (Additional File 2 – Figure S4-B-D).

To contextualise the specificity of this chromatin response, we converted the 5,482 upregulated peaks to unique gene symbols, yielding 3,705 non-redundant genes, as multiple peaks can map to the same locus, and compared this gene set with those derived from published chromatin accessibility datasets from regeneration-competent peripheral neurons following sciatic nerve axotomy[35] and from retinal ganglion cells (RGCs) following optic nerve injury[19]. Across all comparisons, gene-level overlap was minimal: 9 genes shared with RGCs at 2 days post-injury (0.2%), 53 with sciatic nerve axotomy (1.3%), and 12 with dorsal column axotomy (0.3%) (Additional File 2 – Figure S21A–D), indicating that the corticospinal chromatin response to distal spinal injury is largely non-overlapping with both regeneration-competent PNS programmes and other CNS injury paradigms.

Thus, distal spinal injury re-opens regulatory DNA across major growth pathways, yet the depth of opening is low-to-moderate. Previous transcriptomic studies of distal spinal injury reported only negligible changes in growth-gene expression[6]. In contrast, our data suggest that the chromatin landscape does register the injury, albeit in a muted fashion.

### Lesion proximity amplifies chromatin reopening across all growth pathways

Lesion distance from the soma may limit the strength of injury-evoked growth programmes[6,19,22,31,36], so we compared the thoracic crush with a 350 micrometre intracortical micro-stab injury (ICI) that transects corticospinal axons within the motor cortex (Additional File 2-Figure S6-A-C). Nuclei harvested seven days after ICI were processed side-by-side with the thoracic cohort, wherein the thoracic cord is crushed and motor cortices are isolated 7 days later (Figure 4A; Additional File 1-Table S4). Representative histology demonstrating the thoracic crush injury and extent is shown in Additional File 2-Figure S5-A-B.

**Figure 4.**
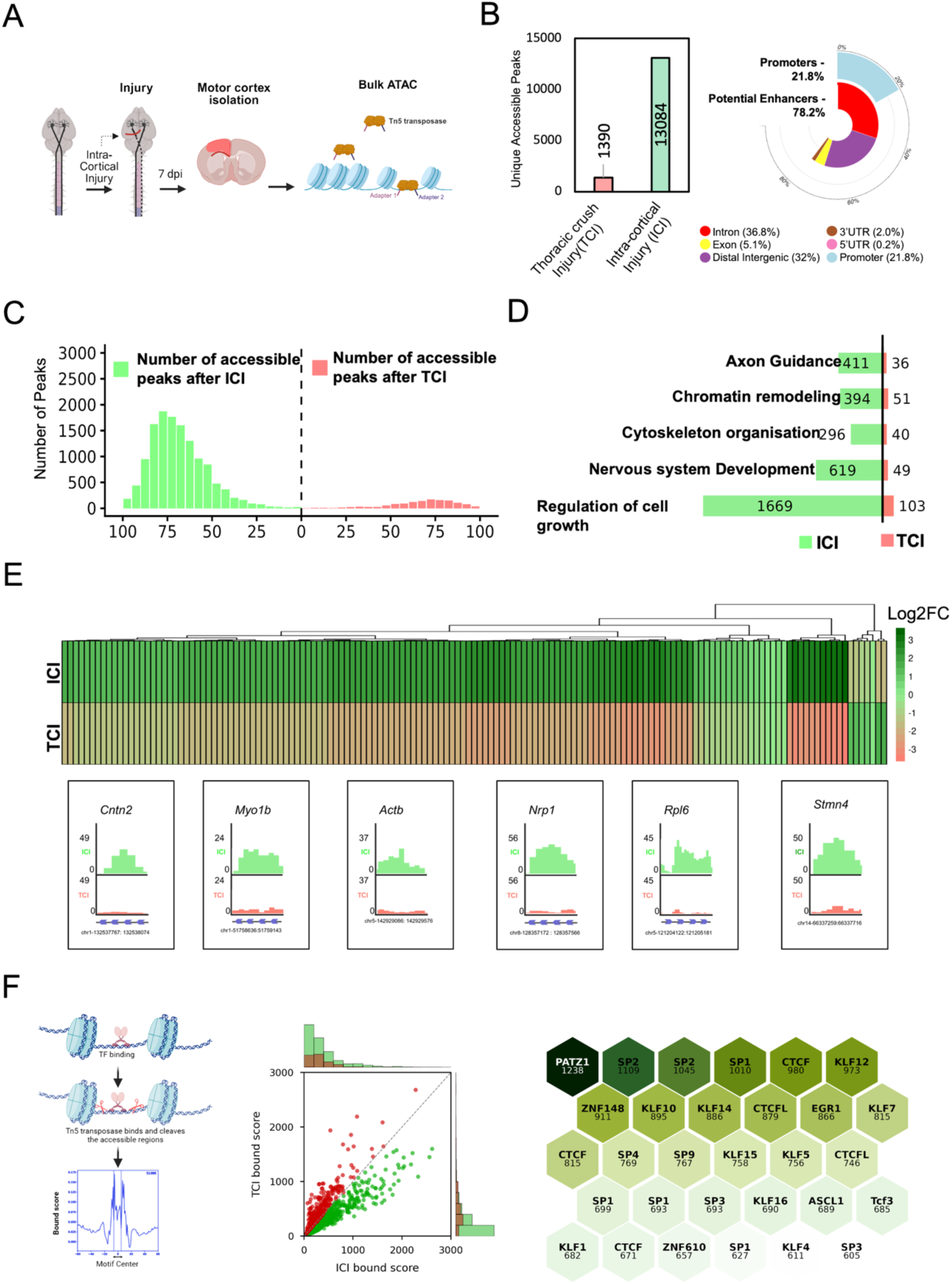
ATAC-seq reveals injury site proximity affects the re-opening of key loci approximately nine-to ten-fold. **(A)** Schematic of the bulk ATAC-seq experimental workflow for the intra-cortical injury (ICI) model. Intra-cortical injury was performed and motor cortices were isolated at 7 days post-injury (dpi). Chromatin was fragmented and tagged at accessible regions by Tn5 transposase with simultaneous adapter insertion (Adapter 1 and Adapter 2), followed by library preparation and sequencing. **(B)** *(Top left)* Bar graph comparing the total number of unique accessible chromatin peaks identified in Thoracic Crush Injury (TCI; n=1,390, red) versus Intra-cortical Injury (ICI; n=13,084, green), demonstrating an approximately six-fold greater chromatin opening response following ICI. *(Top right)* Donut chart showing the genomic distribution of the 13,084 ICI-specific accessible peaks: 78.2% fall within potential enhancer regions (comprising Intron 36.8%, Distal Intergenic 32%, Exon 5.1%, 3’ UTR 2.0%, 5’ UTR 0.2%) and 21.8% within promoter regions, indicating that the majority of injury-induced chromatin opening occurs at distal regulatory elements. **(C)**Mirror histogram comparing the distribution of accessible peaks by percentage fold change between ICI (green, left) and TCI (red, right). The x-axis represents percentage fold change (0–100%) and the y-axis shows the number of accessible peaks. ICI neurons display a substantially greater number of peaks across all fold-change values, with the distribution peaking around 75–80% fold change, while TCI neurons show far fewer peaks and a flatter distribution. **(D)** Diverging bar chart comparing the number of differentially accessible peaks in ICI (green) versus TCI (red/orange) across five GO-term-defined pro-growth biological pathways. Numbers indicate the peak count per condition for each pathway: Axon Guidance (ICI=411, TCI=36), Chromatin Remodeling (ICI=394, TCI=51), Cytoskeleton Organisation (ICI=296, TCI=40), Nervous System Development (ICI=619, TCI=49), and Regulation of Cell Growth (ICI=1,669, TCI=103). ICI consistently shows 8–16-fold more accessible peaks than TCI across all pathways. **(E)** *(Top)* Hierarchically clustered heatmap showing Log2FC of chromatin accessibility at pro-growth gene promoters, displayed as two rows corresponding to ICI (top) and TCI (bottom) conditions. Each column represents an individual pro-growth gene, clustered by accessibility profile. Colour scale indicates Log2FC (dark green = strongly increased accessibility, range 0 to +3; red/pink = decreased accessibility, range –3 to 0). ICI neurons show broadly elevated chromatin accessibility (predominantly green) across most pro-growth loci, while TCI neurons display markedly lower or negative accessibility (red) at the same loci. *(Bottom)* Representative UCSC Genome Browser tracks showing ATAC-seq read depth (y-axis) at six pro-growth gene loci in ICI (green) versus TCI (red): *Cntn2* (chr1:132537767–132538074), *Myo1b* (chr1:51758636–51759143), *Actb* (chr5:142929086–142929576), *Nrp1* (chr5:128357172–128357566), *Rpl6* (chr5:121204122–121205181), and *Stmn4* (chr14:66337259–66337716). In all six examples, ICI tracks show substantially higher accessibility peaks than TCI, consistent with the heatmap (p < 0.05). **(F)** Transcription factor (TF) footprinting analysis comparing TF occupancy at accessible chromatin regions between ICI and TCI conditions. *(Left)* Schematic illustrating the footprinting principle: TFs bound to DNA occlude Tn5 transposase cleavage, producing a characteristic depletion of reads at the motif centre flanked by elevated insertion signal — a “footprint” — whose depth reflects the degree of TF occupancy. The representative trace shows bound score (y-axis) plotted against distance from the motif centre (x-axis). *(Centre)* Scatter plot comparing TF bound scores calculated from ICI-accessible peaks (x-axis) versus TCI-accessible peaks (y-axis) for individual TF motifs. Each dot represents one TF; green dots indicate TFs with higher bound scores in ICI, and red dots indicate TFs with higher bound scores in TCI. Marginal histograms show the distribution of bound scores for each condition. The majority of TFs fall below the diagonal, indicating that TF occupancy is broadly greater in ICI than TCI. *(Right)* Hexbin plot displaying the top 30 TFs ranked by the magnitude of differential bound score (ICI bound score minus TCI bound score). Each hexagon represents one TF, with the numeric value indicating its differential score. Colour intensity (light to dark green) reflects the magnitude of the difference, with darker hexagons indicating greater ICI-biased TF occupancy. PATZ1 (score=1,238) shows the greatest differential occupancy between ICI and TCI, followed by SP2 (1,109 and 1,045), SP1 (1,010), CTCF (980), and KLF12 (973), among others.

Intra-cortical injury (ICI) resulted in 13,084 accessible unique peaks, while thoracic injury (TCI) resulted in 1,390 peaks only(Figure 4B; Additional File 1-Table S4). Fold-change values also shifted upward: gains after thoracic injury seldom exceeded a log2 change of 2.5, whereas ICI peaks were largely distributed between log2 3 and 4 (Figure 4C; Additional File 1-Table S4). Pathway tallies reflected this difference; injury-responsive axon-guidance elements rose from 36 to 411, chromatin-remodeling from 51 to 394, cytoskeleton organisation from 40 to 296, nervous-system development from 49 to 619, and regulation of cell growth from 103 to 1,669 (Figure 4D; Additional File 1-Table S4). Unbiased GO analysis of the top enriched terms across ICI and TCI confirmed a dramatic shift toward neuroregenerative pathways, including nervous system development, chromatin remodeling, and axon guidance, that was far more pronounced after ICI than TCI (p < 0.05; Additional File 2-Figure S6-A).

To confirm that this amplified response was selective for growth-associated loci rather than a global increase in chromatin accessibility, we examined fold enrichment at housekeeping genes. Non-growth control loci such as *Gapdh*, *Pten*, *Msh6*, and *Ctpb1* showed minimal difference in fold enrichment between ICI and TCI conditions (Additional File 2-Figure S6-C). By contrast, the pro-growth loci *Chchd3* and *Tubb4a* showed substantially greater accessibility in ICI compared to TCI (Additional File 2-Figure S6-D), consistent with the genome-wide patterns and confirming that ICI-driven chromatin opening is selective rather than genome-wide. This selectivity was further validated statistically: violin plots of sum accessibility signal at pro-growth gene loci showed a highly significant increase in ICI relative to TCI (Wilcoxon rank-sum test, p = 3.48×10⁻²¹, ***p < 0.001; Additional File 2-Figure S6-B).

Genome-browser views and heatmap analysis of pro-growth promoters further illustrated this pattern. The heatmap of ICI and TCI conditions showed broadly elevated accessibility at pro-growth loci in ICI (high log2FC, dark green) relative to the same loci in TCI (Figure 4E, top). Representative UCSC browser tracks at *Cntn2*, *Myo1b*, *Actb*, *Nrp1*, *Rpl6*, and *Stmn4* confirmed markedly higher accessibility peaks in ICI compared to TCI at all six loci (Figure 4E, bottom).

Since regeneration-associated pathways were broadly upregulated following ICI, we asked whether this chromatin opening was accompanied by concordant transcriptional changes. Side-by-side comparison of bulk ATAC-seq accessibility and snRNA-seq expression[6], following ICI revealed that chromatin opening at pro-growth loci was broadly mirrored by increased transcription (Additional File 2-Figure S8-B,C). A dot plot of selected pro-growth genes, including *Sox11*, *Patz1*, *Tubb2a*, and *Tubb2b*, showed that ATAC log2FC values were generally higher than RNA log2FC values at the same loci, consistent with chromatin opening preceding or exceeding transcriptional activation (Additional File 2-Figure S8-D).

Footprint analysis was consistent with these patterns: PATZ1, SP1/2/3, and several KLF motifs were prominent after ICI but weak or undetectable after thoracic crush (Figure 4F; Additional File 1-Table S4). To contextualise PATZ1 occupancy across injury conditions, we compared bound scores across three pairwise comparisons. ICI showed the greatest PATZ1 footprint score relative to both TCI (Δ≈0.10) and the uninjured state (Δ≈0.17), while the uninjured versus TCI comparison yielded a near-zero or slightly negative difference (Δ≈−0.01), indicating that PATZ1 binding is specifically elevated in the context of intracortical injury and is not simply a consequence of injury per se (Additional File 2-Figure S6-E).

Collectively, these data indicate that bringing the lesion closer to the soma markedly amplifies both the number of accessible regions and their fold-change across all growth-related pathways, with PATZ1 emerging as a transcription factor whose occupancy scales with this proximity effect.

### Developmental loss and injury-dependent resurgence nominate PATZ1 as a candidate chromatin regulator

Motivated by the observation that proximal injury restores extensive enhancer accessibility, we searched for transcription factors whose developmental decline and injury-dependent recovery might correlate with this difference. We analysed 616 vertebrate motifs with TOBIAS footprinting across the embryonic-to-adult ATAC-seq series and ranked them by footprint abundance, loss between P0 and adulthood, and gain after intracortical relative to thoracic injury (Additional File 2-Figure S9-A-B). To assess the genome-wide distribution of candidate factors, we first examined footprint counts across all accessible chromatin regions (Embryonic to Post-natal day 0 (P0)). PATZ1 occupied approximately 100,000 accessible regions genome-wide, placing it among the most broadly distributed transcription factors in the dataset (Additional File 2-Figure S9-C). Restricting this analysis to pro-growth regulatory elements further highlighted PATZ1 as enriched at growth-associated loci relative to other transcription factors (Additional File 2-Figure S9-D).

Intersecting the highest-scoring motifs with the growth-gene enhancer set highlighted PATZ1. At E18 and P0, PATZ1 footprints numbered 826 and 801, respectively; counts fell to 462 at P4, 119 at P7, and 27 in the adult cortex, representing a 97% reduction. After intracortical injury, 644 PATZ1 footprints reappeared, whereas thoracic crush restored only 88. Co-occupancy analysis showed that 72% of PATZ1 sites lay within 25 bp of SP1 or SP2 motifs and 61% within 50 bp of KLF7 or KLF4 motifs, a configuration associated with GC-rich stripe elements (Additional File 2-Figure S7-C). Stripe factors are transcription factors that bind to GC-rich sequences and are thought to help define regulatory domains by clustering at active enhancers and promoters; they often co-occupy sites and may act cooperatively to maintain open chromatin. The PATZ1-bound enhancers mapped to genes involved in axon extension, cytoskeletal regulation, chromatin organisation, DNA repair, and growth signalling, the same pathways that reopen most strongly after proximal injury (Additional File 2-Figure S9-D). These converging patterns nominated PATZ1 as a candidate regulator whose presence correlates with, and may contribute to, the more extensive chromatin access achieved when the lesion is close to the soma.

Immunohistochemical analysis confirmed that PATZ1 protein is expressed in postnatal day 0 (P0) mouse brain but substantially reduced in adult brain (Additional File 2-Figure S10-A-B). Quantification of PATZ1 fluorescence intensity revealed significantly higher expression in P0 samples compared to adult brain (Additional File 2-Figure S10-D), consistent with the developmental decline observed in footprinting analysis.

This developmental restriction was further supported by publicly available ENCODE bulk datasets, which showed high PATZ1 transcript levels and chromatin accessibility at the PATZ1 locus during embryonic stages, with both declining progressively toward adulthood (Additional File 2-Figure S20-C). A comparable developmental trajectory was observed in the peripheral nervous system, where PATZ1 mRNA expression in sciatic nerve[45] was highest at embryonic stages and declined progressively through postnatal timepoints (Additional File 2-Figure S20-D), suggesting that PATZ1 downregulation during maturation is a shared feature of both CNS and PNS development rather than a CNS-specific phenomenon. Consistent with this CNS-specific pattern, PATZ1 expression was largely absent across sensory ganglia datasets spanning multiple species[46] (Additional File 2-Figure S20-A), and while PATZ1 showed increased accessibility and transcription following peripheral nerve injury in dorsal root ganglion neurons[35], no comparable upregulation was observed following CNS dorsal column axotomy (Additional File 2-Figure S20-B), reinforcing the specificity of PATZ1 regulation to the CNS injury context studied here. Notably, snRNA-seq analysis following intracortical injury[6] confirmed that PATZ1 is endogenously upregulated at the transcriptional level in this condition (log2FC = 1.29; Additional File 2-Figure S8-H), providing direct evidence that the injury-dependent resurgence of PATZ1 footprints reflects, at least in part, transcriptional reactivation of the factor itself.

### Exogenous PATZ1 expression elevates chromatin accessibility after distal injury

Loss of PATZ1 footprints in the adult and their strong resurgence after intracortical injury suggested that reintroducing this factor might elevate the epigenetic response to a thoracic lesion toward the proximal-injury benchmark. To test this idea, we packaged murine PATZ1 under a CAG promoter in AAV9 with a P2A-GFP reporter (AAV9-CAG-PATZ1-P2A-GFP), enabling identification of transduced cells by GFP fluorescence. We injected the vector bilaterally into layer V motor cortex and performed a complete T8 crush two weeks later. A separate cohort received AAV9-CAG-GFP (control vector lacking PATZ1) followed by the same injury. Motor cortices were collected seven days after injury, and GFP-positive nuclei were isolated by FANS before generating 3’ snATAC-seq libraries (Figure 5A). We focused on the injured condition because our primary question was whether PATZ1 could enhance the injury response; comparison with the developmental data and with uninjured adult cortex (presented above) provides the relevant baselines.

**Figure 5.**
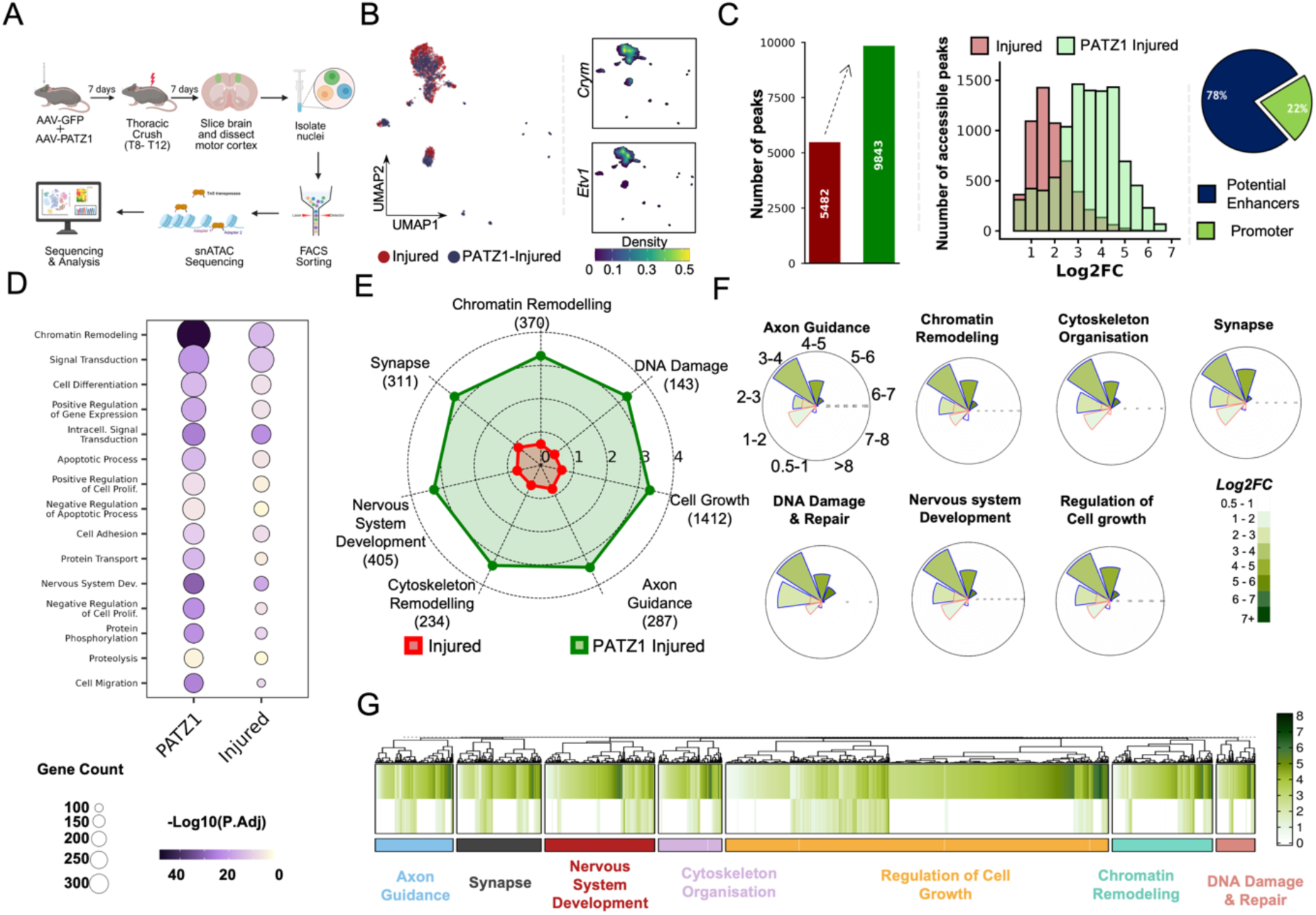
Single-nuclear (sn)ATAC-seq reveals that chromatin around pro-growth genes becomes more accessible upon PATZ1 over-expression. (A) Schematic of the snATAC-seq experimental workflow. Mice received dual AAV injections (AAV-GFP as control and AAV-PATZ1 for overexpression) followed by thoracic spinal cord crush injury. Motor cortices were dissected, nuclei isolated and FACS-sorted, and processed for snATAC-seq library preparation, sequencing, and bioinformatic analysis.(B) UMAP plot showing snATAC-seq nuclei from Injured control (red) and PATZ1-Injured (dark blue) conditions. Dot density is indicated by the colour scale (0–0.5). Feature activity for the L5 extratelencephalic (ET) corticospinal neuron markers *Crym* and *Etv1* (right panels) was used to identify and subset L5 ET neuronal clusters for downstream differential accessibility analysis. (C) (*Far left*) Bar graph comparing the total number of differentially accessible chromatin peaks in Injured control (red, n=5,482) versus PATZ1-Injured (green, n=9,843) neurons, demonstrating that PATZ1 overexpression substantially increases the number of accessible chromatin regions. (*Centre*) Overlaid histogram showing the distribution of accessible peaks by Log2FC magnitude in Injured control (red) and PATZ1-Injured (green) conditions (x-axis: Log2FC; y-axis: number of accessible peaks). PATZ1-Injured neurons show a notably greater number of peaks across all fold-change bins. (*Right*) Pie chart showing the genomic distribution of the additional peaks gained in PATZ1-Injured neurons: 78% located in potential enhancer regions (dark blue) and 22% in promoter regions (green). (D) Dot plot showing the top 15 enriched Gene Ontology (GO) biological process terms identified by unbiased analysis of differentially accessible peaks in PATZ1-Injured versus Injured control conditions. Each row is a GO term; columns represent PATZ1 and Injured conditions. Dot size reflects the number of genes associated with each GO term (scale: 100–300 genes), and dot colour indicates statistical significance expressed as –log10(adjusted p-value), with darker purple denoting greater significance. (E) Radar (spider) chart comparing mean chromatin accessibility Log2FC between PATZ1-Injured (green polygon) and Injured control (red polygon) neurons across seven GO-term-defined biological pathways: Chromatin Remodeling (n=370), DNA Damage (n=143), Cell Growth (n=1,412), Axon Guidance (n=287), Cytoskeleton Remodeling (n=234), Nervous System Development (n=405), and Synapse (n=311). Concentric circles represent Log2FC values from 0 to 4. The green polygon extends substantially beyond the red polygon across all axes, indicating genome-wide chromatin opening in PATZ1-Injured neurons. (F) Circular bar plots comparing the distribution of differentially accessible peaks by Log2FC magnitude between PATZ1-Injured (blue bars) and Injured control (red/orange bars) neurons for the same seven biological pathways. Angular position represents the Log2FC bin (0.5–1 through 7+), bar length indicates peak count within each bin, and shading intensity (light to dark green) reflects increasing Log2FC magnitude, as shown in the shared legend (right). Across all pathways, PATZ1-Injured neurons display greater peak counts and higher fold-change magnitudes than Injured controls. (G) Hierarchically clustered heatmap showing chromatin accessibility Log2FC values for individual genes across the seven biological pathways in PATZ1-Injured neurons. Columns represent individual genes (clustered by accessibility profile); rows are grouped by pathway, indicated by the colour-coded annotation bar beneath (Axon Guidance, Synapse, Nervous System Development, Cytoskeleton Organisation, Regulation of Cell Growth, Chromatin Remodeling, DNA Damage & Repair). Colour intensity (white to dark green) reflects Log2FC magnitude according to the scale bar (right, range 0–8). Only peaks meeting statistical significance thresholds are displayed (p < 0.05, Bonferroni correction, FDR < 0.05).

Representative immunofluorescence images confirmed successful AAV transduction in layer V motor cortex, with colocalization of PATZ1 and GFP signals verifying overexpression in transduced neurons (Additional File 2-Figure S11-A). Quantitative RT-qPCR demonstrated robust PATZ1 mRNA overexpression in AAV-PATZ1-injected animals compared to GFP controls (Additional File 2-Figure S11-B). To further validate that AAV-PATZ1-transduced neurons were captured in the transcriptomic dataset, we quantified the number of AAV-PATZ1-positive nuclei recovered from the snRNA-seq libraries; 2,301 PATZ1+ nuclei were identified compared to 1,255 GFP+ nuclei, confirming successful recovery of the transduced population for downstream analysis (Additional File 2-Figure S11-C).

snATAC-seq library quality for injured and PATZ1-injured samples was confirmed by TSS enrichment and fragment size distribution (Additional File 2-Figure S12-A-B). UMAP visualisation revealed distinct chromatin accessibility profiles between conditions, with CST and non-CST populations successfully identified; the CST fraction comprised 82% and 72% of nuclei in the injured and PATZ1-injured conditions, respectively (Additional File 2-Figure S12-C-D). Cell type-specific chromatin accessibility markers confirmed successful identification of neuronal populations (Additional File 2-Figure S12-E-F).

PATZ1 changed the distal-injury landscape both quantitatively and qualitatively. In PATZ1-positive nuclei, we detected 9,843 differential accessibility peaks relative to control GFP injured cortex (Figure 5B-C; Additional File 1-Table S6). The median log2 fold change for gained peaks doubled from 1.4 in GFP-control nuclei to 3.0 in PATZ1-expressing cells, and the high-amplitude tail expanded markedly: 7,022 peaks exceeded log2 4 after PATZ1, whereas only 71 did so in GFP controls. Consistent with PATZ1’s stripe-factor profile, 78% of new gains resided within distal enhancer annotations.

All core growth pathways showed increased accessibility with PATZ1, although the magnitude varied across categories. Axon-guidance peaks that gained accessibility rose from about 128 to 287; chromatin-remodeling peaks from roughly 72 to 214; cytoskeleton-organisation peaks from around 63 to 153; nervous-system-development peaks from about 59 to 266; and regulation-of-cell-growth peaks from approximately 112 to 524 (Figure 5D-E; Additional File 1-Table S6). Circular ridge-line plots showed a parallel shift toward higher fold-change values in every pathway, with the majority of peaks now falling between log2 2 and log2 4 (Figure 5F, Additional File 1-Table S6). To directly quantify the pathway-specific contribution of PATZ1, we calculated the difference in mean log2 fold change between PATZ1-injured and control GFP injured samples for each functional category; regulation of cell growth showed the largest differential effect, with PATZ1 increasing accessibility by an additional 4–5 log2FC units at 1,412 peaks, and no pathway showed a negative differential, indicating that PATZ1 universally enhances rather than suppresses injury-induced chromatin opening (Figure 5H; Additional File 1-Table S6). Alternative visualisations including grouped bar graphs, paired heatmaps, and diverging heatmaps of per-gene log2FC differences corroborated these pathway-level findings (Additional File 2-Figure S13-A-C).

Hierarchical clustering of the 9,843 PATZ1-induced accessibility gains revealed functional organisation of the chromatin response, with distinct blocks of co-regulated peaks distributed across all growth-related pathways (Figure 5G; Additional File 1-Table S6). The vertical banding pattern in the heatmap indicated that individual loci respond with varying sensitivity to PATZ1, ranging from modest (log2FC 1–2) to robust (log2FC > 4) increases in accessibility.

Together, these data indicate that forced PATZ1 expression substantially broadens the chromatin response to distal spinal injury, markedly increasing both the number of accessible regions and their fold change across all growth-related pathways, thereby converting a muted distal epigenomic response into one approaching the breadth, though not necessarily the full transcriptional output, of proximal cortical injury.

### PATZ1 couples enhancer reopening to transcriptional changes after distal injury

Because chromatin accessibility does not invariably translate into transcription, we next asked whether the expanded enhancer landscape produced by PATZ1 alters gene expression. Using matched samples from the same cohort analysed for snATAC-seq, we generated 3’ single-nucleus RNA-seq (snRNA-seq) libraries and retained 9,871 L5 ET nuclei after quality control (Figure 6A). UMAP visualization of snRNA-seq data showed cell clusters in injured and PATZ1-injured conditions (Figure 6B; Additional File 2-Figure S14-A-F). L5 ET neurons were identified by expression of marker genes including *Satb2*, *Crym*, and *Etv1* [42–44] (Figure 6B; Additional File 2-Figure S11-C-D). Differential expression analysis relative to GFP-injured controls (p-value < 0.05; |log2FC| ≥ 0.5) identified 1,825 upregulated and 4,369 downregulated genes (Figure 6C).

**Figure 6.**
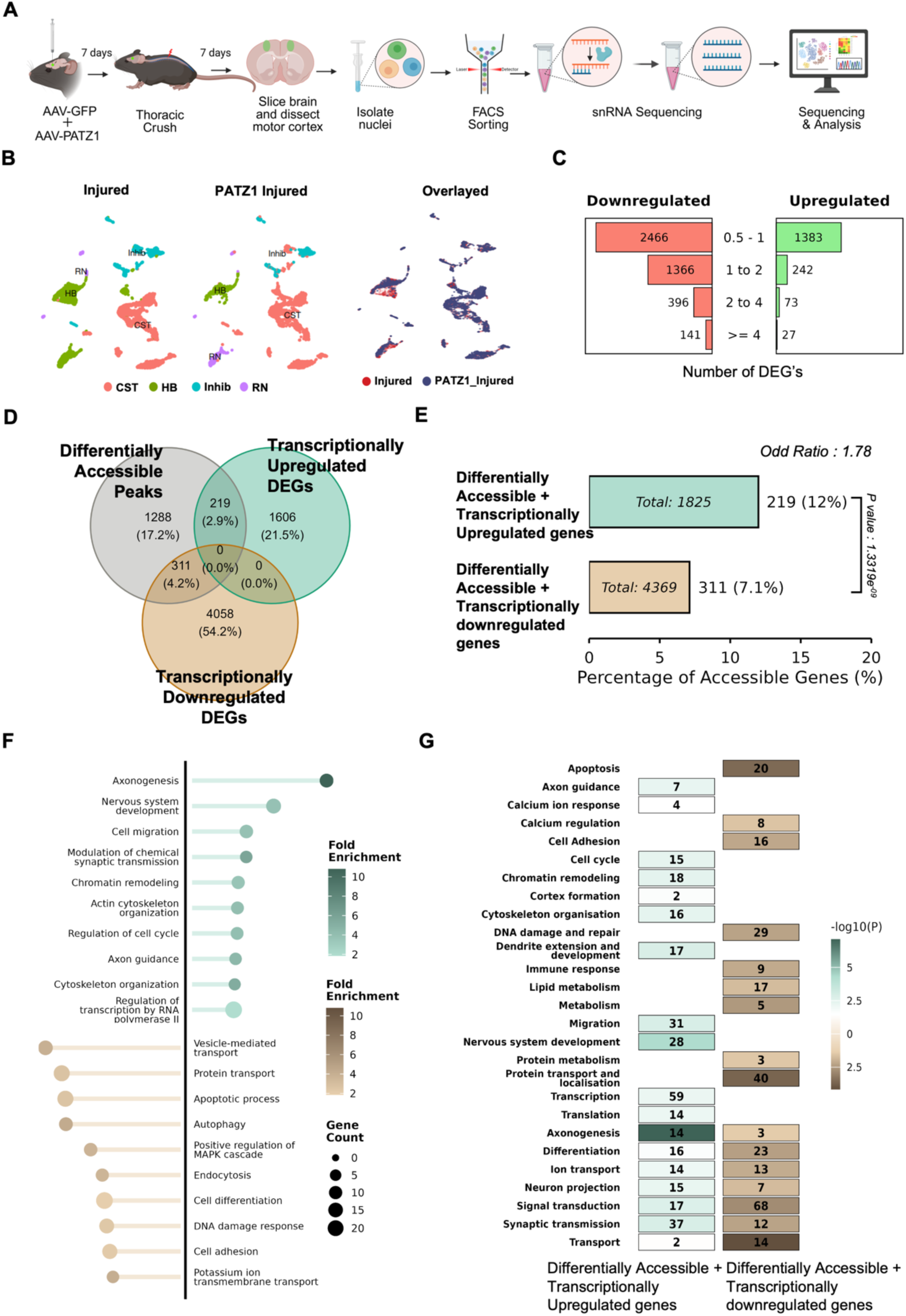
Integrative analysis of single-nucleus RNA sequencing and ATAC sequencing reveals transcriptional and chromatin accessibility profiles in PATZ1 tissue sample. (A) Schematic illustration of the experimental workflow. (B) UMAP visualisation of single-nucleus RNA-seq (snRNA-seq) data from the PATZ1 sample, with nuclei coloured by cluster identity. Abbreviations: CST, Corticospinal Tract neurons; HB, Hindbrain neurons; Inhib, Inhibitory neurons; RN, Red Nucleus neurons. (C) Distribution of differentially expressed genes (DEGs) by fold change categories. The horizontal bar plot displays the number of upregulated (green) and downregulated (red) genes across four log2 fold-change ranges (0.5 to 1, 1 to 2, 2 to 4, and ≥4), with counts indicated for each category. (D) Three-way Venn diagram showing the overlap between genes with increased chromatin accessibility (Differentially Accessible Peaks, grey circle) identified by snATAC-seq, transcriptionally upregulated DEGs (teal circle), and transcriptionally downregulated DEGs (orange circle) identified by snRNA-seq, in PATZ1-Injured versus Injured neurons. Numbers and percentages indicate the size of each intersection: Differentially accessible only — 1,288 (17.2%); Transcriptionally upregulated only – 1,606 (21.5%); Transcriptionally downregulated only — 4,058 (54.2%); Increased accessibility + transcriptionally upregulated — 219 (2.9%); Increased accessibility + transcriptionally downregulated – 311 (4.2%); Upregulated + downregulated overlap — 0 (0.0%); Triple overlap — 0 (0.0%). The 219-gene intersection of increased accessibility and transcriptional upregulation represents loci where PATZ1 overexpression drives coordinated chromatin opening and gene activation. (E) Bar chart quantifying the percentage of genes with increased chromatin accessibility (differentially accessible peaks) that also show transcriptional changes, comparing two gene sets. (Top, teal) Genes with increased chromatin accessibility that are also transcriptionally upregulated: 219 of 1,825 total (12%). (Bottom, orange) Genes with increased chromatin accessibility that are also transcriptionally downregulated: 311 of 4,369 total (7.1%). The x-axis shows the percentage of accessible genes (0–20%). The odds ratio of 1.78 indicates that transcriptional upregulation is 1.78-fold more likely to co-occur with increased chromatin accessibility than transcriptional downregulation, a statistically significant enrichment (p = 1.3319e⁻⁰⁸), supporting a functional link between PATZ1-driven chromatin opening and transcriptional activation. (F) Lollipop dot plot showing GO biological process enrichment for genes at the intersection of increased chromatin accessibility and transcriptional upregulation (top, teal/green bubbles) and at the intersection of increased chromatin accessibility and transcriptional downregulation (bottom, brown/tan bubbles). GO terms are listed on the y-axis. Bubble size reflects gene count associated with each term (scale: 0–20 genes). Bubble colour intensity indicates fold enrichment (scale: 2–10), with darker colour indicating greater enrichment. Upregulated accessible genes are significantly enriched for pro-regenerative terms including Axonogenesis, Nervous System Development, Cell Migration, Modulation of Chemical Synaptic Transmission, Chromatin Remodeling, Actin Cytoskeleton Organisation, Regulation of Cell Cycle, Axon Guidance, and Cytoskeleton Organisation. Downregulated accessible genes are enriched for terms including Vesicle-mediated Transport, Protein Transport, Apoptotic Process, Autophagy, Positive Regulation of MAPK Cascade, Endocytosis, Cell Differentiation, DNA Damage Response, Cell Adhesion, and Potassium Ion Transmembrane Transport. (G) Heatmap showing –log10(adjusted p-value) of parental GO term enrichment across two gene sets: genes with increased chromatin accessibility and transcriptional upregulation (left column) and genes with increased chromatin accessibility and transcriptional downregulation (right column). GO terms are listed on the y-axis; the number of genes contributing to each term is indicated within each cell. Colour intensity reflects –log10(p-value) (teal scale: 0–5 for upregulated; brown scale: 0–2.5 for downregulated), with darker colour indicating greater significance. Terms enriched in the upregulated accessible set include Protein Transport and Localisation (n=40), Transcription (n=59), Signal Transduction (n=68), Migration (n=31), Nervous System Development (n=28), DNA Damage and Repair (n=29), and Axonogenesis (n=14). Terms enriched in the downregulated accessible set include Apoptosis (n=20), Ion Transport (n=13), and Transport (n=14). The divergent enrichment profiles confirm that increased chromatin accessibility following PATZ1 overexpression is selectively associated with activation of pro-growth and pro-regenerative gene programmes, while a separate subset of accessible loci undergoes transcriptional downregulation enriched for stress-response and apoptotic pathways.

To characterise the relationship between PATZ1-induced chromatin accessibility and transcriptional activation, we performed integrative analysis of snATAC-seq and snRNA-seq data using a three-way Venn diagram (Figure 6D; Additional File 1-Table S7). Of genes with increased chromatin accessibility, 17.2% (1,288) showed accessibility changes alone without corresponding transcriptional changes, while 2.9% (219) exhibited both increased accessibility and transcriptional upregulation, and 4.2% (311) showed increased accessibility accompanied by transcriptional downregulation. Transcriptionally upregulated genes without detectable chromatin changes comprised 21.5% (1,606 genes), while transcriptionally downregulated genes without accessibility changes accounted for the largest fraction at 54.2% (4,058 genes).

To formally test whether transcriptional upregulation is preferentially associated with chromatin opening, we quantified the co-occurrence rates for each direction of transcriptional change (Figure 6E). Of 1,825 upregulated genes, 219 (12.0%) showed concordant increased accessibility, compared to 311 of 4,369 downregulated genes (7.1%). An odds ratio of 1.78 confirmed that transcriptional upregulation is significantly more likely to co-occur with PATZ1-driven chromatin opening than transcriptional downregulation (p = 1.33 × 10⁻⁸), establishing a functional link between PATZ1-mediated enhancer remodeling and gene activation at growth-relevant loci.

Unbiased Gene Ontology analysis of the 219 genes showing both increased accessibility and upregulated expression (top 10 terms, p < 0.05) revealed selective enrichment for pro-regenerative programmes (Figure 6F). Strikingly, the most significantly enriched terms included axonogenesis, axon guidance, nervous system development, cell migration, chromatin remodeling, actin cytoskeleton organisation, regulation of cell cycle, modulation of chemical synaptic transmission, and cytoskeleton organisation, a functional signature that directly maps onto the core requirements for axon regrowth. The genes contributing to the axonogenesis, nervous system development, and chromatin remodeling terms are displayed as a paired ATAC-RNA heatmap in Additional File 2 – Figure S15B (upper panel). This enrichment pattern demonstrates that PATZ1-driven chromatin opening is not random but is disproportionately concentrated at loci whose activation would be most relevant to regeneration. By contrast, the 311 genes showing increased accessibility alongside transcriptional downregulation were enriched for vesicle-mediated transport, protein transport, apoptotic process, autophagy, MAPK cascade regulation, endocytosis, cell differentiation, DNA damage response, cell adhesion, and potassium ion transmembrane transport (Figure 6F, lower panel; Additional File 2 – Figure S15B, lower panel).

A parallel heatmap of parental GO term enrichment across both gene sets confirmed this divergent pattern (Figure 6G): upregulated accessible genes were most significantly enriched for protein transport and localisation (n = 40), transcription (n = 59), signal transduction (n = 68), migration (n = 31), nervous system development (n = 28), DNA damage and repair (n = 29), and axonogenesis (n = 14), while downregulated accessible genes were enriched for apoptosis (n = 20), ion transport (n = 13), and general transport (n = 14).This bidirectional pattern suggests that PATZ1 does not uniformly activate growth programmes but selectively reshapes the transcriptional landscape, reactivating developmental gene programmes while partially suppressing mature neuronal identity genes.

Importantly, while the identity of the 219 concordantly upregulated genes is pro-regenerative, the magnitude of their transcriptional induction is modest. Examination of the flanking heatmaps of Additional File 2 – Figure S15A reveals that the vast majority of these genes fall into the low transcriptional output bin (log2FC 0.5–2), with very few reaching medium (log2FC 2–4) or high (log2FC ≥ 4) expression levels. Of the upregulated genes with detectable accessibility changes, 55 fall in the low accessibility/low transcription quadrant and 113 in the medium accessibility/low transcription quadrant, with only 44 genes reaching high chromatin opening and none achieving high transcriptional output. This pattern, in which the correct genes are opened but insufficiently to drive robust transcription, indicates that while PATZ1 establishes a broadly permissive chromatin landscape, the transcriptional machinery required to capitalise on this newly accessible chromatin is not yet fully engaged.

To assess whether this epigenomic broadening was accompanied by a transcriptional programme that genuinely approached that of proximal cortical injury, we compared snRNA-seq datasets from PATZ1-overexpressing injured cortex and intracortically injured (ICI) cortex[6]. UpSet plot analysis of differentially expressed genes (DEGs; p < 0.05, log2FC > 0.5) revealed that while both conditions engaged partially overlapping transcriptional programmes, the majority of DEGs were condition-specific: PATZ1 overexpression produced 1,528 uniquely upregulated and 3,639 uniquely downregulated genes, while ICI produced 1,233 and 819 unique up- and downregulated genes respectively (Additional File 2-Figure S8-G). The shared upregulated intersection comprised 189 genes co-induced by both conditions, representing a core injury-responsive pro-growth programme common to both paradigms. Scatter plot analysis of log2FC concordance across the 1,027 shared DEGs confirmed that 55.8% were regulated in the same direction by both conditions, indicating that PATZ1 overexpression partially but incompletely recapitulates the transcriptional landscape of proximal cortical injury (Additional File 2-Figure S8-H).

A separate question is whether PATZ1 engages the mTOR/PTEN pathway, which has independently been shown to promote axon regeneration following spinal cord injury[9,47]. To address whether PATZ1 operates through the related mechanism employed by *Pten* knockout (*Pten* KO), we compared the transcriptional programme induced by PATZ1 overexpression with that of *Pten* KO in injured retinal ganglion cells[8], a well-characterised pro-regenerative intervention that acts primarily through mTOR pathway activation. Of the 1,825 genes upregulated by PATZ1, only 57 (2.7%) overlapped with genes upregulated in the Pten knockout injury dataset (Additional File 2 – Figure S20-E). Moreover, scatter plot analysis of log2FC values for these 57 shared genes revealed no significant correlation between the two datasets (Pearson R = 0.079, p = 0.56), indicating that despite partial gene overlap, PATZ1 overexpression and *Pten* knockout drive largely distinct transcriptional programmes in injured neurons.

These findings suggest that PATZ1 promotes chromatin remodeling through a mechanism that is largely independent of mTOR signalling, and that its combination with mTOR-activating interventions could therefore produce additive rather than redundant effects on axon regeneration. Taken together, the transcriptomic comparisons above reveal that PATZ1 neither recapitulates the full transcriptional programme of proximal cortical injury nor engages the mTOR-dependent pathway activated by Pten deletion — the two best-characterised routes to CNS axon regeneration. This positions PATZ1 as operating through a distinct, chromatin-priming mechanism that prepares growth-associated loci for activation without itself supplying the transcriptional drive needed to complete that activation. In other words, PATZ1 opens the chromatin gate at the right genes, but the pro-growth transcription factors required to walk through that gate and convert accessibility into a functional regenerative programme are absent. The prediction that follows directly from this model is that PATZ1 alone, delivered without complementary transcriptional activators, should be insufficient to produce meaningful axon growth, a prediction we tested directly in a pyramidotomy model that assesses the axonal sprouting.

AAV-PATZ1 or AAV-GFP was injected into motor cortex, followed by unilateral pyramidotomy 60 days later, and corticospinal axon sprouting was quantified in coronal spinal cord sections at increasing distances from the lesion border (Additional File 2-Figure S18-A). CST labelling was confirmed with medullary pyramids, and pyramidotomy was verified by PKCγ immunohistochemistry (Additional File 2-Figure S18-B). Qualitative examination of GFP-labelled corticospinal axons suggested a trend toward increased sprouting in AAV-PATZ1-treated animals compared to AAV-GFP controls (Additional File 2-Figure S18-C). Quantification of fibre index at increasing distances from the pyramidotomy lesion (0-200 μm, 200-400 μm, 400-600 μm, and greater than 600 μm) revealed a trend toward increased sprouting at larger distances from the midline in PATZ1-treated animals, but this difference was not statistically significant (Additional File 2-Figure S18-D-E). Two-way ANOVA revealed a significant main effect of distance (F(3,9) = 9.689, P = 0.0035), indicating progressive decrease in fibre density with increasing distance from the lesion, but no significant main effect of PATZ1 (P = 0.4497) or interaction between PATZ1 and distance (P = 0.8538).

This null sprouting result is mechanistically consistent with the transcriptomic analysis above. PATZ1 opens chromatin at exactly the right genes, those annotated to axonogenesis, axon guidance, nervous system development, chromatin remodeling, and cytoskeleton organisation, yet their transcriptional output remains low. The adult cortex, even after PATZ1-mediated epigenomic remodeling following distal injury, lacks the full complement of pro-growth transcription factors required to translate open chromatin into meaningful transcription and axon growth. These findings establish chromatin accessibility as the rate-limiting epigenetic gate for regeneration-associated transcription: PATZ1 opens that gate at the correct loci, but pro-growth transcription factors are required to walk through it, making their combinatorial delivery the essential next step toward functional CNS repair.

### PATZ1 enhances enhancer acetylation and alters CTCF distribution after distal injury

To mechanistically dissect PATZ1-mediated epigenetic priming, we sought to determine whether the chromatin accessibility changes observed upon PATZ1 overexpression are accompanied by deposition of active enhancer marks and reorganisation of architectural boundaries. We performed CUT&RUN profiling for H3K27ac (a marker of active enhancers) and CTCF (an insulator protein that demarcates topological domains) in motor cortex tissue following thoracic crush injury (Figure 7A, Additional File 2-Figure S16-A).

**Figure 7.**
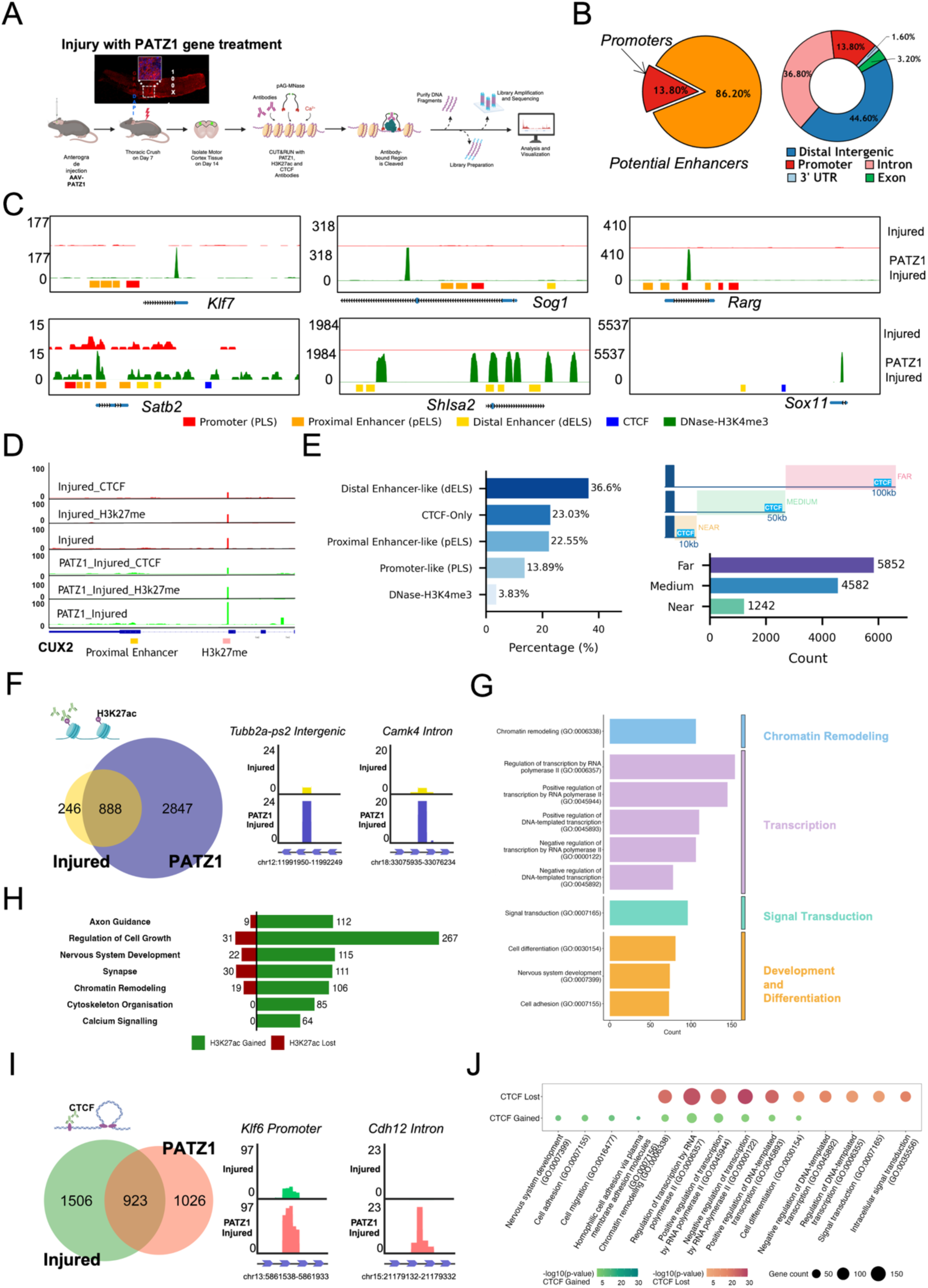
Genome-wide profiling of PATZ1 binding sites using CUT&RUN. (A) = Schematic of the CUT&RUN experimental workflow for genome-wide profiling of PATZ1 binding, CTCF occupancy, and H3K27ac marks. AAV-PATZ1 was injected anterogradely into the motor cortex; thoracic spinal cord crush was performed on Day 7; motor cortex tissue was isolated on Day 14. Chromatin was incubated with antibodies against PATZ1, H3K27ac, and CTCF, followed by pAG-MNase cleavage of antibody-bound regions, DNA purification, library preparation, and sequencing. (B) Genomic distribution of PATZ1-bound sites. (Left) Pie chart showing the broad classification of PATZ1 binding: 86.2% of sites fall within potential enhancer regions and 13.8% within promoters. (Right) Detailed donut chart showing the full genomic feature breakdown of PATZ1-bound sites: Distal Intergenic (44.60%), Intron (36.80%), Promoter (13.80%), Exon (3.20%), and other elements (1.60%). Together these indicate that the majority of PATZ1 binding occurs at distal regulatory elements rather than at gene promoters. (C) UCSC Genome Browser tracks showing CUT&RUN signal for Injured control (red) and PATZ1-Injured (green) at six representative pro-growth gene loci. Top row: *Klf7*, *Sog1*, and *Rarg*; bottom row: *Satb2*, *ShIsa2*, and *Sox11*. For each locus, the upper track shows Injured signal and the lower track shows PATZ1-Injured signal, with the y-axis indicating read depth. Regulatory element annotations are shown below each track: Promoter/PLS (red), Proximal Enhancer/pELS (orange), Distal Enhancer/dELS (yellow), CTCF binding sites (dark blue), and DNase-H3K4me3 active regulatory regions (dark green). PATZ1-Injured samples show increased signal at enhancer and promoter elements relative to Injured control at these pro-growth loci. (D) Genome Browser tracks at the Cux2 locus showing six CUT&RUN signal tracks: Injured_CTCF (red), Injured_H3K27ac (red), Injured PATZ1 (red), PATZ1_Injured_CTCF (green), PATZ1_Injured_H3K27ac (green), and PATZ1_Injured (green). Annotated regions below the tracks indicate a Proximal Enhancer (orange) and H3K27ac mark (pink). Increased H3K27ac and PATZ1 signal in the PATZ1-Injured condition at the proximal enhancer region of Cux2 illustrates PATZ1-dependent enhancer activation at a representative pro-growth locus. (E) (Left) Distribution of PATZ1-bound sites across cis-regulatory element categories: Distal Enhancer-like (dELS, 36.60%), CTCF-only (23.03%), Proximal Enhancer-like (pELS, 22.55%), Promoter-like (PLS, 13.89%), and DNase-H3K4me3 (3.83%). (Right) Bar graph showing the genomic distance of PATZ1-bound sites from the nearest CTCF binding site, categorised into three distance bins: Near (<10kb, n=1,242, teal), Medium (∼50kb, n=4,582, green), and Far (>100kb, n=5,852, purple). (F) (Left) Venn diagram showing the overlap of H3K27ac-enriched regions between Injured control (yellow, n=1,134 total: 246 unique + 888 shared) and PATZ1-Injured (purple, n=3,735 total: 2,847 unique + 888 shared) conditions. PATZ1 overexpression results in a substantial gain of H3K27ac-marked regions (2,847 unique to PATZ1-Injured), indicating widespread enhancer activation. (Right) Representative UCSC Genome Browser tracks showing H3K27ac signal (Injured in yellow; PATZ1-Injured in blue) at two loci: Tubb2a-ps2 Intergenic region (chr12:11991950–11992249) and Camk4 Intron (chr18:33075935–33076234), illustrating PATZ1-dependent gain of H3K27ac at these sites. (G) Horizontal bar chart showing GO biological process enrichment of H3K27ac regions uniquely gained in PATZ1-Injured samples (p < 0.05). Terms are grouped by functional category and colour-coded: Chromatin Remodeling (blue — Chromatin remodeling GO:0006338), Transcription (purple — Regulation of transcription by RNA Polymerase II, Positive regulation of transcription by RNA Polymerase II, Positive regulation of DNA-templated transcription, Negative regulation of transcription by RNA Polymerase II, Negative regulation of DNA-templated transcription), Signal Transduction (teal — Signal transduction GO:0007165), and Development and Differentiation (orange — Cell differentiation GO:0030154, Nervous system development GO:0007399, Cell adhesion GO:0007155). Bar length reflects gene count (x-axis, 0–150+). (H) Diverging bar chart showing the number of H3K27ac-marked regions gained (green) and lost (red) in PATZ1-Injured versus Injured control conditions across seven pro-regenerative biological pathways. All the GO terms enriched in Injured and PATZ1_injured are curated into these 7 regeneration-associated parent terms. Values shown: Axon Guidance (Gained=112, Lost=9), Regulation of Cell Growth (Gained=267, Lost=31), Nervous System Development (Gained=115, Lost=22), Synapse (Gained=111, Lost=30), Chromatin Remodeling (Gained=106, Lost=19), Cytoskeleton Organisation (Gained=85, Lost=0), and Calcium Signalling (Gained=64, Lost=0). PATZ1 overexpression results in a marked net gain of active enhancer marks across all pro-regenerative pathways, with the largest gain in Regulation of Cell Growth (n=267). (I) (Left) Venn diagram showing the overlap of CTCF binding sites between Injured control (green, n=2,429 total: 1,506 unique + 923 shared) and PATZ1-Injured (salmon, n=1,949 total: 1,026 unique + 923 shared) conditions. The reduction in unique CTCF sites in PATZ1-Injured relative to Injured (1,026 vs 1,506) suggests partial CTCF site reorganisation following PATZ1 overexpression. (Right) Representative UCSC Genome Browser tracks showing CTCF signal (Injured in green; PATZ1-Injured in red/salmon) at two loci: Klf6 Promoter (chr13:5861538–5861933) and Cdh12 Intron (chr15:21179132–21179332), illustrating condition-specific CTCF occupancy changes. (J) Bubble dot plot showing GO biological process enrichment for genes associated with CTCF-lost (red/salmon, top row) and CTCF-gained (green, bottom row) sites in PATZ1-Injured versus Injured conditions. GO terms are shown on the x-axis. Dot size represents gene count (scale: 50–150); colour intensity represents –log10(adjusted p-value) for CTCF Gained (green scale: 5–30) and CTCF Lost (red scale: 5–30). CTCF-lost sites are associated with terms including Nervous System Development, Cell Adhesion, Chromatin Remodeling, and Signal Transduction, while CTCF-gained sites show enrichment for overlapping transcription and differentiation pathways, suggesting dynamic CTCF reorganisation at functionally relevant loci following PATZ1 overexpression.

We first established a baseline by examining how injury alone affects these chromatin marks in the absence of PATZ1 overexpression. Comparing thoracic-injured cortex with uninjured controls, H3K27ac profiling revealed 483 regions unique to uninjured tissue, 690 overlapping regions, and 444 regions unique to injured tissue (Additional File 2-Figure S16-B). Representative genome browser tracks at loci such as *Fosb* promoter and *Cdh11* intron illustrate these dynamics (Additional File 2-Figure S16-B). Gene Ontology analysis showed that H3K27ac-lost regions were enriched for regulation of transcription by RNA polymerase II, nervous system development, and chromatin remodeling, while H3K27ac-gained regions were enriched for cell adhesion, intracellular signal transduction, extracellular matrix organisation, and neurogenesis (Additional File 2-Figure S16-C-D). The genomic distribution of differentially marked H3K27ac sites showed that most changes occurred at distal regions, with both gained and lost peaks distributed across distances from transcription start sites (Additional File 2-Figure S16-E).

CTCF profiling revealed a similarly balanced redistribution upon injury: 725 regions were unique to uninjured tissue, 1,008 were shared, and 1,421 were unique to injured tissue (Additional File 2-Figure S16-F). Representative tracks at *Rara* promoter and *Cdk20* intergenic regions demonstrate these boundary shifts (Additional File 2-Figure S16-F). Lost CTCF sites were enriched for negative regulation of DNA-templated transcription, regulation of DNA-templated transcription, and intracellular signal transduction, whereas gained CTCF sites were associated with negative regulation of apoptotic process, cell migration, and positive regulation of gene expression (Additional File 2-Figure S16-G-H). Together, these data confirm that distal injury triggers a balanced yet modest reshaping of enhancer marks and insulator boundaries, providing a baseline against which PATZ1-driven remodeling can be assessed.

Given this modest baseline response to injury, we next asked whether PATZ1 overexpression could drive more extensive chromatin remodeling. A widened ATAC footprint can reflect either direct binding by the introduced factor or secondary recruitment of chromatin modifiers, so we examined whether PATZ1 occupies the newly opened regions and whether its presence is accompanied by enhanced H3K27ac deposition or alterations in CTCF distribution. We generated CUT&RUN libraries from a separate cohort of animals that received either AAV-PATZ1 or AAV-GFP, followed by thoracic crush, and harvested on post-injury day 7.

Prior to profiling PATZ1-overexpressing tissue, we validated antibody specificity and AAV transduction efficiency. AAV-PATZ1-GFP targeting and transduction in layer V motor cortex neurons was confirmed by immunofluorescence (Additional File 2-Figure S17-A). CTCF antibody specificity was confirmed by appropriate nuclear localisation in both control and PATZ1 overexpression conditions, and H3K27ac antibody performance was verified by nuclear localisation consistent with active chromatin marks in both conditions (Additional File 2-Figure S17-B). Quantification of nuclear size and fluorescence intensity between AAV-GFP and AAV-PATZ1 conditions revealed no significant differences for either CTCF or H3K27ac staining, confirming that observed chromatin mark changes reflect PATZ1-dependent remodeling rather than technical artefacts (Additional File 2-Figure S17-B).FRiP scores and TSS enrichment profiles for all CUT&RUN libraries confirms consistent library quality and antibody specificity across uninjured, injured, and PATZ1-injured conditions for both CTCF and H3K27ac(Additional File 2-Figure S23-24).

CUT&RUN with a PATZ1 antibody identified 5,803 binding sites in PATZ1-overexpressing injured cortex (Figure 7B; Additional File 1-Table S9). Genomic distribution analysis revealed that 86.2% of these sites lay within regulatory regions classified as potential enhancers, with the remaining 13.8% at promoters (Figure 7B). Genome browser tracks at pro-growth genes including *Klf7*, *Satb3*, *Sog1*, *Rarg*, *ShIsa2* and *Sox11* illustrated PATZ1 binding at intergenic regions overlapping ENCODE-annotated regulatory elements and distal enhancers (Figure 7C). Additional examples at *Cux2* promoter regions demonstrated PATZ1 binding patterns, CTCF occupancy and H3K27ac marks in PATZ1 treated samples (Figure 7D). Classification of PATZ1-bound cis-regulatory elements showed that distal enhancer-like regions comprised the largest fraction (36.60%), followed by CTCF-only sites (23.03%), proximal enhancer-like regions (22.55%), promoter-like regions (13.89%), and DNase-H3K4me3 sites (3.83%). Motif analysis revealed enrichment of CTCF motifs at specific distances from PATZ1 binding sites (Figure 7E).

H3K27ac profiling in the same samples revealed that PATZ1 expression was associated with 2,847 new acetylation peaks compared to control GFP injured cortex, while only 246 were lost (Figure 7F; Additional File 1-Table S9). Notably, 94% of these de novo H3K27ac sites centred within plus or minus 250 bp of a PATZ1 summit, and 81% overlapped enhancers linked to genes upregulated in the snRNA-seq dataset. Browser tracks at *Tubb2a*-*ps2* and *Camk4*, illustrate the tight spatial coupling between PATZ1 occupancy and de novo H3K27ac signal (Figure 7F). Functional grouping of the acetylation gains highlighted transcriptional regulation, chromatin organisation, nervous-system development, and axon guidance, mirroring the categories that gained accessibility in the snATAC-seq dataset (Figure 7H; Additional File 1-Table S9).

We then profiled CTCF to assess whether PATZ1 influences domain boundaries. Relative to control GFP injured cortex, PATZ1-expressing tissue gained 1,026 new CTCF peaks and lost 1,506, leaving 923 shared sites (Figure 7I-J; Additional File 1-Table S9). Newly acquired CTCF binding occurred preferentially within 50 kb of genes involved in cell adhesion and axon guidance, whereas lost sites clustered near transcriptional repressors and differentiation inhibitors. Motif-distance analysis revealed an over-representation of CTCF motifs 120 to 180 kb away from PATZ1 summits, suggesting that enhancer activation by PATZ1 is accompanied by repositioning of distal insulator boundaries rather than wholesale redistribution across the genome (Figure 7E).

In summary, PATZ1 binds directly to thousands of distal enhancers opened after injury, and this binding is associated with H3K27ac deposition at those sites and selective reorganisation of CTCF boundaries around growth-related loci. These data provide biochemical evidence that PATZ1 converts accessible chromatin into functionally marked enhancer domains while modifying higher-order insulation to potentially facilitate long-range regulatory contacts.

### PATZ1 restructures higher-order chromatin topology to establish a growth-permissive three-dimensional landscape at growth loci

Our CUT&RUN data demonstrated that PATZ1 binds distal enhancers, deposits H3K27ac, and redistributes CTCF boundaries at growth-associated loci. These local chromatin changes raised the question of whether PATZ1 also reorganises higher-order genome architecture. To address this, we prepared Hi-C libraries from matched AAV-PATZ1 and AAV-GFP injured cortices harvested at 7 dpi, enabling simultaneous interrogation of chromatin compartments, topologically associating domains (TADs), and chromatin loops (Figure 8A).

**Figure 8:**
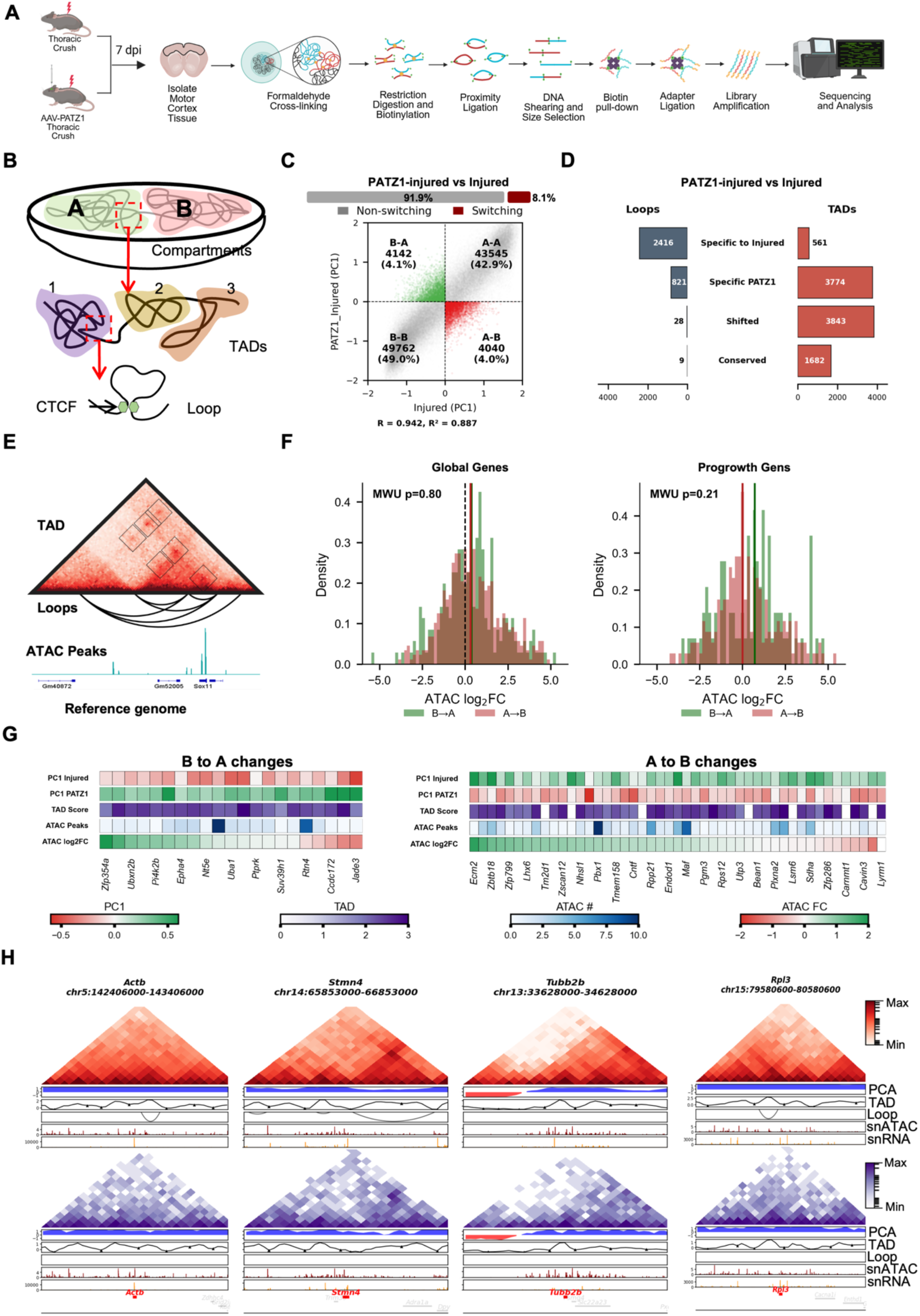
Functional analysis of compartment switching by PATZ1. (**A**) Experimental workflow for Hi-C analysis. (B)Schematic representation of hierarchical chromatin organisation. Illustration depicting the multi-level folding of the genome into A/B compartments (top), Topologically Associating Domains (TADs, middle), and chromatin loops (bottom). (C) Compartmental switching upon PATZ1 overexpression. (Top) Bar plot indicating the percentage of genomic bins maintaining their compartment status (non-switching, 91.9%) versus those undergoing transitions (switching, 8.1%) between Injured and PATZ1-treated Injured states. (Bottom) Scatter plot of PC1 values (Eigenvector 1) for Injured vs. PATZ1-treated conditions. Quadrants represent stable A-type (A-A, 42.9%), stable B-type (B-B, 49.0%), and switching compartments (B-A, 4.1%; A-B, 4.0%). R = 0.942 and R^2 = 0.887 values indicate Pearson correlation. (D) Reorganisation of loops and TADs. Diverging bar plots showing the number of chromatin loops (blue, left) and TADs (red, right) in four categories: Specific to Injured (Loops=2,416; TADs=561), Specific to PATZ1 (Loops=821; TADs=3,774), Shifted (Loops=28; TADs=3,843), and Conserved (Loops=9; TADs=1,682). PATZ1 overexpression is associated with a marked gain in TAD number and a reduction in unique loops. (E) Integration of 3D architecture and accessibility. Representative illustration showing the spatial overlap between TAD boundaries (triangular heatmaps), chromatin loops (arc plots), and ATAC-seq peaks (signal tracks) mapped to the reference genome. (F) Correlation between compartment switching and chromatin accessibility. Density plots showing the distribution of ATAC-seq Log_2(Fold Change) for genomic regions undergoing B-to-A (green) or A-to-B (salmon) transitions. (Left) Global genes; (Right) Pro-growth genes. Mann-Whitney U (MWU) test p-values are shown within each plot (Global genes: MWU p=0.80; Pro-growth genes: MWU p=0.21) (G) Multi-omic landscape of compartment transitions. Heatmaps showing coordinated changes in B-to-A (left) and A-to-B (right) switching regions. Tracks display PC1 values (Injured and PATZ1), TAD scores, ATAC peak counts, and ATAC-seq log_2FC for specific gene loci associated with these transitions. PC1 uses a red-to-green scale (–0.5 to +0.5), TAD score uses white-to-violet (0–3), ATAC peak number uses white-to-blue (0–10), and ATAC FC uses red-to-green (–2 to +2). (H) Genomic snapshots of pro-growth gene loci. Hi-C contact matrices and integrated tracks for *Actb*, *Stmn4*, *Rpl3*, and *Tubb2b* in Injured (red) and PATZ1-treated (purple) conditions. Tracks from top to bottom include: Hi-C heatmaps, PC1 Eigenvector, Insulation Score (Ins), chromatin loops, snATAC-seq accessibility, and snRNA-seq expression profiles.

Genome-wide principal component analysis of Hi-C contact maps revealed that PATZ1-expressing cortex was highly correlated with injured control (R = 0.942, R² = 0.887), indicating that the global compartment landscape is broadly preserved (Figure 8B, Additional File 1-Table S10). Nevertheless, 8.1% of genomic bins underwent compartment switching: 4,142 regions (4.1%) transitioned from inactive B to active A compartment, while 4,040 regions (4.0%) moved from A to B, with the remaining 91.9% of bins retaining their compartment identity.

At the sub-compartment level, TAD reorganisation was substantially more extensive than compartment switching (Figure 8C, Additional File 1-Table S10). While only 1,682 TADs were conserved between conditions, PATZ1 generated 3,774 unique TADs and shifted a further 3,843. By contrast, chromatin loop dynamics were far more restricted: PATZ1-expressing cortex contained only 821 unique loops compared to 2,416 loops specific to the injured control, with very few shifted (28) or conserved (9) loops, suggesting that PATZ1 preferentially reshapes domain boundaries rather than individual long-range contacts.

To understand the hierarchy of gene regulation operating within PATZ1-reorganised chromatin, we integrated compartment-switched regions with TAD boundaries and differentially accessible ATAC-seq peaks (Figure 8D). This multi-layer view illustrates how compartment identity, TAD structure, loop architecture, and local chromatin accessibility converge at individual growth-associated genes, providing a framework for interpreting the genome-wide patterns that follow.

To determine whether compartment switching was accompanied by preferential chromatin remodeling, we compared ATAC-seq log2 fold changes between B-to-A and A-to-B compartment-switched regions (Figure 8E, left). Genome-wide, the two distributions were indistinguishable (MWU p = 0.80), indicating that PATZ1 reshapes compartment identity broadly and without global chromatin bias. However, when the same analysis was restricted to pro-growth genes specifically, the B-to-A distribution showed a rightward shift relative to A-to-B, with the median accessibility gain in B-to-A regions exceeding that of A-to-B regions, though this trend did not reach statistical significance (MWU p = 0.21) (Figure 8E, right). This directional pattern, while not conclusive, suggests that PATZ1 may preferentially couple compartment activation with chromatin opening at growth-relevant loci, a selectivity that is diluted when all genomic regions are considered together.

To characterise the molecular features of individual genes within each compartment class, we integrated PC1 values from injured and PATZ1 conditions, TAD score, number of ATAC peaks, and ATAC log2FC for the top genes within B-to-A and A-to-B switched regions (Figure 8F). Genes undergoing B-to-A switching, including *Sox11*, *Klf6*, *Nt5e*, and *Syn1*, displayed negative PC1 values in the injured condition that shifted positive upon PATZ1 overexpression, confirming genuine compartment activation. Several of these loci additionally showed positive ATAC log2FC values and elevated ATAC peak counts, consistent with the directional trend observed in panel E. Conversely, genes within A-to-B switched regions, including *Myo1b*, *Sco1*, *Ecm2*, and *Maf*, showed the reciprocal PC1 trajectory and a more variable chromatin accessibility profile, with ATAC log2FC values distributed around zero, suggesting that compartment inactivation at these loci is not uniformly accompanied by chromatin closure.

Taken together, these data reveal that PATZ1 drives extensive three-dimensional genome reorganisation that operates primarily at the level of TAD boundaries rather than individual loops, with compartment switching affecting a selective but functionally relevant subset of genomic regions. Genome-wide, chromatin opening is architecturally agnostic, but a directional preference for chromatin gain in B-to-A-switched pro-growth loci suggests an emergent layer of regulatory specificity at the genes most relevant to axon regeneration. The absence of proportionate transcriptional activation despite this topological and chromatin priming is not a failure of PATZ1-mediated remodeling, but rather a precise mechanistic definition of the remaining barrier, the adult cortex following distal injury lacks the transcriptional complement needed to capitalise on an architecturally primed genome, positioning combinatorial delivery of PATZ1 with pro-growth transcription factors as the logical and necessary next step toward functional CNS repair.

## Discussion

During maturation, forebrain neurons undergo two major waves of chromatin restriction that progressively close promoters and enhancers associated with axon extension[13,14]. The second and more severe wave, occurring between P7 and adulthood, coincides with the known decline in corticospinal neuron regenerative capacity during the first two postnatal weeks[1,2]. Our multi-omic time-course suggests that this loss of accessibility is accompanied by staged withdrawal of GC-stripe factors, culminating in the near-complete disappearance of PATZ1 in adulthood[38]. Although our developmental bulk ATAC-seq was generated from heterogeneous forebrain tissue, these data provide a broad framework for understanding how CNS maturation imposes chromatin-level constraints on regenerative competence.

Lesion proximity strongly influenced the magnitude of this chromatin response. A distal thoracic lesion induced only modest reopening of growth-associated elements in L5 ET neurons, whereas an intracortical lesion near the soma reactivated an order of magnitude more sites, reinstating PATZ1 footprints and enhancer activity[6,19,22,31,36]. Because L5 ET neurons include corticospinal as well as other subcortical projection populations, these findings should be interpreted as reflecting an enriched projection-neuron population rather than purified corticospinal neurons[43]. Cross-injury comparisons further showed minimal overlap between the thoracic corticospinal response and injury-induced accessibility changes in retinal ganglion cells[19] or dorsal root ganglion neurons[35], with shared peaks comprising less than 2% in each comparison. This context-specificity argues that corticospinal neurons mount a distinct and limited epigenomic response to distal injury, rather than a generic neuronal injury programme. Together, these observations link chromatin plasticity to lesion proximity and nominate PATZ1 as a molecular correlate of this distance effect.

Reintroducing PATZ1 after thoracic injury converted the muted distal response into a chromatin state that partially resembled proximal injury. PATZ1 bound thousands of GC-rich enhancers, was associated with H3K27ac deposition, altered CTCF distribution and inter-TAD architecture, and produced selective B-to-A compartment shifts around pro-growth genes. These same regions gained accessibility, acquired active histone marks, and showed increased transcription, while immune and metabolic programmes were reduced. Thus, PATZ1 extends the stripe-factor paradigm from *in vitro* chromatin maintenance[38] to *in vivo* epigenetic remodeling in mature CNS neurons[20], although the causal sequence linking PATZ1 binding, enhancer acetylation, and higher-order chromatin reorganisation remains to be established.

A central question in CNS regeneration is why interventions that remodel the epigenome often fail to produce proportionate transcriptional or axonal responses[13,14,34]. Our data suggest that PATZ1 acts primarily as an architectural and epigenetic priming factor rather than as a complete pro-growth driver. Genome-wide, PATZ1 reshaped compartments and TADs without a global bias toward chromatin opening, as ATAC log2FC distributions were similar in B-to-A and A-to-B switched regions. However, at pro-growth loci, B-to-A-switched regions showed preferential chromatin opening, indicating that PATZ1 imposes regulatory selectivity specifically at growth-relevant genes. This selectivity is not apparent at the whole-genome scale but emerges when the analysis is focused on regeneration-associated loci.

Transcriptomic comparison between PATZ1-overexpressing thoracic injury and intracortical injury[6] further supports this model. Although many shared differentially expressed genes were regulated concordantly, only 189 genes were co-upregulated in both conditions, with most transcriptional changes remaining condition-specific. PATZ1 therefore moves the injured adult CNS epigenome toward a proximal-injury-like state but does not fully reproduce the proximal-injury transcriptional programme, likely because the pro-growth transcription factors normally induced by proximal injury are absent or insufficient after distal injury[4,20]. The concordantly upregulated genes may represent a functional core of the shared pro-growth programme and provide a rational target set for future combinatorial strategies.

This model is consistent with observations in regeneration-competent peripheral neurons, where three-dimensional genome reorganisation can precede transcriptional activation after injury[36]. PATZ1 appears to perform an analogous priming function in adult CNS neurons: it reorganizes the regulatory landscape and opens enhancers, but full regenerative output likely requires additional transcriptional engagement. Accordingly, PATZ1 overexpression alone did not produce statistically significant axonal sprouting in the pyramidotomy model[48]. This result supports the interpretation that chromatin opening is necessary but not sufficient for robust axon growth[13,34]. PATZ1 functions as an epigenetic enabler that lowers the barrier to gene activation, but additional pro-growth transcription factors are likely required to occupy newly accessible enhancers and drive the growth programme[4,7,20].

These findings refine current models of CNS regeneration in three ways. First, they separate the breadth of the injury response from its depth: distal axotomy engages growth-associated pathways, but at insufficient amplitude. Second, they place enhancer activation at the centre of regenerative amplification, with PATZ1 preferentially reopening enhancer-rich regulatory regions associated with growth competence. Third, they connect local enhancer priming to large-scale genome topology, showing that PATZ1 acts broadly on chromatin architecture while selectively promoting accessibility at pro-growth loci within that reorganized landscape.

The growth-associated categories reopened by PATZ1 span multiple barriers to regeneration. Axon guidance and cytoskeletal genes support structural remodeling[35]; nervous system development genes reflect partial reinstatement of juvenile growth states[6],consistent with transcriptomic profiles observed after proximal cortical axotomy; cell-growth and translational regulators provide the metabolic and signaling infrastructure for long-distance extension[20]; chromatin-remodeling genes may reinforce the permissive state; and DNA damage and repair genes may protect highly active enhancer-promoter contacts during transcriptional activation[36], a function supported by their enrichment at looped regulatory elements required for regeneration. The coordinated reopening of these categories suggests that PATZ1 does not act on a single pathway, but instead lowers several independent barriers to regrowth.

PATZ1 is well positioned to perform this role. It is a POZ/BTB and AT-hook zinc finger protein implicated in development, stem-cell maintenance, and gene regulation[39,49–51]. Its expression is high during early neurogenesis, declines toward adulthood, and remains associated with developmental plasticity. PATZ1 also belongs to the stripe transcription factor family, which binds GC-rich DNA, maintains open chromatin, recruits additional transcription factors, and stabilizes regulatory-element occupancy[38,52]. This prior biology aligns with our finding that PATZ1 binds growth-associated loci and promotes chromatin reorganisation in injured CNS neurons. The relative absence of PATZ1 in adult sensory ganglia[45,46], together with its injury-associated reactivation in peripheral but not central axotomy contexts[35], may partly explain why peripheral neurons mount a more robust epigenomic injury response than central projection neurons.

Prior studies also support a role for PATZ1 in acetylation and chromatin architecture. PATZ1 interacts with *NCoR* and *Sirt1*, both of which influence histone deacetylation,[39,53,54] and altered PATZ1 dosage during reprogramming affects H3K27ac and other permissive chromatin marks[39]. Our finding that PATZ1 overexpression drives H3K27ac deposition at growth-associated enhancers extends this regulatory role to injured cortical neurons. PATZ1 has also been described as an architectural transcription factor rather than a classical trans-activator[55,56], consistent with our Hi-C data showing changes in TAD boundaries and compartment identity. These data position PATZ1 as a multimodal chromatin regulator whose effects span enhancer acetylation, insulator redistribution, and large-scale topological reorganisation.

Mechanistically, PATZ1 may act through several cooperative layers: direct binding to GC-rich motifs[38], recruitment or redistribution of acetylation machinery[39,56], interaction with architectural proteins such as CTCF or cohesin,[22,36] and nucleosome eviction at selected regulatory elements[57]. Its selective action at growth-associated loci may reflect binding to GC-rich regions that retain residual accessibility or specific histone features in adult neurons. NucleoATAC analysis further supports the possibility that PATZ1 exposes regulatory elements by displacing nucleosomes, creating competent platforms for downstream transcription factor binding[57].

The major conceptual advance of this work is the identification of an epigenetic priming strategy for CNS regeneration. Much of the field has focused on delivering pro-growth transcription factors[4,5,7,8,20,58], but adult CNS neurons may fail to respond because key regulatory elements remain inaccessible[13,14,35]. PATZ1 provides a way to reopen growth-associated enhancers and render the adult epigenome more receptive to pro-growth signals. This distinguishes PATZ1 from classical pro-regenerative interventions such as PTEN knockout: only 57 genes, representing 2.7% of the PATZ1-upregulated programme, overlapped with PTEN knockout injured retinal ganglion cells[8], and their fold changes were not significantly correlated. PATZ1 therefore appears to engage a complementary programme, raising the possibility that chromatin priming and growth-pathway activation could be combined synergistically.

Several limitations should be considered. First, lesion distance and injury modality are not fully separable: intracortical injury involves local axon transection, whereas thoracic injury involves calibrated crush. Differences in inflammatory environment, Wallerian degeneration, retrograde signaling, and axon-end geometry may all contribute to the stronger proximal response[6]. Second, the thoracic model used crush rather than complete transection, which may influence local axon-stump interactions and retrograde signaling. Third, molecular analyses of thoracic and intracortical injury were performed at 7 dpi, whereas pyramidotomy sprouting was assessed at two months, reflecting the distinct biological logic and timescale of each model[48]. These differences should be considered when comparing effect sizes across paradigms.

Additional limitations relate to cell identity and assay design. The curated growth-associated gene list was supported by unbiased GO analysis, but alternative pathway definitions could influence interpretation. L5 ET neurons are enriched for corticospinal neurons but are not equivalent to a retrogradely purified corticospinal population[43]. Finally, chromatin and transcriptional datasets were generated in parallel rather than using single-cell multi-omic profiling from the same cells. Nevertheless, the convergence of accessibility, PATZ1 binding, active histone marks, chromatin architecture, and gene-expression changes at shared loci supports the overall model.

Future work should test whether PATZ1-mediated priming can be converted into long-distance axon growth and functional recovery when paired with pro-growth transcription factors such as KLF6, KLF7, SOX11, NR5A2,[4,7,20,58–60] or with mTOR activation[9,47] and guidance-cue scaffolds. Loss-of-function experiments in proximal injury would determine whether PATZ1 is required for the enhanced chromatin response to local axotomy. Defining PATZ1-interacting proteins through co-immunoprecipitation, mass spectrometry, BioID, or related proximity-labeling approaches will also clarify how PATZ1 coordinates enhancer acetylation, nucleosome displacement, and 3D genome remodeling in injured CNS neurons.

Taken together, these findings suggest that regenerative failure in the adult CNS has a substantial epigenetic component that can be therapeutically targeted[1–3]. PATZ1 does not appear to be a complete regeneration factor on its own. Instead, it broadens access to enhancer networks that feed into multiple pro-growth pathways, creating a permissive chromatin state that could be exploited by transcription factors, mTOR potentiators, scaffolds, or guidance cues[4,9,20,47]. This positions PATZ1-mediated epigenetic priming as a complementary strategy for spinal cord injury and other neurological contexts in which dormant growth programmes must be reactivated.

## Resource availability

All genomics datasets generated in this study have been deposited in the NCBI Gene Expression Omnibus (GEO) under accession number **PRJNA1217518**. A detailed summary of datasets and corresponding sample information is provided in **Additional File 1-Table 12**.

The metadata detailing the Supplementary Tables used for the results is provided in **Additional File 1-Table 13**. The code used for data processing and figure generation is available at https://github.com/VenkateshLab/Targeted-chromatin-remodeling-by-PATZ1.

Additionally, all datasets have been integrated into an interactive, user-friendly web interface accessible at https://patz1.netlify.app/ to facilitate data exploration.

## Supporting information

Addtional File 2 FIgure S 1 - S24

Addtional File 1 Table S1 to S 14

## Acknowledgements

We thank CSIR-CCMB for providing research facilities and support staff. We acknowledge Mrs. Zareena Begum (Tissue Culture), Mr. S. Prasanth and Mr. N. Sai Ram (Animal House), Dr. Karthik Baradwaj, Ms. Tulasi Nagabandi, and Dr. Md. Jafurulla (Genomics), Mr. G. Srinivas (FACS), and Mr. B. Suman and Dr. Nitla Venkata Mahesh (Microscopy) for their technical assistance. We also appreciate Fine BioChemical for the timely chemical supplies and Dr. Prakash Chermakani for reviewing spacing and errors in the final draft. We would also like to thank the funding agencies Council of Scientific and Industrial Research (CSIR), Department of Biotechnology (DBT), Science and Engineering Research Board (SERB), and BFI Biome, Government of India.

## Author contributions

Experimental design: A.S.M. and I.V. Anterograde injections, Intra-cortical surgeries, Thoracic crushes and pyramidotomies: A.S.M., N.K., D.S.B., S.S. Bulk ATAC sequencing: A.S.M., D.K.K. snATAC-seq: A.S.M., S.M. snRNA-seq: A.S.M., Y.S. Hi-C library preparation: D.S.B. Hi-C analysis: A.S.M., M.K.K. CUT&RUN library preparation: D.S.B., K.S. CUT&RUN analysis: D.S.B., M.K.K. Microscopy and imaging: A.S.M. Immunohistochemistry: A.S.M., D.K.K. Molecular cloning: Y.S. AAV production: Y.S., M.K.K, D.K.K. Sample processing and tissue sectioning: D.K.K. Neurite quantification: A.S.M., M.K.K. Data repository and pipeline generation for all genomics data: M.K.K. Manuscript writing: I.V. All authors read and approved the final manuscript.

## Supplemental information

**Document S1. Figures S1–S24,**

**Additional File 1-Table S1**: Excel file containing data that is used to generate Figure 1

**Additional File 1-Table S2**: Excel file containing data that is used to generate Figure 2

**Additional File 1-Table S3**: Excel file containing data that is used to generate Figure 3

**Additional File 1-Table S4**: Excel file containing data that is used to generate Figure 4

**Additional File 1-Table S6**: Excel file containing data that is used to generate Figure 5

**Additional File 1-Table S7**: Excel file containing data that is used to generate Figure 6

**Additional File 1-Table S9**: Excel file containing data that is used to generate Figure 7

**Additional File 1-Table S10**: Excel file containing data that is used to generate Figure 8

**Additional File 1-Table S5**: Excel file containing data that is used to generate Supplementary Figure S7

**Additional File 1-Table S8**: Excel file containing data that is used to generate Supplementary Figure S10

**Additional File 1-Table S11**: Excel file containing data that is used to generate Supplementary Figure S15

**Additional File 1-Table S12**: A detailed summary of datasets and corresponding sample information

**Additional File 1-Table S13**: The metadata detailing the Supplementary Tables used for the results

**Additional File 1-Table S14**: The metadata detailing the Supplementary Tables used for the Supplementary Figures

## Materials and Methods

### Animal Husbandry

All experimental protocols involving animals were approved by the Institutional Animal Ethics Committee (IAEC) of the Centre for Cellular and Molecular Biology (CCMB), Hyderabad, India (IAEC 14/2023, 29/2023, 51/2023, 10/2024, 53/2024). Mice (C57BL/6J strain) were bred and maintained under a 24-hour light-dark cycle with 12 hours of light and 12 hours of darkness. The ambient temperature was maintained between 23-25°C with humidity levels between 40% and 60%.

### Cloning and Virus Production

Constructs for candidate genes were purchased from Horizon Discovery: pCMV-SPORT6-PATZ1 (Clone ID: 4167023) from the MGC Fully Sequenced Mouse cDNA (pCMV) transcription factors (Catalog ID: MMM5173, Expression Ready MGC cDNA Libraries), while pAAV-CAG-tdTomato (codon diversified) and pAAV-CAG-GFP were purchased from Addgene (Plasmid catalogue ID# 59462, #37825). For viral production, the ORF of each gene was cloned into an AAV-CAG backbone (Addgene Plasmid #59462) using a standard cut-and-paste method. Maxipreps were prepared (Qiagen Endo-free kits, #1236) and fully sequenced. AAV9-tdTomato (Addgene Plasmid #59462) was produced in-house, achieving a titer value of 10×10^11 to 10^13 before injection. For experiments, viruses were mixed at a 1:3 ratio of pAAV-CAG-GFP to pAAV-CAG-PATZ1.

### Animal Surgeries Anterograde Injections

Following deep anaesthesia with ketamine (100mg/ml)/Xylazine (20mg/ml), mice were prepared by shaving and disinfecting the scalp and neck. They were positioned in a stereotaxic frame, and a 1 cm midline incision was made in the scalp. The skin was retracted using a Bulldog clamp, and a thin layer of the skull was gently peeled away. Non-replicating viral particles were injected using a Hamilton syringe (Sigma Aldrich, #20972) at four bilateral points relative to Bregma, with coordinates (x = −2, +2; y = −0.5, +0.5, z= 0.8, 0.6), delivering 1 µl of AAV (0.2 µl/2 min) to ensure widespread diffusion. After withdrawing the pipette, the incision was sutured, and appropriate postoperative care was administered.

### Thoracic Crush

Mice (n=3) post seven days of injection, were anaesthetised with ketamine (100mg/ml)/Xylazine(20mg/ml) and prepared for surgery by shaving and disinfecting the upper back and neck. A midline incision was made from the ears to approximately 2 cm down the neck and back. Fatty tissue was displaced laterally, and a bulldog clamp was used to retract skin and maintain exposure of the surgical site. The underlying paraspinal musculature was carefully dissected to expose the spinal column. The spinous processes at the T10-T12 level were identified and trimmed, if necessary, to facilitate access. The mouse was then mounted in a custom spine stabiliser to ensure proper immobilisation during the procedure. A laminectomy was performed at the T10-T12 level using fine forceps to carefully remove the dorsal lamina and expose the spinal cord. The exposed spinal cord was then crushed with calibrated blunt forceps for a precise duration of 10 seconds, applying consistent pressure across the full width of the cord. Following a crush injury, the vertebral column was repositioned, the surrounding tissues were returned to their anatomical positions, and the skin was sutured with black nylon thread. Post-operative care included analgesics, antibiotics, manual bladder expression twice daily until function returned, and maintenance on a heating pad during recovery. Visible hindlimb paralysis was consistently observed post-surgery, confirming successful injury. All procedures were performed in a type II safety hood under aseptic conditions.

### Intra-cortical Surgery

This procedure was modified from Wang *et al*., 2025 [6]. After deep anaesthesia with ketamine (100mg/ml)/Xylazine (20mg/ml), mice (n=3) were prepared by shaving the scalp and neck and disinfecting with betadine. Mice were positioned in a stereotaxic frame, and a 1 cm midline incision was made in the scalp. The skin was retracted with a Bulldog clamp, and the skull over the injection area was gently peeled away. Stereotaxic coordinates (x = −2, y = −0.5, y = +0.5, z = 0.6) were used to guide the procedure at specified points. A 30½ gauge needle bent into an L-shape was inserted 0.6 mm from the brain surface and turned 90 degrees to sever axons. The needle was retracted, the skin was sutured, and post-operative care was provided.

### Pyramidotomy

The procedure was adapted from [48]. Seven days post-injection, deep anaesthesia with ketamine (100mg/ml)/Xylazine (20mg/ml) was administered to mice (n=3 for control and n=4 for PATZ1 treated animals). A ventral incision was made near the oesophageal region, and after displacing the tissue layer beneath with blunt forceps, the medullary pyramid was exposed. The dura was cut, and the right pyramidal tract was incised using iridectomy scissors (0.5 mm width, 0.25 mm depth). Post-surgery, the oesophagus, trachea, and muscles were repositioned, and the skin was sutured.

### Axon-growth quantification for Pyramidotomy cord slices

Tissue preparation and imaging: At 8 weeks post-pyramidotomy, animals were transcardially perfused and spinal cords were processed as described below. For each animal, six transverse sections (50 µm thickness) spanning the C2-C4 spinal cord segments (1.5 mm spacing between sections) were selected for analysis. Sections were imaged using a Leica STED/confocal microscope with a 63× objective (tile scan in 520 × 520 µm format, 0.5-1 µm z-steps, scan speed 400 Hz; detector gain 1,250 V, laser power 40%). Maximum intensity projections were generated from the z-stacks for quantification.

### Fibre index calculation

Axonal regeneration was quantified by calculating a Fiber Index (FI) based on the number of GFP-positive axons crossing predefined virtual distance bins distal to the lesion. For each spinal cord section, the midline border was first identified anatomically using the spinal canal as a reference. A reference line was then drawn through the central canal at the lesion epicenter (0 µm) to establish the point of origin for subsequent measurements. Automated quantification was performed using DeNAT, a machine-learning–based tool for neurite detection and spatial measurement [61]. After uploading each GFP-labeled spinal cord image, users manually defined (i) the midline and (ii) the region of interest (ROI) corresponding to the injured and regenerating side. Based on the image’s spatial calibration and pixel-to-micron ratio, DeNAT automatically generated four parallel virtual lines caudal to the lesion border at the following distances: 0–200 µm, 200–400 µm, 400–600 µm and >600 µm. The integrated ML model then detected GFP-positive fibers and quantified the number of axons crossing each virtual line. The Fiber Index for each distance bin was calculated as the number of GFP-positive axons was normalized based on each bins.

To normalize for variability in viral transduction efficiency, GFP-positive neurons in the medullary pyramids were quantified. Coronal sections (50 µm) through the medullary pyramids were imaged (3-4 slices per) animal) using a Zeiss ApoTome microscope at 10× magnification. GFP-positive cell bodies were counted from 3-4 sections per animal using ImageJ software [62]. The fiber index for each distance bin was calculated by dividing the average number of axons crossing each distance line by the normalized pyramid cell count, providing a measure of axonal density that accounts for differences in initial labeling efficiency between animals.

### Statistical analysis

Fiber index data were analysed using two-way ANOVA between control and treated fibre indices(AAV-GFP vs. AAV-PATZ1) and distance from the lesion (0-200 µm, 200-400 µm, 400-600 µm, >600 µm) as fixed factors, and subject as a random effect to account for repeated measurements across distances within individual animals. This measures both main effects and the treatment × distance interaction term. Post-hoc pairwise comparisons between treatment groups at each distance were performed using Sidak’s multiple comparisons test to control for family-wise error rate. Data are presented as mean ± SEM with n = 3-4 animals per group. Statistical significance was set at α = 0.05. All statistical analyses were performed using GraphPad Prism version 10.5.0 (GraphPad Prism Software, Boston, Massachusetts USA, www.graphpad.com)

### Immunohistochemistry

For spinal injury experiments, adult animals were decapitated following anaesthesia, and the brain and spinal cord were extracted and post-fixed overnight in 4% PFA (Sigma Aldrich, #158127) at 4°C. Spinal cord tissue was embedded in 12% gelatin (Sigma Aldrich, G2500) in PBS, and 50 µm sagittal sections were obtained using a vibratome. Sections were permeabilised in PBST, blocked with 5% normal goat serum, and incubated overnight at 4°C with the following primary antibodies: anti-GFAP (Cell Signalling Technology, mAb #3670, 1:500) to assess injury depth; PKCγ (GeneTex, GTX107639, 1:200) to confirm unilateral pyramidotomy; CTCF (Cell Signalling Technology, #3418, 1:500) and H3K27ac (Abcam, ab4729, 1:500) for antibody validation; and PATZ1 (MyBioSource, MSB9403528, 1:100) to confirm PATZ1 overexpression. After three PBS washes, sections were incubated with Alexa Fluor-conjugated secondary antibodies (Thermo Fisher, R37119, 1:500) for 2 hours at room temperature. Following counterstaining with DAPI (F6057-Sigma Aldrich), sections were mounted and imaged using a confocal microscope for high-resolution analysis.

Intensity and nuclear size Calculation

### Fluorescence Intensity and Nuclear Size Quantification

Fluorescence intensity of CTCF and H3K27ac immunolabeling was quantified from confocal images using ImageJ software[62] (NIH). For each marker, only the red channel was used for intensity measurements, corresponding to the Alexa Fluor-conjugated secondary antibody signal for CTCF (Cell Signalling Technology, #3418) and H3K27ac (Abcam, ab4729) respectively. For each image, a region of interest (ROI) was drawn around individual immunopositive cells and mean fluorescence intensity was measured. Background fluorescence was subtracted by measuring signal from an adjacent region devoid of cellular staining. Measurements were performed across both experimental conditions: AAV-GFP (control) and AAV-GFP + AAV-PATZ1 (overexpression, 1:3 ratio).

Nuclear size was quantified from the DAPI channel of the same confocal images using ImageJ software. DAPI-stained nuclei were thresholded to create binary masks, and the cross-sectional area of individual nuclei was measured using the “Analyze Particles” function. Only nuclei meeting minimum size and circularity criteria were included to exclude debris and non-nuclear signals. Nuclear size measurements were obtained from both control and PATZ1 overexpression conditions to confirm that any observed differences in CTCF or H3K27ac signal were not attributable to changes in nuclear morphology.

Statistical comparisons of fluorescence intensity and nuclear size between AAV-GFP and AAV-PATZ1 conditions were performed using a two-tailed non-parametric Mann–Whitney U test, with significance set at p < 0.05.

### Nuclear extraction and Fluorescence-activated nuclei sorting (FANS)

Adult mice (n=3 per experimental group) received anterograde injections of viral vectors for Control and PATZ1 treatment. One week later, animals were challenged with thoracic crush injuries, as described above. One week post-injury, animals were sacrificed, and the motor cortices were dissected. Tissue was collected only from the motor cortex region, which shows green fluorescence under UV light when viewed through a yellow filter. The dissected motor cortex tissues were added to pre-chilled 2 ml EZ lysis buffer (Sigma, NUC-101) and dounced (Sigma, D8938) slowly and carefully (Dounce A, 40 times; Dounce B, 45 times), then incubated for 5 minutes at room temperature. Post incubation, the content was transferred to a 15ml falcon tube and centrifuged at 500g with acceleration and deceleration set to 2. The supernatant was removed, and the pellet was dissolved in Nuclear Suspension Buffer (freshly prepared: 5 ml PBS, 25 uL BSA (20 mg/ml) and 25 uL RNase inhibitor (40 U/ul) (Invitrogen, 10777019)) and strained through a Polystyrene tube with cell strainer (Falcon, 352235). Dissociated nuclei were flow-sorted on a BD FACS using an appropriate gating strategy (Sort type: Purity). Nuclei were first gated by side scatter height (SSC-H) vs. forward scatter height (FSC-H) and side scatter area (SSC-A) vs. FSC-A to separate debris from intact nuclei. Nuclei that passed these filters were further selected by SSC-W vs. SSC-H to eliminate potential doublets. Finally, nuclei were gated by fluorescence marker intensity to collect only the brightest approximately 5000 nuclei for ATAC sequencing. Note that in this nuclear isolation protocol, DAPI was used as a positive marker to identify intact nuclei rather than as a viability dye, where DAPI-positive selection enriches for intact, well-preserved nuclei, which is the appropriate gating strategy for single-nucleus genomics applications.

### Single-Nucleus ATAC Sequencing Library construction (sn-ATAC)

Single-cell ATAC-seq datasets were generated using the Chromium Single Cell ATAC kit from 10x Genomics, following the manufacturer’s instructions (10x Genomics, CG000496 Chromium Next GEM Single Cell ATAC Reagent Kit v2). Nuclei were isolated from the motor cortex of mice (n=3 per experimental group). The isolated nuclei suspensions (3,000-5,000 nuclei per sample) were added to the transposition mix containing Tn5 transposase enzyme, which simultaneously fragments DNA and inserts adapter sequences at accessible chromatin regions. Gel beads in emulsion (GEMs) were generated on the Chromium Next GEM Chip H by combining barcoded gel beads with the transposed nuclei suspension and partitioning oil. After GEM generation, the barcoded DNA fragments were recovered and purified using streptavidin beads, followed by size selection with SPRIselect beads (Beckman Coulter, B23318) and a series of 80% ethanol washes. The purified DNA fragments were eluted in elution buffer (10 mM Tris-HCl pH 8.0), followed by indexing PCR amplification (12 cycles) to add sample indices and Illumina sequencing adapters. Final library cleanup was performed using an additional round of size selection with SPRIselect beads. Library quality was assessed using a Bioanalyzer High Sensitivity DNA Kit (Agilent, 5067-4626) to confirm appropriate fragment size distribution (200-800 bp). Sequencing was performed on an Illumina NovaSeq 6000 instrument using a paired-end configuration with read length of 2X150 bp, approximately 300 million paired-end reads per sample to achieve sufficient coverage for downstream analysis.

### Bulk Assay for Transposase-Accessible Chromatin Sequencing (Bulk ATAC)

Motor cortices were isolated from mouse pups at P0, P4, P7, and from adult mice (n=3 per experimental group) that underwent intra-cortical and thoracic crush surgery. Tissue samples (20-30 mg) were minced and placed into ice-cold PBS. Nuclei extraction was performed following the protocol described above. Nuclei were counted, and approximately 50,000 to 100,000 nuclei per sample were used for the assay. The nuclear pellet was resuspended in Tagmentation Master Mix containing transposase enzyme (Active Motif, #53150), which inserts sequencing adapters into accessible regions of chromatin according to the manufacturer’s protocol. Tagmentation was carried out by incubating the nuclei with the Tagmentation Master Mix at 37°C for 30 minutes with gentle agitation. Following tagmentation, the DNA was purified using a DNA purification column (Active Motif, included in kit). The purified DNA was eluted in 10 mM Tris-HCl (pH 8.0) and subjected to PCR amplification (12-15 cycles) using indexed primers compatible with Illumina sequencing. Post-PCR, a final purification step was performed using SPRI beads (Beckman Coulter, B23318) at a 1.8× ratio to remove primer dimers and unincorporated primers. Libraries were quantified using Qubit dsDNA HS Assay Kit (Thermo Fisher, Q32851), and DNA size distribution was assessed with a TapeStation 4200 (Agilent) using a D1000 ScreenTape (Agilent, 5067-5582) before pooling and loading the libraries onto the Illumina NovaSeq 6000 sequencing platform for ∼30 million paired-end reads per library.

### High-throughput Chromosome Confirmation Capture (Hi-C)

Mice (n=3 to 4 per experimental group) used for the assay were subjected to anterograde injections of AAV9-PATZ1, followed by a thoracic crush injury one week later. Hi-C was performed on two types of tissues: mice at 7 days post-injury, and injured mice injected with AAV9-PATZ1, with the protocol individually performed for each tissue type. Hi-C was carried out following the in situ Hi-C protocol provided by the ENCODE Consortium. Briefly, the motor cortices (35-40mg, from 3 to 5 animals) were isolated from the mouse brain, and the tissues were crosslinked with 1% formaldehyde (Thermofisher, 12755) for 10 minutes at room temperature. After quenching with 2.5M glycine solution (MP Biomedicals, 808822) to a final concentration of 0.2M for 5 minutes, nuclei were isolated through cell lysis using cold Hi-C lysis buffer (5mM CaCl2, 3mM MgAc, 2mM EDTA, 0.5mM EGTA, 10mM Tris-HCl pH8.0, 1mM DTT, 0.1mM PMSF) with protease inhibitors (MCE, HY-K0010). After density gradient centrifugation with 1M Sucrose solution, nuclei were resuspended in 0.5% SDS and incubated at 62°C for 10 minutes. SDS was quenched with 10% Triton X-100, and chromatin was digested with 100U of MboI restriction enzyme (NEB, R0147) at 37°C overnight. Following enzyme inactivation at 62°C for 20 minutes, DNA ends were filled in with a mix containing biotin-14-dATP (0.4mM) (Thermo Scientific, 19524-016), dCTP, dGTP, dTTP (10mM each) (Thermo Scientific, 18254011, 18253013, 18255018) and DNA Polymerase I Klenow Fragment (NEB, M0210) at 37°C for 45 minutes. Ligation was performed in a master mix containing T4 DNA ligase buffer, 10% Triton X-100, 20mg/ml BSA (Catalog no: A7906-Sigma), and T4 DNA Ligase (NEB, M0202) at room temperature for 4 hours. After proteinase K (Sigma, P2308) treatment and crosslink reversal at 65°C overnight, DNA was purified, sheared using Covaris (M220, Condition Peak Incident: 50, Duty Factor= 10, Cycles/Burst=350, Treatment Time= 70), and size-selected for 300-500bp fragments using AMPure XP beads (Beckman Coulter, A63881). Biotin-labeled fragments were captured with Streptavidin T1 beads (Invitrogen, 65601), end-repaired, dA-tailed, and ligated to Illumina adapters (Illumina TruSeq UD Indexes, 20040870). The final library was PCR amplified (Illumina, 20015964), purified again with AMPure XP beads, and sequenced to a depth of ∼500 million paired-end reads on the Illumina NovaSeq 6000 platform.

### Cleavage Under Targets and Release Using Nuclease Sequencing (CUT&RUN)

Mice (n=3 per experimental group) received anterograde intracortical injections of AAV9-PATZ1, followed by thoracic spinal cord injury seven days later. CUT&RUN was performed using the DNA-Protein Interaction Assay Kit (Cell Signalling Technology, #86652) according to the manufacturer’s protocol. Motor cortices were dissected and fixed with 0.1% formaldehyde (Cell Signalling Technology, #12606P), then quenched using 10x glycine (Cell Signalling Technology, #7005S). Tissue was dissociated into cell suspensions using a Dounce homogenizer (Sigma, D8938) in wash buffer containing spermidine (Cell Signalling Technology, #27287) and protease inhibitors (Cell Signalling Technology, #7012). Cells were bound to activated Concanavalin A beads and incubated overnight at 4°C with the PATZ1 antibody (MyBioSource, MSB9403528).

The following day, pAG-MNase enzyme was added in a buffer premix (Cell Signalling Technology, #15338) containing digitonin (Cell Signalling Technology, #16359), spermidine, and protease inhibitors, and incubated with the sample for 1 hour. DNA cleavage was activated on ice by adding CaCl₂ in a digitonin-containing buffer and incubating at 4°C for 30 minutes.

DNA fragments were released by adding Stop Buffer (Cell Signalling Technology, #48105) containing digitonin and RNase A (Cell Signalling Technology, #7013), followed by a 10-minute incubation at 37°C. Crosslinks were reversed using 10% SDS and 20 mg/ml Proteinase K (Cell Signalling Technology, #10012), with incubation at 65°C for 2 hours. DNA was purified using DNA Purification Buffers and Spin Columns (Cell Signalling Technology, #14209) following the manufacturer’s recommended protocol. Libraries were prepared by end repair, adaptor ligation, and PCR amplification using the NEBNext® Ultra™ II DNA Library Prep Kit (NEB, #E7103).

Input samples were prepared using a similar protocol, except they were not treated with antibody or pAG-MNase. Instead, DNA was extracted using DNA Extraction Buffer (Cell Signalling Technology, #42015) for sonication (Covaris M220; 100–600 bp, Peak Incident: 75, Duty: 10, Cycles/Burst: 200, 75 s), followed by library preparation. Both CUT&RUN and input libraries were sequenced to 5 million paired-end reads (Illumina NovaSeq 6000).

Additionally, motor cortices were isolated from uninjured mice, mice 7 days post-injury, and mice treated with AAV-PATZ1 prior to thoracic injury. CUT&RUN assays were performed using the H3K27ac antibody (Abcam, ab4729) and CTCF antibody (Cell Signalling Technology, #3418), following the manufacturer’s protocol outlined above. These libraries were then amplified and sequenced to a depth of 5 million paired-end reads.

### Single-nuclei RNA Sequencing

Single-nuclei RNA sequencing (snRNA-seq) was performed using the Chromium Next GEM Single Cell 3’ v3.1 (Dual Index) platform (10x Genomics) following the manufacturer’s protocol (Document CG000315 Rev F). Nuclei were isolated from the motor cortex of mice (n=3 per experimental group) injected with AAV-GFP (control) and AAV-PATZ1 according to the nuclei isolation procedure mentioned above. It was then FACS sorted to isolate only GFP and DAPI-positive nuclei (DAPI used here as a nuclear marker to select intact nuclei, not as a viability exclusion dye) to isolate only GFP and DAPI (F6057-Sigma Aldrich) positive nuclei population. The cell suspensions were loaded onto the Chromium Next GEM Chip G (PN-2000177) for generating Gel Beads-in-Emulsion (GEM). Single cell suspensions were combined with Master Mix containing RT Reagent B (PN-2000165), Template Switch Oligo (PN-3000228), Reducing Agent B (PN-2000087), and RT Enzyme C (PN-2000085/2000102). A total of approximately 4000 cells were targeted. Cell suspensions and Master Mix were loaded into the Chromium Next GEM Chip G along with Single Cell 3’ v3.1 Gel Beads (PN-2000164) and Partitioning Oil (PN-2000190).

The chip was processed on the Chromium Controller to generate GEMs. Within each GEM, cells were lysed, and mRNAs were captured by poly(dT) oligonucleotides attached to the gel beads, which also contained a 16-nucleotide 10x Barcode, a 12-nucleotide Unique Molecular Identifier (UMI), and adaptor sequences. Reverse transcription was performed within the GEMs using the following thermal cycling protocol: 53°C for 45 minutes, followed by 85°C for 5 minutes. Following reverse transcription, GEMs were broken using Recovery Agent (PN-220016), and the barcoded, full-length cDNA was purified using Dynabeads MyOne SILANE (PN-2000048). Cleanup Buffer (PN-2000088) and Reducing Agent B (PN-2000087) were used in the Dynabeads Cleanup Mix. The purified cDNA was amplified using cDNA Primers (PN-2000089) and Amp Mix (PN-2000047/2000103) with the following cycling parameters mentioned in the protocol. The number of PCR cycles was optimised based on the targeted cell recovery, with 12 cycles used for 500-6,000 cells. Amplified cDNA was cleaned up using SPRIselect reagent (Beckman Coulter, PN B23318) at a ratio of 0.6X, and the purified cDNA was quality-checked and quantified using the Agilent Tapestation 4200 using D1000 tape (PN-5067-5586). All data are hosted at https://patz1.netlify.app/ for easy access.

For library construction, 25% of the total cDNA yield was used as input. The cDNA was enzymatically fragmented using Fragmentation Enzyme (PN-2000090/2000104) and Fragmentation Buffer (PN-2000091), followed by end-repair and A-tailing. The fragmentation reaction was incubated at 32°C for 5 minutes, followed by 65°C for 30 minutes. Size selection was performed using SPRIselect reagent with a double-sided protocol (0.6x followed by 0.8x) to obtain fragments of optimal size. Adaptor ligation was performed using Ligation Buffer (PN-2000092), DNA Ligase (PN-220110/220131), and Adaptor Oligos (PN-2000094) at 20°C for 15 minutes. After another round of cleanup using SPRIselect (0.8x), Sample Index PCR was performed using Amp Mix (PN-2000047/2000103) and dual indices from the Dual Index Kit TT Set A (PN-1000215, containing index plate PN-3000431). PCR cycling conditions were: 98°C for 45 seconds, followed by 12 cycles of (98°C for 20 seconds, 54°C for 30 seconds, 72°C for 20 seconds), with a final extension at 72°C for 1 minute. The final libraries were purified using a double-sided size selection with SPRIselect (0.6x followed by 0.8x) to remove adapter dimers and were eluted in Buffer EB (Qiagen, PN-19086).

Libraries were quality-checked using an Agilent Tapestation with D1000 tape to verify fragment size distribution (∼200-2,000 bp with a peak around 400 bp). Libraries were sequenced on an Illumina NovaSeq 6000 sequencer with a depth of 300 million paired-end reads.

### Adeno-Associated Virus Production

Adeno-associated viral (AAV) particles, serotype 9, were packaged and produced in HEK293T cells and then purified using a commercial AAV purification kit (Takara, #6666). Briefly, HEK293T cells were transiently transfected with pAAV-CAG-Patz1, pAAV2/9n (Addgene, #112865), and pAdDeltaF6 (Addgene, #112867) using Transfection Grade Linear Polyethylenimine Hydrochloride (MW 40,000) (Polysciences, #24765). At 48 hours post-transfection, cells were dislodged and transferred to sterile 50 ml falcon tubes. Viral isolation (Takara, #6666) was performed following the manufacturer’s instructions. To assess AAV capsid formation, proteins were separated on a 10% SDS-PAGE gel at 35 mA for 120 minutes. The gel was stained with Coomassie Brilliant Blue (Himedia, #17524.02) for 20 minutes, followed by overnight destaining in a solution of 10% acetic acid (Himedia, #90868) and 40% methanol (Himedia, #AS059) in water. SDS-PAGE analysis confirmed the presence of all three AAV capsid proteins: VP1 (83kDa), VP2 (73kDa), and VP3 (62kDa), with VP3 being the most abundant, as expected for correctly assembled AAV particles. For viral titer quantification, qPCR was performed using the Takara TB Green Premix Ex Taq II kit (Takara, #RR82WR). Viral particles were first treated with DNase for 30 minutes at 37°C, then serially diluted ten-fold across four dilutions. The viral titer was estimated by comparing the Ct values of the viral particles with a known plasmid standard (Addgene Plasmid, #59462) from a standard curve. All AAV preparations yielded titers of 10^11 viral genomes per ml or higher, with intact capsid formation confirmed by SDS-PAGE analysis.

### Quantitative Reverse Transcription PCR (qRT-PCR)

To validate PATZ1 overexpression at the mRNA level, total RNA was extracted from motor cortices of AAV-GFP injected (control) and AAV-GFP + AAV-PATZ1 co-injected (1:3 ratio) mice at 7 days post-injection. Motor cortices were dissected and homogenised in TRIzol reagent (Takara, 9108) following the manufacturer’s protocol (NucleoSpin RNA extraction kit, 740955). RNA concentration and purity were assessed spectrophotometrically (260/280 ratio). Complementary DNA (cDNA) was synthesised with 800ng of RNA using a standard reverse transcription kit with oligo(dT) primers according to the manufacturer’s instructions (Takara, Prime script 1st strand cDNA synthesis kit, 6110A)

Quantitative PCR was performed using a SYBR Green-based detection system (Takara TB Green Premix Ex Taq II, RR820). The following primer pairs were used: mouse Actb (β-actin, reference gene): forward 5’-GCATCTTCTTGTGCAGTGCC-3’, reverse 5’-TACGGCCAAATCCGTTCACA-3’; mouse PATZ1: forward 5’-ATGGCAGACCACCTGAAGAAGC-3’, reverse 5’-GTTTCTCTGGCGTCTTCAGGTC-3’. Each reaction was performed in triplicate. Cycling conditions consisted of an initial denaturation step followed by 40 cycles of amplification, with a final melt curve analysis to confirm primer specificity and the absence of non-specific amplification products.

Relative mRNA expression was calculated using the 2^(−ΔΔCt) method. Briefly, the ΔCt for each sample was calculated by subtracting the Ct value of the reference gene (Actb) from the Ct value of PATZ1. The ΔΔCt was then calculated by subtracting the mean ΔCt of the AAV-GFP control group from the ΔCt of each AAV-PATZ1 sample. Fold change in PATZ1 expression relative to control was expressed as 2^(−ΔΔCt).

### Curation of Growth-Associated Promoter and Enhancer Sets

A set of 291 genes was assembled using a literature-guided approach, selecting genes with documented expression or functional relevance in regeneration-competent neurons, including dorsal root ganglion sensory neurons following peripheral axotomy and zebrafish retinal ganglion cells following optic nerve crush[2,3,58,63–68]. Genes were required to fall within one or more of the following functional categories, chosen because they represent distinct mechanistic barriers or enablers of axon regrowth: axon guidance, cytoskeleton organisation, regulation of cell growth, chromatin remodeling, DNA damage and repair, and nervous system development. Gene Ontology (GO) term annotations were used to verify category membership for each gene.

Promoter regions were defined as the 2 kb window centred on the annotated transcription start site (TSS) of each gene. Chromatin accessibility at these regions was quantified from ATAC-seq data as described below.

Enhancer–gene associations were derived from the Activity-By-Contact (ABC) algorithm applied from previously published co-occupancy study[20]. Briefly, the ABC model integrates chromatin accessibility (ATAC-seq) and H3K27ac signal with Hi-C contact frequency to assign distal regulatory elements to their most probable target genes. We retrieved the ABC-predicted enhancer set for our 291 curated genes from this published resource and used the resulting 1,228 enhancer coordinates to define the growth-associated enhancer set analysed throughout this study. We note that these enhancer–gene links represent computational predictions based on chromatin state and three-dimensional proximity, and have not been independently validated by reporter assay or perturbation in the current study.

To independently validate the curated gene set without prior assumptions, we performed a parallel unbiased analysis. All mouse genes annotated to GO terms relevant to neuronal growth and plasticity were retrieved, including axonogenesis (GO:0007409), axon guidance (GO:0007411), neuron projection guidance (GO:0097485), regeneration (GO:0031099), axon extension (GO:0048675), actin cytoskeleton organisation (GO:0030036), chromatin remodeling (GO:0006338), central nervous system development (GO:0007417), RNA polymerase II-dependent transcription (GO:0006366), signal transduction (GO:0007165), and neurogenesis (GO:0022008). Accessibility heatmaps generated from this unbiased gene set were consistent with those from the curated list (Supplementary Figure S1B), supporting the robustness of the curation strategy.

To further assess whether the observed developmental restriction of chromatin accessibility is selective for growth-associated loci rather than reflecting a global reduction during maturation, we performed an additional unbiased analysis. Specifically, we extracted genes associated with broad GO categories, including nervous system development, axon guidance, and cytoskeleton organisation, and examined their chromatin accessibility patterns across developmental stages. In parallel, we generated genome-wide accessibility heatmaps across all genes. These analyses confirmed that the observed accessibility decline is not a general feature of the genome but is preferentially enriched at growth-associated loci.

### Single Nuclear ATAC Sequencing Analysis Pipeline

Cell Ranger ATAC version 7.2 was used to process Chromium scATAC-seq data following protocols described previously [69]. Fragment files were imported into ArchR version 1.0.2 [70]. We applied stringent quality control filters, retaining only cells with >1,000 fragments and >4 transcription start site (TSS) enrichment score. After quality control and filtering of empty droplets and doublets, we identified approximately 72,000 peaks from 1,500 nuclei per sample. Gene activities for each nucleus were calculated using the GeneScoreMatrix() function by summing peak counts in the gene body plus 2 kilobase pairs (kb) upstream[70]. Data were normalized using term frequency-inverse document frequency (TF-IDF) normalization (RunTFIDF), followed by dimensionality reduction using singular value decomposition (RunSVD). K-nearest neighbors were calculated using FindNeighbours (reduction = “lsi”, dims = 2:30). Cell clusters were identified using a shared nearest neighbor (SNN) modularity optimization-based clustering algorithm FindClusters (algorithm = 3, resolution = 2). UMAP visualization was generated using the RunUMAP function with reduction = “lsi” and dims = 2:30. Peaks were called using the addReproduciblePeakSet() function in ArchR using the Tiles matrix method with a fixed width of 501 base pairs. Differential chromatin accessibility between conditions was assessed using ArchR’s built-in statistical framework via the getMarkerFeatures() function. Peaks were then filtered using a threshold of adjusted p-value < 0.05 and an absolute Log2 fold-change > 0.5 to identify significantly upregulated or downregulated regions.

The complete automated analysis workflow is documented in detail on our GitHub repository https://github.com/VenkateshLab/Targeted-chromatin-remodeling-by-PATZ1. We utilised in R shiny app to provide pre-built clusters of the analysed snATAC-seq data, enabling direct visualization and interactive exploration. The clustered data is publicly accessible through our website (https://patz1.netlify.app/).

### Bulk ATAC Sequencing Analysis Pipeline

Mouse forebrain samples were collected from the ENCODE[37] and in-house data sets (P4 and P7) project at various developmental stages, including embryonic days E11 (ENCSR273UFV), E12 (ENCSR559FAJ), E13 (ENCSR903GMO), E14 (ENCSR810HQR), and E18 (ENCSR836PUC), as well as postnatal stages P0 (ENCSR310MLB), P4, P7 and adult (P36) (ENCSR976LWP). Additional Bulk ATAC-seq datasets were generated for other developmental stages as described above. Paired-end reads were trimmed using fastPto remove adapter sequences and low-quality bases [71]. Trimmed reads were aligned to the mouse genome (mm10) using the STAR version 2.7.11a aligner with default parameters [72]. We filtered out reads with mapping quality scores lower than 10 using SAMtools version 1.19 and removed duplicate reads using the MarkDuplicates function from the Picard toolkit [73]. For downstream analysis, we employed the TOBIAS-snake pipeline with default parameters [74]. BigWig files were generated for visualization in the UCSC Genome Browser [75]. The complete automated pipeline details are available in our GitHub repository https://github.com/VenkateshLab/Targeted-chromatin-remodeling-by-PATZ1.Furthermore, the bed files from Embryonic, Postnatal, and Adult stages were used for Transcription Factor Co-Occurrence analysis using TF-COMB[76].

### Batch Correction

To associate chromatin accessibility with specific genes, ATAC-seq reads were quantified at TSS-proximal regions using featureCounts [77] with exon-level annotations from the mm10 genome, generating per-sample read count matrices for E11–E18, P0, P4, P7, and Adult samples. Batch correction was performed on these TSS-proximal count matrices using ComBat-seq from the sva R package [78], which employs a negative binomial regression framework designed for sequencing count data. Developmental stage was specified as the biological covariate to ensure that stage-specific variation in chromatin accessibility was preserved during correction. FPKM values were subsequently computed from the corrected counts to normalize for both library size and feature length, enabling gene-level comparisons of TSS accessibility across developmental stages. PCA was performed on both the raw and batch-corrected matrices to evaluate the effectiveness of the correction. The first principal component accounted for 29.49% of the variance prior to correction and 31.56% after correction, confirming that the developmental signal was preserved.

### CUT&RUN Analysis Pipeline

Raw sequencing reads were processed with fastP for adapter removal and low-quality read trimming. Clean reads were aligned to the mouse genome (mm10) using BWA-mem version 0.7.19 with default parameters[79]. Aligned reads were converted to BAM format and sorted using SAMtools, and PCR duplicates were removed with Picard. Peak calling was performed using MACS2 with default parameters[80]. Identified peaks were annotated using GenomicRanges and ChIPseeker packages in R version 4.4.2 [81]. The annotated BED files were further analysed to identify CTCF binding sites and enhancer regions using resources from the SCREEN project from ENCODE[37]. The complete automated analysis workflow is documented in detail on our GitHub repository mentioned above.

### snRNA-seq Data Analysis Pipeline

Our workflow began with raw data quality assessment using FastQC ensuring sequencing data met basic quality thresholds for base quality, adapter contamination, and overall read quality. Following quality assessment, we used FastPto remove low-quality reads, filtering out those with poor quality scores or excessive adapter sequences [71]. The cleaned data were processed using CellRanger, which was specifically designed for single-nucleus RNA-seq data alignment and quantification [82]. We used the mm10 mouse reference genome with default parameters to generate high-quality alignments. Samples were sequenced to an average depth of ∼70,000 reads/nucleus (3,000-4,000 nuclei per library) on an Illumina NovaSeq 6000 platform.

The datasets were integrated using Seurat v5 [83] and differential expression analysis was conducted using the “FindMarkers” function, applying a nonparametric Wilcoxon rank-sum test. Cluster visualization in Seurat was generated using the “scCustomize” package in R (https://github.com/samuel-marsh/scCustomize), while MA-plots, histograms, and heatmaps were created using the “ggplot2” package (https://ggplot2.tidyverse.org/). We utilised ShinyCell in R to provide pre-built clusters of the analysed scRNA-seq data, enabling direct visualization and interactive exploration [84]. The clustered data is publicly accessible through our dedicated (https://patz1.netlify.app/).

### Hi-C Data Processing and Bioinformatic Pipeline Quality Control and Read Pre-processing

Raw paired-end sequencing reads (FASTQ format) from the PATZ1_Injured and Injured (control) Hi-C libraries were assessed for quality using FastQC (v0.11.9) [85], and per-sample reports were aggregated using MultiQC [86]. To remove technical artifacts and low-quality sequences prior to alignment, Hi-C chimeric reads were trimmed at MboI restriction sites (GATC) using homerTools (HOMER v4.11) [87]with the parameters −3 GATC -mis 0 - matchStart 20 -min 20. This step ensures that only the informative portion of each chimeric read is carried forward to alignment, reducing spurious ligation products.

### Alignment and Tag Directory Construction

Trimmed paired-end reads were aligned independently to the mouse reference genome (GRCm38/mm10) using Bowtie2 (v2.4.2) [4] in end-to-end mode with the --very-sensitive preset. Independent alignment of each mate pair is a standard requirement for Hi-C data to avoid suppression of valid trans-chromosomal contacts. Valid Hi-C read pairs were then parsed and organized into HOMER Tag Directories using the makeTagDirectory command [87].

Several stringent filtering criteria were applied to remove technical noise and ensure data quality: (i) PCR duplicates were collapsed by restricting reads to one tag per base pair (-tbp 1); (ii) self-ligation fragments, reads terminating at restriction sites, and background pair-end reads were discarded (-removePEbg -removeSelfLigation -removeRestrictionEnds); (iii) sequencing spikes exceeding 10,000 tags per bin were removed (-removeSpikes 10000 5); and (iv) GC-content bias was corrected during normalization (-checkGC). These filtering parameters are consistent with established best practices for Hi-C data processing [87,88].

### Identification of Chromatin Architectural Features

A/B Compartment Analysis. Chromatin compartments were identified by performing Principal Component Analysis (PCA) on normalized Hi-C contact matrices using the HOMER command runHiCpca.pl [3]. The first principal component (PC1) was computed at 25 kb resolution using a 50 kb sliding window. Genomic bins with PC1 > 0 were assigned to the transcriptionally active ‘A’ compartment, while bins with PC1 < 0 were assigned to the inactive ‘B’ compartment, consistent with the convention established by Lieberman-Aiden et al. [89].

TAD and Loop Identification. Topologically Associating Domains (TADs) and chromatin loops were initially called using HOMER’s findTADsAndLoops.pl [87] at high-resolution (3 kb bin size, 15 kb window). To enhance loop detection sensitivity, Peakachu was applied as an orthogonal, supervised machine learning-based loop caller. Peakachu[90] employs a Random Forest classification framework trained on orthogonal chromatin interaction data (e.g., ChIA-PET or HiChIP) to predict chromatin loops de novo from genome-wide Hi-C contact maps, enabling detection of short-range interactions that enrichment-based methods may miss [90]. Loop calls supported by both HOMER and Peakachu were retained as high-confidence interactions for downstream analysis.

High-Resolution Visualization. For visualization and comparative inspection of Hi-C contact maps, HOMER Tag Directories were converted to .hic format files using tagDir2hicFile.pl and Juicer Tools (v1.22.01)[91]. Contact maps were visualized using Juicebox[91].

### Comparative Genomic Analysis and Annotation

To identify PATZ1-induced 3D architectural changes, TAD and loop 2D BED files from both conditions (PATZ1_Injured vs. Injured) were merged using merge2Dbed.pl [3]. Differential domain insulation strength and loop contact frequency were quantified using the findTADsAndLoops.pl score function. Dynamic compartment regions were isolated by comparing per-bin PC1 values between conditions from bedGraph files to classify regions as B→A (transcriptional activation) or A→B (transcriptional repression) transitions. Genomic annotation of compartment-shifting bins, differential TAD boundaries, and loop anchors was performed using annotatePeaks.pl [3]. All coordinates are referenced to the mm10 assembly.

### Integration of Hi-C, snATAC-seq, and snRNA-seq Data

To link three-dimensional genome architecture with chromatin accessibility and gene expression, Hi-C compartment data were integrated with single-nucleus ATAC-seq (snATAC-seq) and single-nucleus RNA-seq (snRNA-seq) datasets from the same experimental conditions (PATZ1_Injured and Injured control). The primary analytical unit was genomic regions undergoing A/B compartment switching, which were then interrogated for concordant changes in local chromatin accessibility and transcriptional output.

### Hierarchical Multi-omic Classification (The ‘5-Rule’ System)

Compartment-switching regions were classified into five hierarchical evidence categories based on the co-occurrence of architectural and functional features within overlapping genomic coordinates. ATAC-seq peaks were associated with a gene if they fell within the gene body or within a ±50 kb flanking window. The five categories are defined as:

1. Category 1 (C1): Compartment shift + TAD boundary change + snATAC-seq peak + snRNA-seq differentially expressed gene (DEG).
2. Category 2 (C2): Compartment shift + TAD boundary change + snRNA-seq DEG.
3. Category 3 (C3): Compartment shift + TAD boundary change + Hi-C loop.
4. Category 4 (C4): Compartment shift + TAD boundary change + Hi-C loop + snRNA-seq DEG.
5. Category 5 (C5): Compartment shift + TAD boundary change + Hi-C loop + snATAC-seq peak + snRNA-seq DEG (highest confidence).

### Statistical Analysis and Visualization

Integrated multi-layer heatmaps were constructed to display the relationship between PC1 values (Injured vs. PATZ1), TAD insulation scores, snATAC-seq peak counts, and ATAC log2 fold-change (L2FC), with genes sorted by ATAC LFC to highlight concordance between architectural and accessibility shifts. The distribution of snATAC-seq LFC values was compared between B-to-A and A-to-B shifting regions using probability density histograms; statistical significance was assessed using the two-sided Mann-Whitney U (MWU) test[92]. Architectural shift regions were consistently color-coded throughout all figures: green for B-to-A (activation/opening) and red for A-to-B (repression/closing). All manuscript figures were generated at 600 DPI.

### Integration of Bulk ATAC-seq and Bulk RNA-seq Across Developmental Timepoints Data Integration and Pre-processing

Publicly available bulk ATAC-seq and bulk RNA-seq datasets spanning mouse cortical development (embryonic day 11 [E11] through adult) were retrieved from ENCODE [93]and integrated by matching gene symbols across modalities. Mean chromatin accessibility and mean gene expression values were calculated for each gene across all eight developmental timepoints (E11, E12, E13, E14, E16, E18, P0, and Adult). Both modalities were log2-transformed [log2(mean + 1)] prior to correlation analysis and density scatter plotting. For heatmap visualization, row-wise Z-score scaling was applied independently to each modality to enable direct comparison of temporal dynamics [94].

### Chromatin Accessibility and Gene Expression Correlation

The global relationship between chromatin state and transcriptional output was quantified using Spearman’s rank correlation between log2-transformed mean accessibility and expression values [95]. Correlation was visualized using density-weighted hexbin scatter plots. Developmental stage-specific patterns were further examined by grouping samples into three categories: Embryonic (E11–E18), Postnatal (P0), and Adult.

### Quadrant Analysis and Gene Categorization

Genes were classified into four functional quadrants defined by their position relative to the global median of mean accessibility and expression across the full dataset: (i) Open/Expressed (Q1, high accessibility and high expression); (ii) Closed/Expressed (Q2, low accessibility but high expression); (iii) Closed/Low (Q3, low accessibility and low expression); and (iv) Open/Low (Q4, high accessibility but low expression), which may represent developmentally poised or primed regulatory states. The distribution of genes across these quadrants was quantified using 2×2 contingency tables for both the complete gene set and a curated ‘Progrowth’ gene list.

### Heatmap Visualization and Hierarchical Clustering

Hierarchical clustering of Z-score scaled data was performed using Euclidean distance and the complete linkage method, implemented via the fastcluster package in R [13]. Heatmaps were generated using the ComplexHeatmap library[96], with a diverging color gradient from −2 (low, salmon) to +2 (high, light green). Column order was fixed to the chronological developmental sequence from Embryonic to Adult stages.

### Intra-Cortical (IC) Integrated ATAC-seq and RNA-seq Analysis Data Source and Pre-processing

Intra-cortical (IC) injury datasets were processed to characterise the relationship between chromatin accessibility and transcriptional output in the context of acute cortical injury. Differentially accessible regions (DARs) showing increased accessibility in IC-Injured versus Uninjured cortical samples were identified from snATAC-seq data and formatted as a BED file. Gene expression levels from IC samples were quantified using featureCounts (Subread package, v2.0) [77]applied to aligned RNA-seq reads, yielding a raw count matrix.

### Computational Integration Pipeline

Integration of accessibility and transcriptomic data was performed using a custom Python-based analytical framework. To examine how chromatin accessibility correlates with transcriptional output across the full expression dynamic range, the complete gene set was ranked by expression value and partitioned into ten equal-sized deciles (Expression Groups 1–10). Genes were classified as ‘accessible’ if at least one IC-Injured ATAC-seq peak overlapped their gene body or proximal regulatory region. The percentage of accessible genes was calculated for each expression decile, and a cumulative frequency distribution and barplot were generated to visualize the positive correlation between expression level and the frequency of open chromatin peaks. Parallel processing across deciles was implemented using Python’s multiprocessing module to ensure computational efficiency.

### Progrowth Gene Subset Analysis

A curated list of 292 ‘Progrowth’ genes was cross-referenced against the IC integrated dataset to determine whether regeneration-associated transcriptional programs are specifically linked to open chromatin architecture. Decile-based binning and accessibility calculations were repeated for the Progrowth gene subset and compared against the global genomic background. Venn diagrams were constructed to visualize the intersection between highly expressed genes, lowly expressed genes, and accessible Progrowth genes, quantifying the subset of growth-associated genes that are concurrently transcriptionally active and architecturally accessible in the IC-Injured state.

### Injury Response Visualization

Chromatin accessibility and gene expression were compared between Intact (Adult) and Injured (IC) conditions. RNA-seq differential expression was represented as average log2 fold change (Log2FC), while ATAC-seq data were Z-score scaled across conditions and displayed as aligned heatmaps. Lollipop plots were used to visualize concordance or divergence between ATAC-seq and RNA-seq Log2FC for selected candidate genes. Spearman’s rank correlation was used to assess the degree of agreement between accessibility shifts and gene expression changes during the regenerative response [95].

### Software and Statistical Analysis

All data processing and statistical integration were conducted using Python (v3.10+) with the pandas, numpy, scipy, seaborn, and matplotlib libraries[97–100]. Heatmap generation and hierarchical clustering were performed in R (v4.3+) using ComplexHeatmap, circlize, and fastcluster[96,101,102]. The Mann-Whitney U test and Spearman rank correlation were calculated using scipy.stats[92,95].

