## Supplementary material for "PATZ1 Reinstates a Growth-Permissive Chromatin Landscape in Adult Corticospinal Neurons After Injury": Addtional File 2 FIgure S 1 - S24

### Additional File 2 – Supplementary Figure S1 – S24

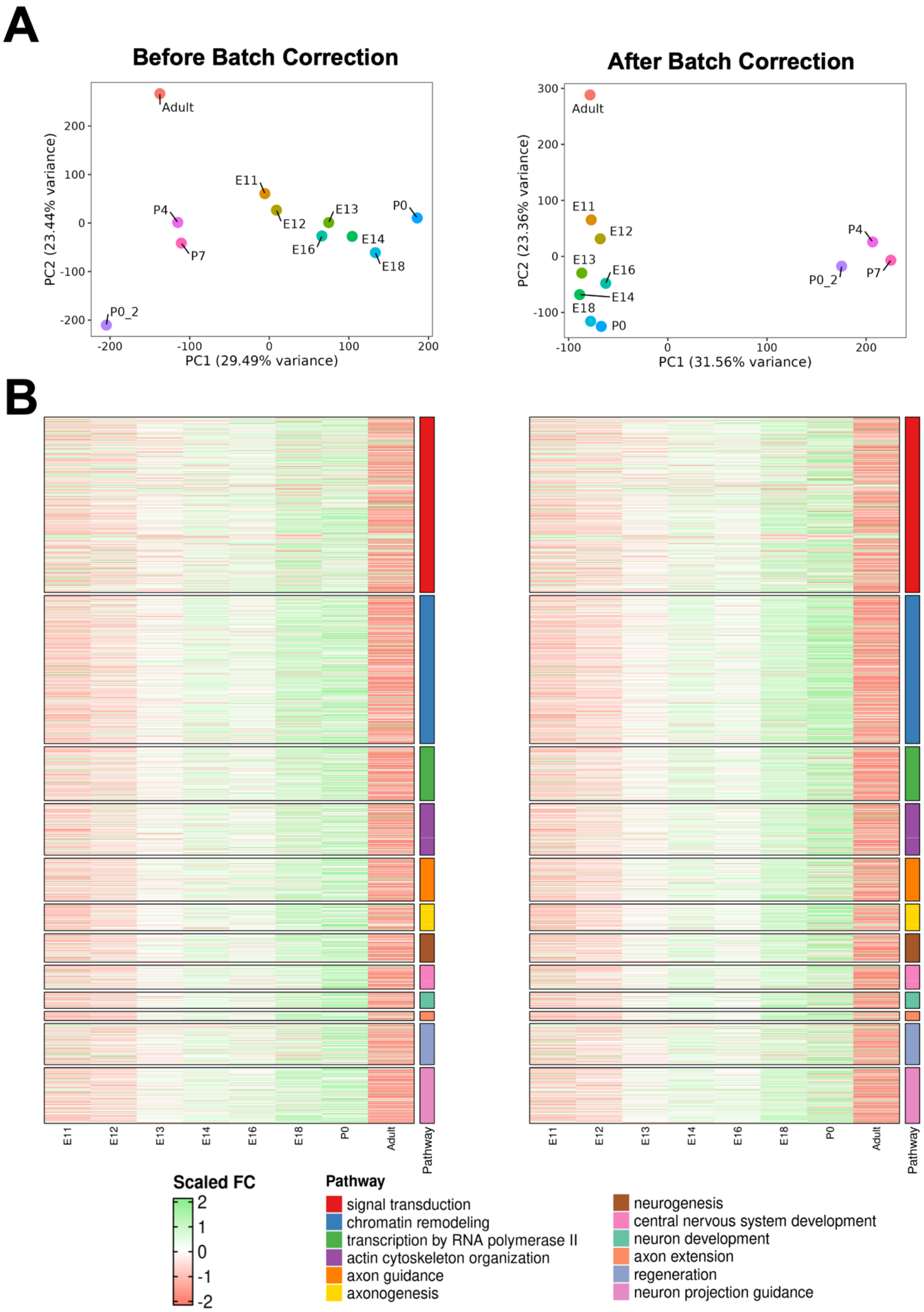
A
Before Batch Correction
After Batch Correction
PC1 (29.49% variance)
PC2 (23.44% variance)
PC1 (31.56% variance)
PC2 (23.36% variance)
Adult
E11
E12
E13
E14
E16
E18
P0
P0_2
P4
P7
B
Pathway
Scaled FC
signal transduction
chromatin remodeling
transcription by RNA polymerase II
actin cytoskeleton organization
axon guidance
axonogenesis
neurogenesis
central nervous system development
neuron development
axon extension
regeneration
neuron projection guidance

**Supplementary Figure S1: Principal Component Analysis (PCA) and Heatmap Analysis of Developmental and Postnatal Samples.**

**(A)** PCA plots before and after batch correction. The plot on the left shows the samples (E11, E12, E13, E14, E16, E18, P0, P4, P7, and Adult) before batch correction, while the plot on the right illustrates the same samples after batch correction. The first principal component (PC1) accounts for 29.49% of the variance before batch correction, and 31.56% of the variance after batch correction.

**(B)** Two side-by-side heatmaps (left before batch correction and right after batch correction) showing scaled fold-change (FC) across different samples based on 12 Gene Ontology (GO) terms. The GO terms, derived through unbiased text mining, include: signal transduction, chromatin remodeling, transcription by RNA polymerase II, actin cytoskeleton organization, axon guidance, axonogenesis, neurogenesis, central nervous system development, neuron development, axon extension, regeneration, and neuron projection guidance. Each row represents a sample, and the color intensity indicates the relative gene expression for each GO term.

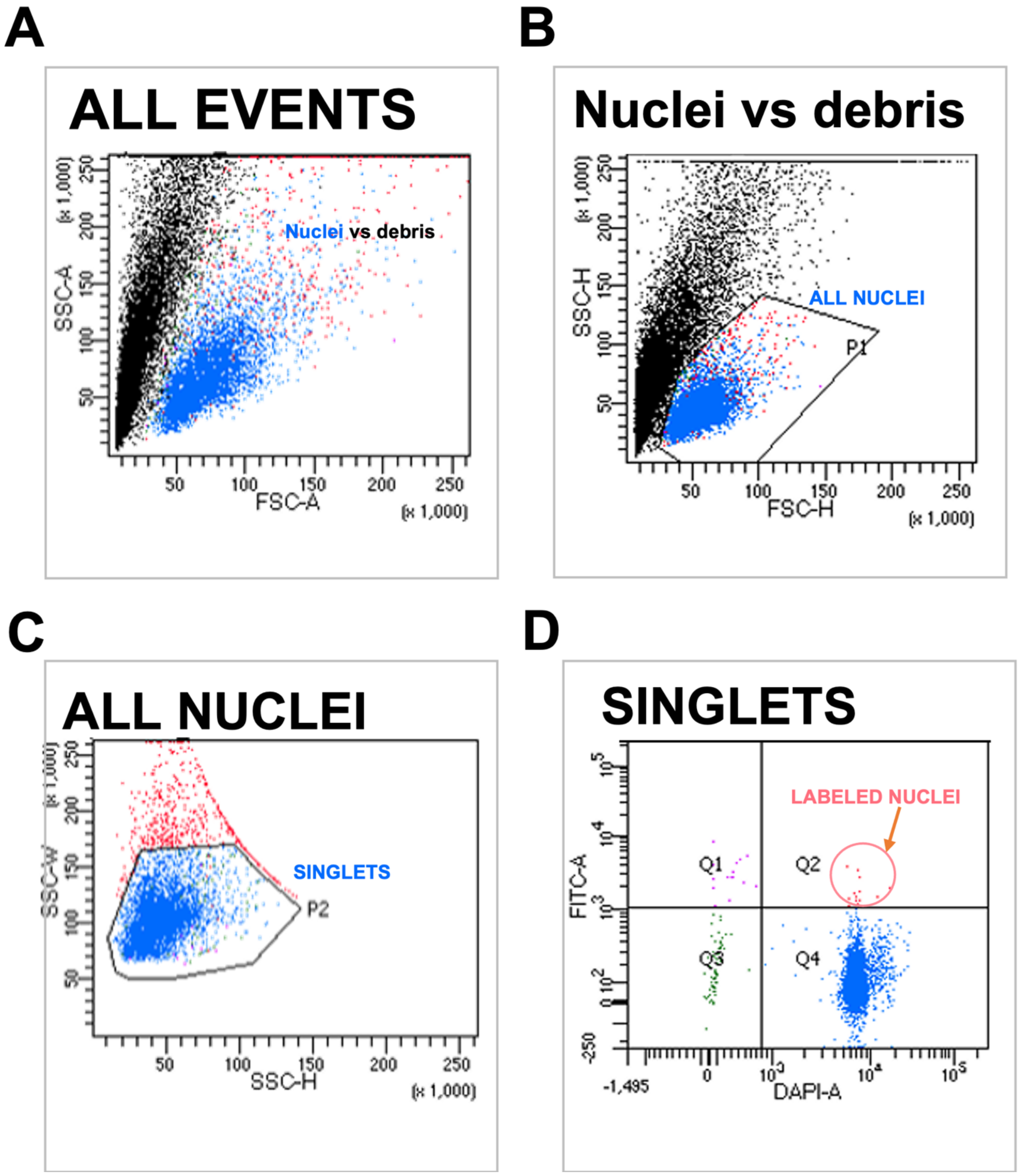


**Supplemental Figure S2: BD-FACS purification of GFP labeled cortical nuclei.**

**(A,B)** Nuclei were sorted by passing them through a series of filters based on forward and side scatter to distinguish them from cellular debris.

**(C)** minimize doublets and **(D)** then nuclei labeled by eGFP are detected in the FITC channel, red arrow

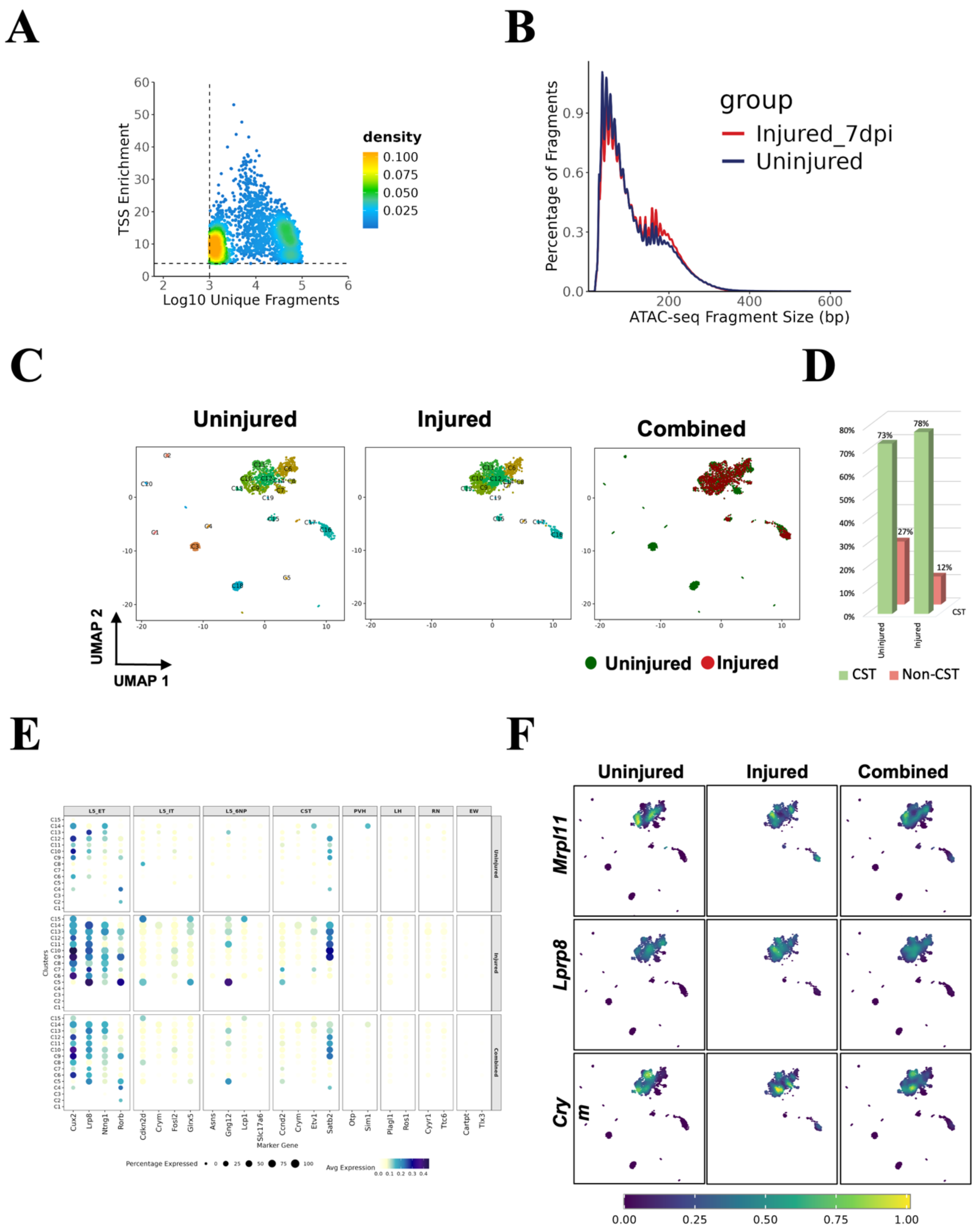


**Supplementary Figure S3: Extended snATAC-seq quality control metrics and chromatin accessibility landscape supporting Figure 3, comparing Uninjured and Injured spinal cord datasets.**

**(A)** snATAC-seq library quality metrics showing TSS enrichment score versus log10 unique fragments per cell. Density gradient (yellow = high, blue = low) confirms high-quality nuclei passing quality thresholds for downstream analysis.

**(B)** ATAC-seq fragment size distribution for Uninjured (blue) and Injured_7dpi (red) samples, demonstrating characteristic nucleosomal periodicity (mono-, di-, and tri-nucleosomal banding), confirming successful snATAC-seq library preparation.

**(C)** UMAP visualization of chromatin accessibility profiles from Uninjured (left), Injured (middle), and Combined (right) datasets. Clusters are color-coded by cell identity and labeled (C0–C19), enabling direct comparison of cluster distribution across conditions.

**(D)** Bar graph quantifying the proportion of CST (green) and Non-CST (red) cell populations in Uninjured and Injured conditions, showing 73% vs 27% and 78% vs 12%, respectively.

**(E)** Dot plot displaying cell type-specific chromatin accessibility at marker gene loci across identified clusters in Uninjured, Injured, and Combined conditions. Dot size reflects the percentage of cells with accessible chromatin at each locus; color intensity represents average accessibility (light = low, dark blue = high). Marker genes are grouped by cell lineage: LS_ET, LS_IT, LS_SNP, CST, PVH, LH, RN, and EW populations. (LS_ET — Layer 5 Extra-Telencephalic (corticospinal/corticobulbar projection neurons), LS_IT — Layer 5 Intra-Telencephalic (corticocortical/corticostriatal projection neurons)LS_SNP — Layer 5/6 Near-Projecting neurons, CST — Corticospinal Tract neurons, PVH — Paraventricular Hypothalamic neurons, LH — Lateral Hypothalamic neurons, RN — Red Nucleus neurons, EW — Edinger-Westphal nucleus neurons)

**(F)** Feature plots showing normalized chromatin accessibility of CST marker genes *Mrpl11*, *Lrp8*, and *Crym* across UMAP embeddings in Uninjured, Injured, and Combined conditions. Color scale ranges from low (dark purple) to high (yellow) accessibility, highlighting CST cluster-specific chromatin accessibility patterns between conditions.

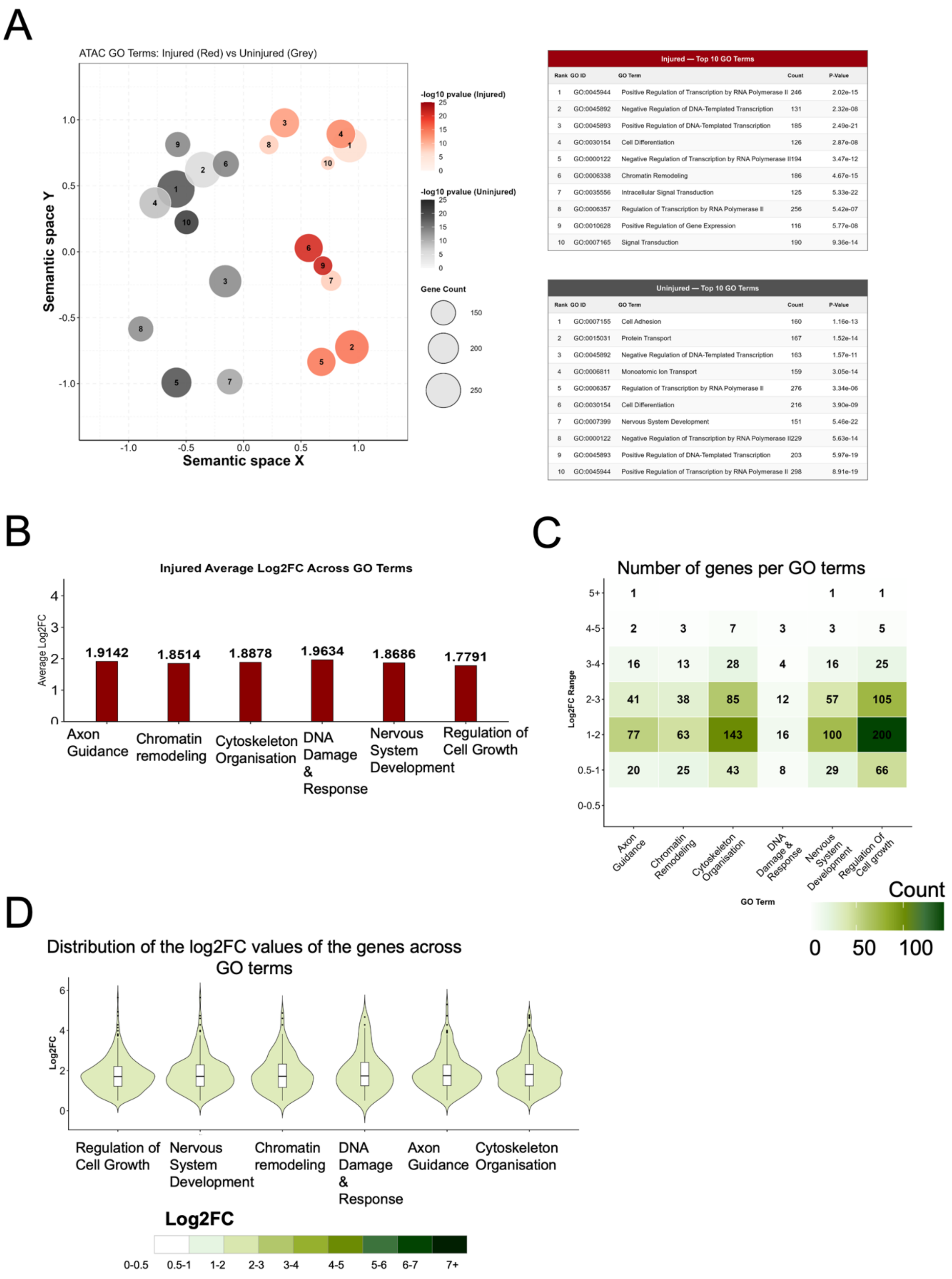


| Injured — Top 10 GO Terms | | | | |
|---|---|---|---|---|
| Rank | GO ID | GO Term | Count | P-Value |
| 1 | GO:0045944 | Positive Regulation of Transcription by RNA Polymerase II | 246 | 2.02e-15 |
| 2 | GO:0045892 | Negative Regulation of DNA-Templated Transcription | 131 | 2.32e-08 |
| 3 | GO:0045893 | Positive Regulation of DNA-Templated Transcription | 185 | 2.49e-21 |
| 4 | GO:0030154 | Cell Differentiation | 126 | 2.87e-08 |
| 5 | GO:0000122 | Negative Regulation of Transcription by RNA Polymerase II | 194 | 3.47e-12 |
| 6 | GO:0006338 | Chromatin Remodeling | 186 | 4.67e-15 |
| 7 | GO:0035556 | Intracellular Signal Transduction | 125 | 5.33e-22 |
| 8 | GO:0006357 | Regulation of Transcription by RNA Polymerase II | 256 | 5.42e-07 |
| 9 | GO:0010628 | Positive Regulation of Gene Expression | 116 | 5.77e-08 |
| 10 | GO:0007165 | Signal Transduction | 190 | 9.36e-14 |

| Uninjured — Top 10 GO Terms | | | | |
|---|---|---|---|---|
| Rank | GO ID | GO Term | Count | P-Value |
| 1 | GO:0007155 | Cell Adhesion | 160 | 1.16e-13 |
| 2 | GO:0015031 | Protein Transport | 167 | 1.52e-14 |
| 3 | GO:0045892 | Negative Regulation of DNA-Templated Transcription | 163 | 1.57e-11 |
| 4 | GO:0006811 | Monoatomic Ion Transport | 159 | 3.05e-14 |
| 5 | GO:0006357 | Regulation of Transcription by RNA Polymerase II | 276 | 3.34e-06 |
| 6 | GO:0030154 | Cell Differentiation | 216 | 3.90e-09 |
| 7 | GO:0007399 | Nervous System Development | 151 | 5.46e-22 |
| 8 | GO:0000122 | Negative Regulation of Transcription by RNA Polymerase II | 229 | 5.63e-14 |
| 9 | GO:0045893 | Positive Regulation of DNA-Templated Transcription | 203 | 5.97e-19 |
| 10 | GO:0045944 | Positive Regulation of Transcription by RNA Polymerase II | 298 | 8.91e-19 |

**Supplementary Figure S4. Alternative visualisations of snATAC-seq chromatin accessibility data from L5 ET neurons following spinal cord injury.**

**(A)** Bubble plot showing the top 10 enriched GO biological process terms from snATAC-seq analysis of L5 ET neurons in injured (red) and uninjured (grey) conditions. Each bubble represents one GO term, with bubble size proportional to the number of associated genes (Count) and colour intensity reflecting statistical significance (−log10 p-value). Terms are positioned in semantic space (X and Y axes) based on functional similarity, such that closely related terms cluster together. Separate colour scales are shown for injured (red gradient) and uninjured (grey gradient) conditions. The accompanying tables (right) list the top 10 terms for each condition, showing GO ID, term name, gene count, and p-value. This plot serves as an alternative to the radar chart shown in Figure 3E.

**(B)** Bar graph showing the average Log2 fold-change (Log2FC) of chromatin accessibility across six GO-term-defined biological pathways in injured L5 ET neurons: Axon Guidance, Chromatin Organization, Cytoskeleton Organization, DNA Damage Response, Nervous System Development, and Regulation of Cell Growth. Each bar represents the mean Log2FC of all differentially accessible peaks assigned to that pathway, with exact values indicated above each bar. All pathways display moderate chromatin opening (Log2FC ~1.8–2.0), suggesting broad and consistent upregulation of accessibility across functional gene categories. This plot serves as an alternative to the radar chart shown in Figure 3E.

**(C)** Heatmap showing the number of differentially accessible peaks binned by Log2FC magnitude across five biological pathways (Axon Guidance, Chromatin Remodelling, Cytoskeleton Organisation, DNA Damage, Nervous System Development). Rows represent Log2FC bins (0–0.5 to 5+, bottom to top) and columns represent GO term pathways. Colour intensity and numerical values within each tile reflect gene count per bin, with darker green indicating higher counts (colour scale, right). The 1–2 Log2FC bin contains the highest number of peaks across all pathways, consistent with moderate chromatin opening as the predominant injury response. This plot serves as an alternative to the circular bar plots shown in Figure 3G.

**(D)** Violin plots showing the distribution of Log2FC values for individual genes within each of the six biological pathways in injured L5 ET neurons. Each violin represents the kernel density estimate of Log2FC values, with an embedded boxplot indicating the median and interquartile range. Violin fill colour reflects the median Log2FC of each pathway (colour scale below; darker green = higher median accessibility). The broad distributions observed within each pathway highlight heterogeneity in the magnitude of chromatin opening across individual loci. This plot serves as an alternative to the heatmap shown in Figure 3F

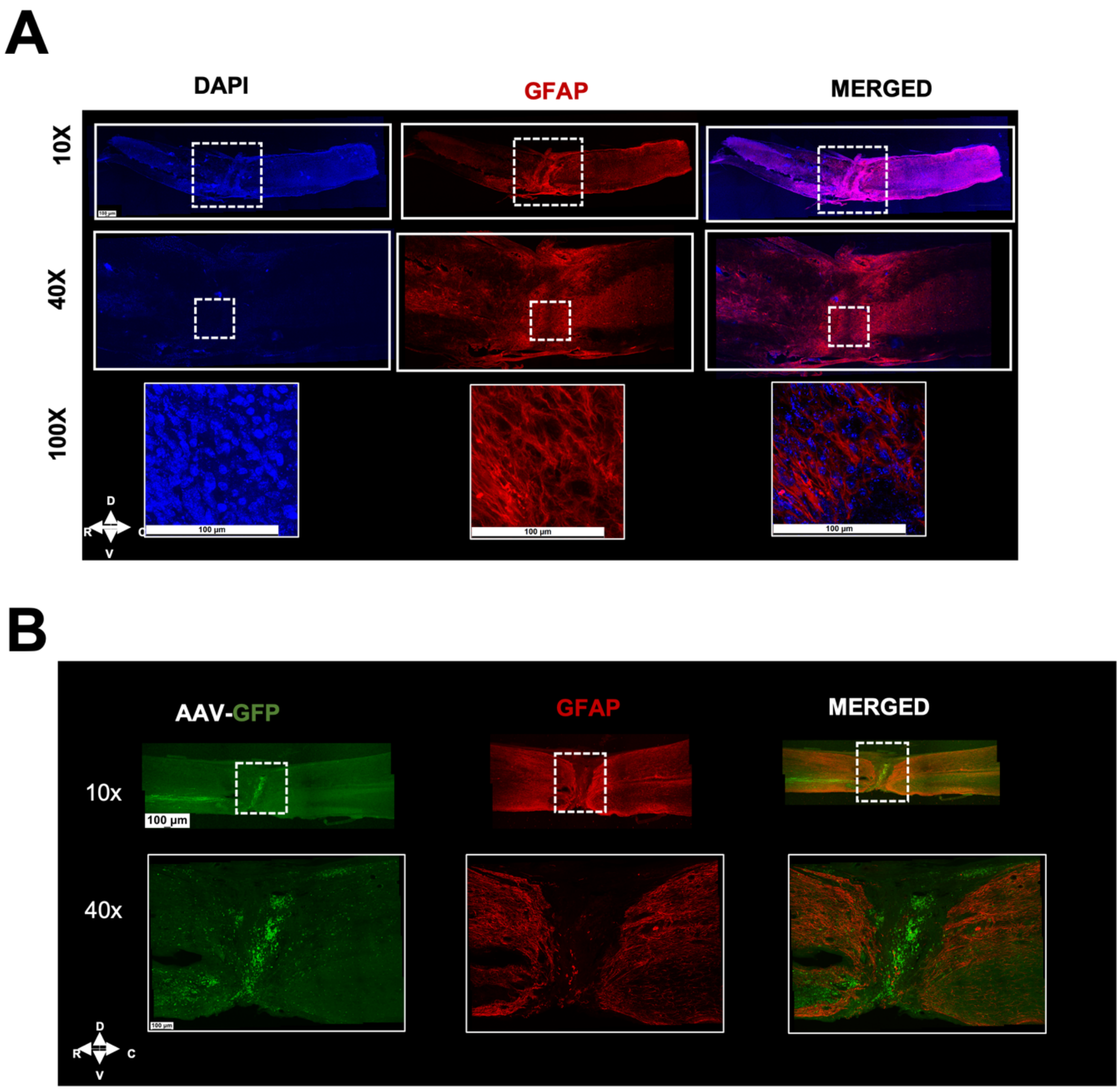


**Supplementary Figure S5. Manual thoracic crush injury produces consistent pathological features with clinically relevant variability.**

**(A)** Cross-sectional analysis at the lesion epicenter showing injury consistency across animals. DAPI (blue), GFAP (red), and merged images at progressive magnifications (10×, 40×, 100×) demonstrate that despite natural variability in absolute lesion size, all injuries exhibit consistent pathological hallmarks: defined lesion boundaries, central cavitation, and robust astrocytic response (GFAP+ reactive glia). This heterogeneity mirrors the variability observed in human spinal cord injuries.

**(B)** Longitudinal assessment of lesion extent showing AAV-GFP expression (green) and GFAP immunoreactivity (red) at the injury site. Images at 10× and 40× magnification reveal the rostral-caudal spread of injury. While individual lesions vary in absolute dimensions, all demonstrate consistent features including glial scar formation, tissue disruption at the epicenter (dashed lines), and preserved tissue architecture in adjacent segments.

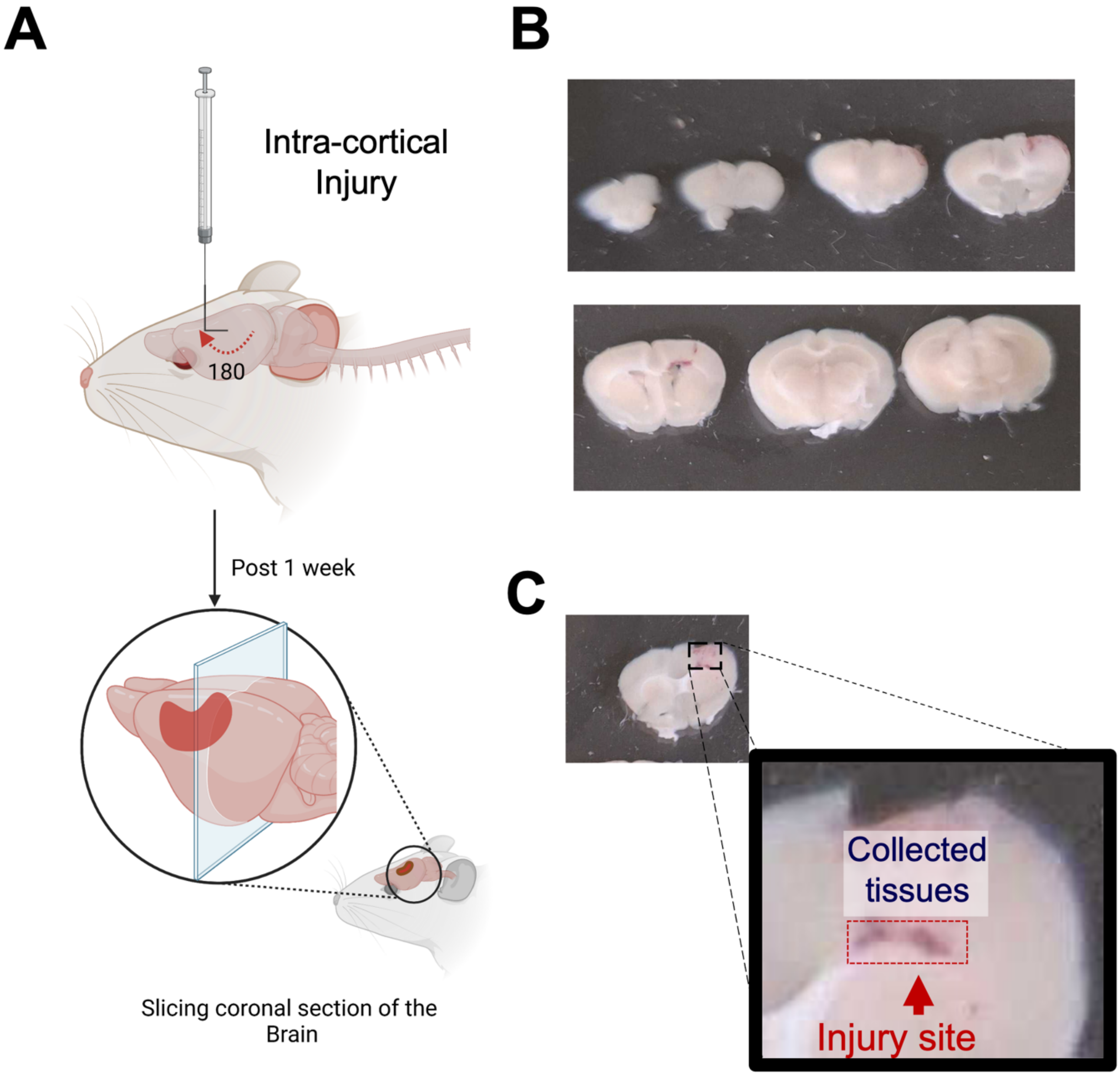


**Supplemental Figure S6: : Intracortical injury and the collection of CST neurons proximal to the Injury.**

**(A)** Experimental Design, a right-angle blade was inserted into the caudal cortex and rotated 180°, sweeping through a path just beneath the CST cell bodies to sever the axons

**(B)** The coronal slices of the Brain is isolated C. In coronal sections of the brain, the region above the hemorrhaged area (marked by red box) is collected for Bulk ATAC library preparations.

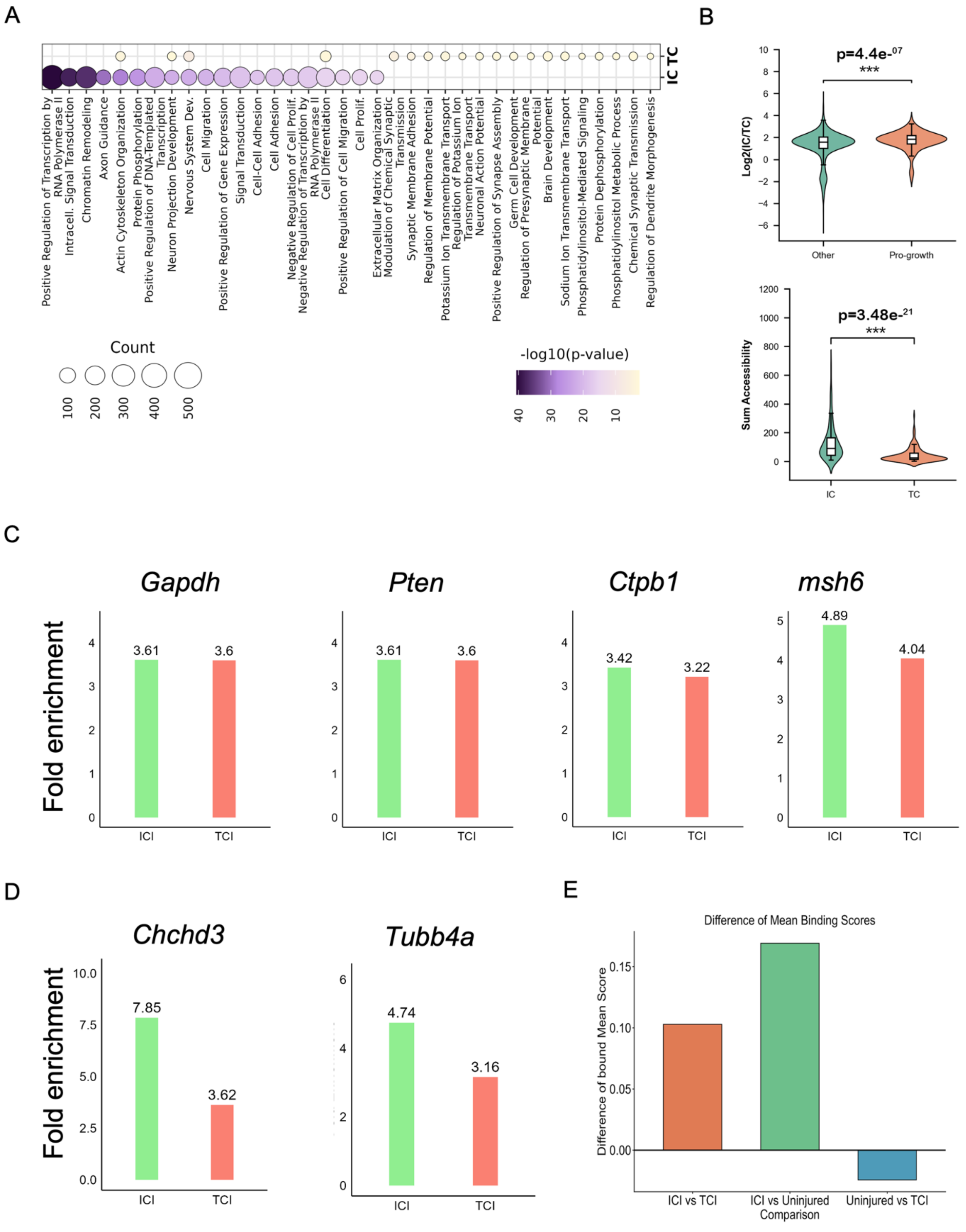


**Supplementary Figure S7. Intra-cortical injury drives disproportionately greater chromatin accessibility at pro-growth gene loci relative to thoracic crush injury, with preferential PATZ1 transcription factor occupancy.**

**(A)** Bubble dot plot showing the top GO biological process terms enriched in differentially accessible peaks comparing Intra-cortical Injury (ICI) versus Thoracic Crush Injury (TCI), determined by unbiased GO enrichment analysis ($p < 0.05$). Terms are displayed for both ICI (left column) and TCI (right column) conditions and sorted by gene count. Bubble size reflects the number of genes associated with each GO term (scale: 100–500 genes), and bubble colour indicates statistical significance expressed as –log10(adjusted p-value), with darker purple indicating greater significance (scale: 10–40). Pro-regenerative terms including Positive Regulation of Transcription by RNA Polymerase II, Chromatin Remodelling, Axon Guidance, and Actin Cytoskeleton Organisation rank among the most significant and gene-rich terms, with ICI showing markedly higher enrichment than TCI across these categories.

**(B)** Violin plots demonstrating that pro-growth gene loci exhibit significantly higher chromatin accessibility in ICI compared to TCI. The y-axis shows sum accessibility signal across accessible peaks at each locus; the x-axis shows condition (IC = green, TC = red/salmon). Internal box plots indicate median and interquartile range. (*Left*) Sum accessibility at pro-growth gene loci specifically labelled as axon growth/pro-growth genes (x-axis label: "Condition (Pro-growth Genes)"). (*Right*) Sum accessibility across the same loci shown without the x-axis label, representing a complementary view of the same comparison. Both plots show a highly significant increase in overall chromatin accessibility at pro-growth loci in ICI versus TCI (Wilcoxon rank-sum test, $p = 3.48e-21$, *** $p < 0.001$), confirming that the chromatin opening response at pro-growth gene loci is disproportionately stronger following injury proximal to the motor cortex.

**(C)** Bar graphs showing ATAC-seq fold enrichment at four non-pro-growth (housekeeping/control) gene loci — *Gapdh*, *Pten*, *Msh6*, and *Ctpb1* — comparing ICI (green) versus TCI (red). Fold enrichment values are displayed above each bar: *Gapdh* (ICI=3.61, TCI=3.60), *Pten* (ICI=3.61, TCI=3.60), *Msh6* (ICI=4.89, TCI=4.04), *Ctpb1* (ICI=3.42, TCI=3.22). In contrast to pro-growth genes, these non-growth genes show minimal difference in fold enrichment between ICI and TCI conditions, confirming that the enhanced chromatin accessibility observed in ICI is selective for pro-growth loci rather than a global, genome-wide effect.

**(D)** Bar graphs showing ATAC-seq fold enrichment at two representative pro-growth gene loci — *Chchd3* and *Tubb4a* — comparing ICI (green) versus TCI (red). Fold enrichment values: *Chchd3* (ICI=7.85, TCI=3.62) and *Tubb4a* (ICI=4.74, TCI=3.16). Both loci show substantially greater chromatin accessibility in ICI compared to TCI, consistent with the genome-wide patterns shown in the main figures and further validating the selective reopening of pro-growth gene chromatin following intra-cortical injury.

**(E)** Bar graph showing the difference in PATZ1 transcription factor bound mean score across three pairwise comparisons of injury conditions: ICI vs TCI (orange, $\Delta \approx 0.10$), ICI vs Uninjured (green, $\Delta \approx 0.17$), and Uninjured vs TCI (blue, $\Delta < 0$, approximately –0.01). The y-axis represents the difference in PATZ1 footprint bound mean score between conditions, derived from TF footprinting analysis of ATAC-seq accessible peaks. A positive value indicates higher PATZ1 occupancy in the first-named condition; a negative value indicates lower occupancy. ICI shows the greatest PATZ1 occupancy relative to both TCI and the uninjured state, while the Uninjured vs TCI comparison yields a near-zero or slightly negative difference, indicating that PATZ1 binding is specifically elevated in the context of intra-cortical injury and is not simply a consequence of injury per se.

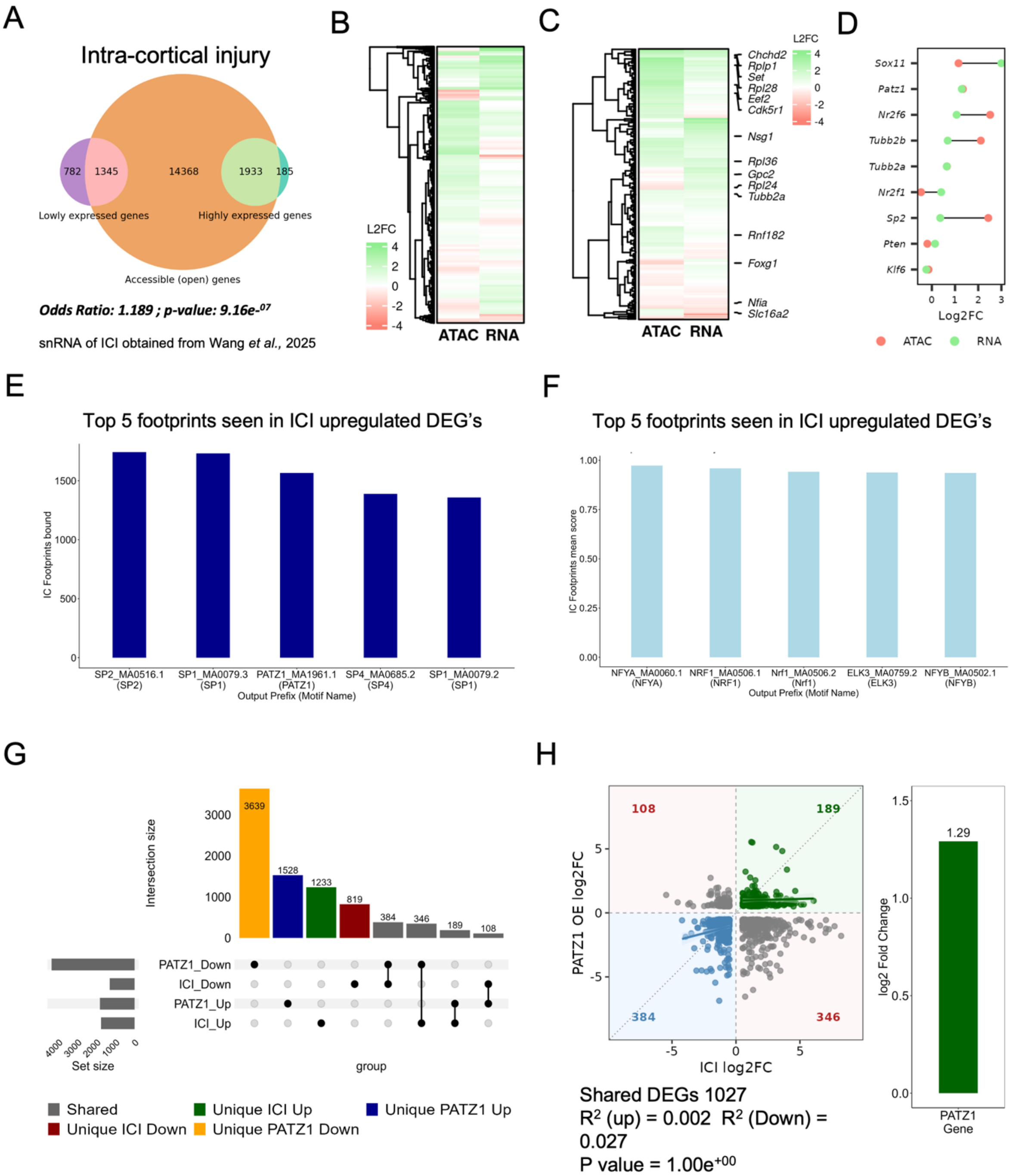


**Supplementary Figure S8. Integration of bulk ATAC-seq and single-nucleus RNA-seq following intra-cortical injury reveals concordant chromatin accessibility and transcriptional changes at pro-growth loci, with PATZ1 as a key shared transcriptional regulator.**

**(A)** Venn diagram showing the overlap between genes with accessible chromatin (open peaks) identified by bulk ATAC-seq and genes detected by single-nucleus RNA-seq (snRNA-seq) following intra-cortical injury (ICI). The three circles represent: Lowly expressed genes (purple, n=782 unique; 1,345 overlapping with accessible genes), Accessible (open) genes (orange, n=14,368 total), and Highly expressed genes (green, n=1,933 overlapping with accessible genes; 185 unique). The large overlap between accessible and highly expressed genes (1,933) confirms that chromatin opening broadly correlates with active transcription following ICI, while a subset of accessible loci (14,368) do not yet show high-level expression, suggesting that chromatin opening precedes or exceeds transcriptional activation at many loci. (Odds Ratio: 1.189 ; p-value: 9.16e-07)

**(B)** Hierarchically clustered heatmap showing paired Log2 fold-change (L2FC) values for ATAC-seq chromatin accessibility (left column) and snRNA-seq gene expression (right column) across all genes following ICI. Each row represents one gene; columns represent the two modalities. Colour scale indicates L2FC (dark green = strongly increased; dark red = strongly decreased; range –4 to +4). The broadly concordant green signal across both ATAC and RNA columns indicates genome-wide co-upregulation of chromatin accessibility and transcription following ICI.

**(C)** Paired heatmap as in (B), restricted to pro-growth genes. Labelled genes on the right include *Chchd2*, *Rplp1*, *Set*, *Rpl28*, *Eef2*, *Cdk5r1*, *Nsg1*, *Rpl36*, *Gpc2*, *Rpl24*, *Tubb2a*, *Rnf182*, *Foxg1*, *Nfia*, and *Slc16a2*. Pro-growth genes show a predominant pattern of increased ATAC accessibility (green) that is broadly mirrored by increased RNA expression, confirming that chromatin opening at these loci is functionally associated with transcriptional activation. A subset of genes (e.g., *Rnf182*, *Foxg1*, *Nfia*, *Slc16a2*) show discordant or reduced signal in one or both modalities, representing exceptions to the general trend.

**(D)** Dot plot showing Log2FC of ATAC-seq accessibility (red dots) and snRNA-seq expression (green dots) for nine selected genes following ICI: *Sox11*, *Patz1*, *Nr2f6*, *Tubb2b*, *Tubb2a*, *Nr2f1*, *Sp2*, *Pten*, and *Klf6*. The x-axis shows Log2FC (range 0–3). For most genes, ATAC Log2FC (red) is higher than RNA Log2FC (green), consistent with chromatin opening preceding or exceeding transcriptional upregulation. *Pten* and *Klf6* show near-equal and low values in both modalities, serving as controls. *Sox11* and *Nr2f6* show the largest ATAC-RNA discordance, suggesting post-transcriptional regulatory mechanisms may be at play.

**(E)** Bar graph showing the top 5 TF motifs ranked by the number of ICI footprints bound (y-axis: IC Footprints bound) within the accessible chromatin regions of ICI-upregulated differentially expressed genes (DEGs). The x-axis displays the motif identifier and TF name. Ranked from highest to lowest: SP2 (MA0516.1, ~1,750), SP1 (MA0079.3, ~1,750), PATZ1 (MA1961.1, ~1,575), SP4 (MA0685.2, ~1,400), and SP1 (MA0079.2, ~1,350). The presence of PATZ1 among the top 5 most-bound TFs at ICI-upregulated DEG loci further supports its role as a key transcriptional regulator of the injury-induced pro-growth chromatin response.

**(F)** Bar graph showing the top 5 TF motifs ranked by mean ICI footprint score (y-axis: IC Footprints mean score, range 0–1.00) within accessible chromatin regions of ICI-upregulated DEGs. Ranked from highest to lowest mean score: NFYA (MA0060.1, ~0.97), NRF1 (MA0506.1, ~0.95), Nrf1 (MA0506.2, ~0.93), ELK3 (MA0759.2, ~0.93), and NFYB (MA0502.1, ~0.92). These TFs show near-maximal footprint occupancy scores at ICI-upregulated DEG loci, indicating robust and consistent TF binding at these accessible regions. Together with panel E, these results identify both the most prevalent (E) and the most strongly occupied (F) TF binding events at transcriptionally active pro-growth loci following ICI.

**(G)** UpSet plot showing the intersection of differentially expressed genes (DEGs; $p < 0.05$, Log2FC > 0.5) between ICI snRNA-seq and PATZ1 overexpression snRNA-seq datasets. The horizontal bars (left, set size) show the total number of DEGs per group: PATZ1_Down (~3,800), ICI_Down (~900), PATZ1_Up (~2,500), and ICI_Up (~1,200). The vertical bars (intersection size) show the number of genes in each intersection category, colour-coded as: Unique PATZ1_Down (orange, n=3,639), Unique PATZ1_Up (dark blue, n=1,528), Unique ICI_Up (dark green, n=1,233), Unique ICI_Down (dark red, n=819), Shared downregulated (grey, n=384), Shared opposing/other (grey, n=346), Shared ICI_Up + PATZ1_Up (n=189), and Shared ICI_Down + PATZ1_Up or other combinations (n=108). The substantial unique fractions in each condition indicate partially distinct transcriptional programmes, while the shared upregulated intersection (n=189) represents genes co-induced by both ICI and PATZ1 overexpression.

**(H)** *(Left)* Scatter plot showing Log2FC concordance between ICI snRNA-seq (x-axis) and PATZ1 overexpression snRNA-seq (y-axis) for the 1,027 shared DEGs. Each dot represents one gene, colour-coded by category: Shared-up (green, n=189, upper-right quadrant), Shared-down (blue, n=384, lower-left quadrant), and Opposing/Other (grey, n=108 upper-left and n=346 lower-right quadrants). Green shading indicates concordant direction; red shading indicates discordant direction. The inset box reports: Shared DEGs=1,027; $R^2$(up)=0.002; $R^2$(down)=0.027; Fischer's test p=1.00e+00. The majority of shared DEGs (55.8%) are regulated in the same direction by both ICI and PATZ1 overexpression, supporting the conclusion that PATZ1 partially recapitulates the transcriptional response to intra-cortical injury. *(Right)* Bar graph showing the Log2FC of *PATZ1* itself in the ICI snRNA-seq dataset (Log2FC = 1.29), confirming that PATZ1 is endogenously upregulated at the transcriptional level following intra-cortical injury.

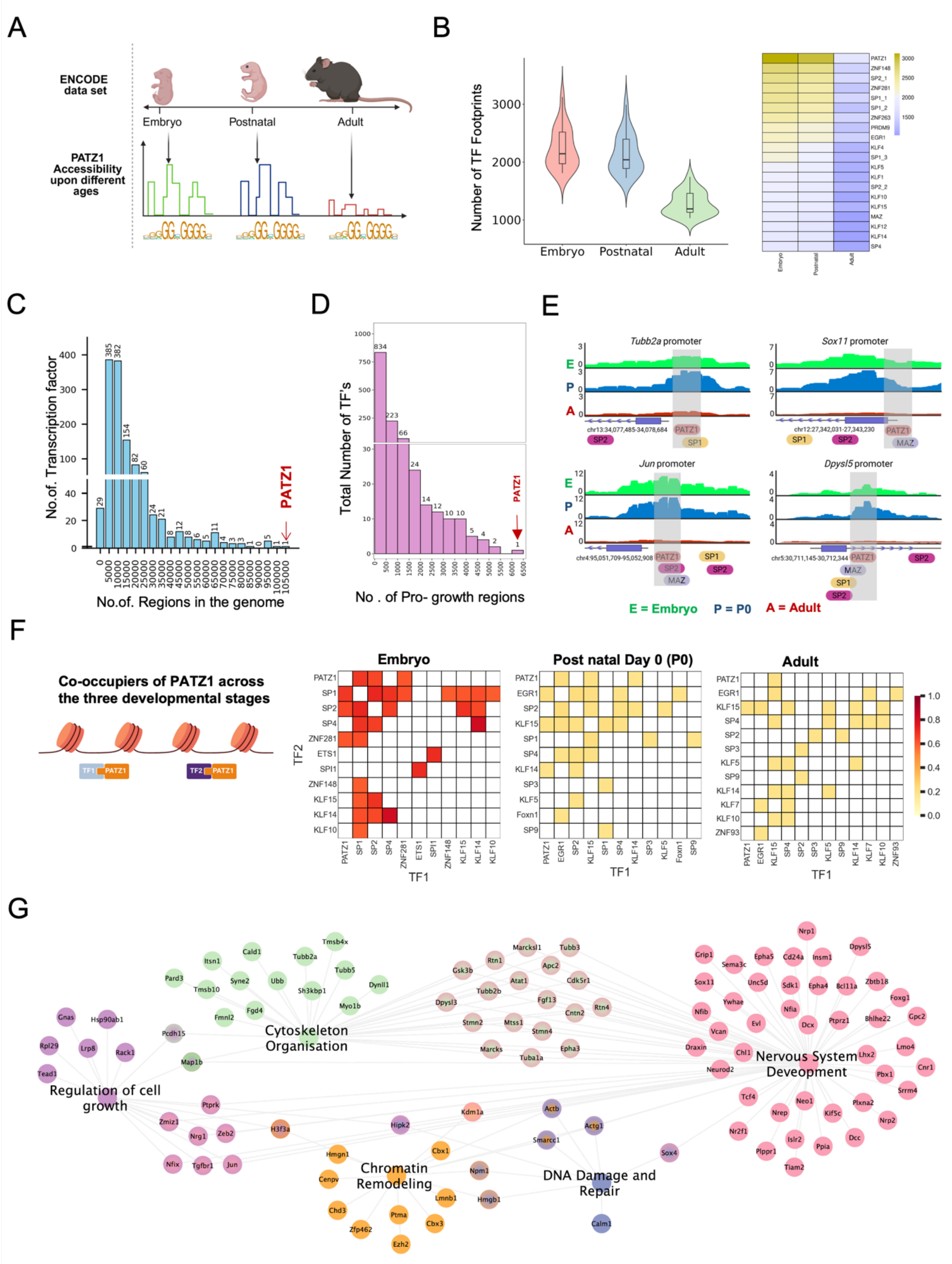


**Supplementary Figure S9: Dynamics of PATZ1 footprinting and transcription factor co-occupancy at pro-growth genes**.

**(A)** Illustration of transcription factor footprinting analysis methodology.

**(B)** Left panel: Violin plots depicting the distribution of transcription factor footprints across embryonic, postnatal, and adult stages, demonstrating progressive reduction in transcription factor (TF) binding sites in the adult. Right panel: Heatmap of key transcription factor footprinting across the three stages (embryo, postnatal, adult), where colour intensity indicates footprint count (lighter colours represent a higher number of footprints).

**(C)** Distribution of PATZ1 footprints across all accessible chromatin regions genome-wide. Histogram showing the frequency distribution of PATZ1 binding across all genomic regions (x-axis: number of regions in the genome; y-axis: number of transcription factors), demonstrating that PATZ1 occupies approximately 100,000 accessible regions genome-wide, representing one of the most broadly distributed transcription factors in the dataset

**(D)** Histogram showing the distribution of PATZ1 binding specifically within pro-growth regulatory regions (x-axis: number of pro-growth regions; y-axis: total number of transcription factors), with PATZ1 indicated by the red arrow, showing its relative frequency among transcription factors bound at pro-growth loci.

**(E)** Right panel: Representative UCSC Genome Browser tracks showing PATZ1 co-occupancy with SP1, SP2, and MAZ at four pro-growth gene promoters — Tubb2a (chr13:34,077,485–34,078,684), Sox11 (chr12:27,342,031–27,343,230), Jun (chr4:95,051,709–95,052,908), and Dpysl5 (chr5:30,711,145–30,712,344). Green tracks represent Embryo, blue tracks represent P0, and the red track represents adult conditions. Grey shaded regions highlight PATZ1 binding sites. Transcription factor motifs co-occurring at each locus are indicated below each track.

**(F)** Co-occupancy of PATZ1 with partner transcription factors across embryonic, postnatal, and adult developmental stages. Left: Schematic illustration of transcription factor co-binding at regulatory elements, showing PATZ1 co-occupying sites with partner TFs (TF1 and TF2). Right: Cosine similarity heatmaps showing pairwise co-occupancy scores between PATZ1 and its co-binding partners at accessible chromatin regions across three developmental stages — Embryonic (left), Postnatal (middle), and Adult (right). Colour intensity reflects cosine similarity score (dark red = high co-occupancy, range 0 to 1.0; yellow/white = low co-occupancy). In the embryonic stage, PATZ1 shows strong co-occupancy with SP1, SP2, SP4, ZNF281, ETS1, SPI1, ZNF148, KLF15, KLF14, and KLF10. Co-occupancy patterns are progressively reduced across postnatal and adult stages, consistent with the developmental decline of PATZ1 footprints observed in panel B.

**(G)** Network analysis of PATZ1 motif-containing regions overlapping with curated pro-growth gene regulatory elements (targeted analysis; see Methods). Genes are grouped into five major functional clusters: regulation of cell growth (purple), cytoskeleton organisation (green), chromatin remodelling (orange), DNA damage and repair (blue), and nervous system development (pink). Nodes represent individual genes, and edges indicate functional connections between genes. This targeted representation was chosen to specifically assess PATZ1 occupancy at growth-associated loci rather than as an unbiased genome-wide binding profile

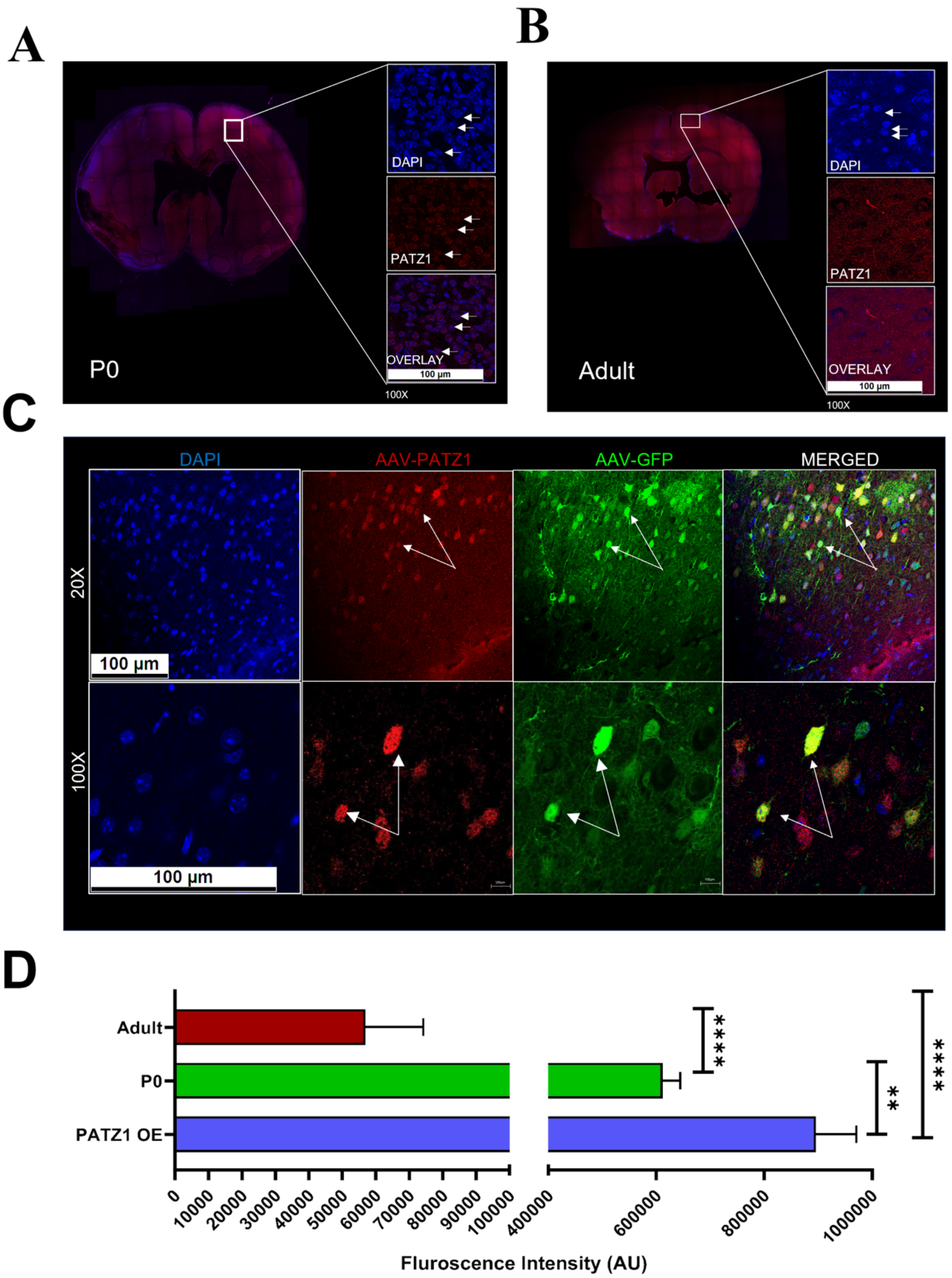


**Supplemental Figure S10: PATZ1 Expression in Developing and Adult Mouse Brain**

**(A)** Coronal section of P0 (postnatal day 0) mouse brain showing PATZ1 expression. Right panels (100X magnification): DAPI nuclear staining (blue, top), PATZ1 immunolabeling (red, middle), and overlay of both channels (bottom). White arrows indicate PATZ1-positive cells showing nuclear localization.

**(B)** Coronal section of adult mouse brain demonstrating baseline PATZ1 expression. Right panels (100X magnification): DAPI nuclear staining (blue, top), PATZ1 immunolabeling (red, middle), and overlay (bottom).

**(C)** Colocalization of PATZ1 and GFP in the transgenic mouse brain. Upper panels (40X magnification): DAPI nuclear staining (blue, first panel), PATZ1 immunolabeling (red, second panel), GFP expression (green, third panel), and merged channels (fourth panel). Lower panels (100X magnification): Higher magnification of the same region showing individual cells. White arrows indicate cells with colocalization of PATZ1 and GFP signals (yellow colour), demonstrating expression in the same cell population.

**(D)** Quantification of PATZ1 fluorescence intensity (in arbitrary units, AU) across different developmental stages and experimental conditions. The graph shows significantly higher PATZ1 expression in PATZ1 overexpression (OE) samples (blue bar, ~900,000 AU) compared to both P0 (green bar, ~600,000 AU; **$p<0.01$) and adult brain samples (red bar, ~500,000 AU; ****$p<0.0001$). Developmental downregulation of PATZ1 is evident in the significant difference between P0 and adult brain samples (****$p<0.0001$). All comparisons were performed using unpaired two-tailed t-tests (n=11 per group). Error bars represent SEM. Statistical significance is indicated as: **$p<0.01$, ****$p<0.0001$.

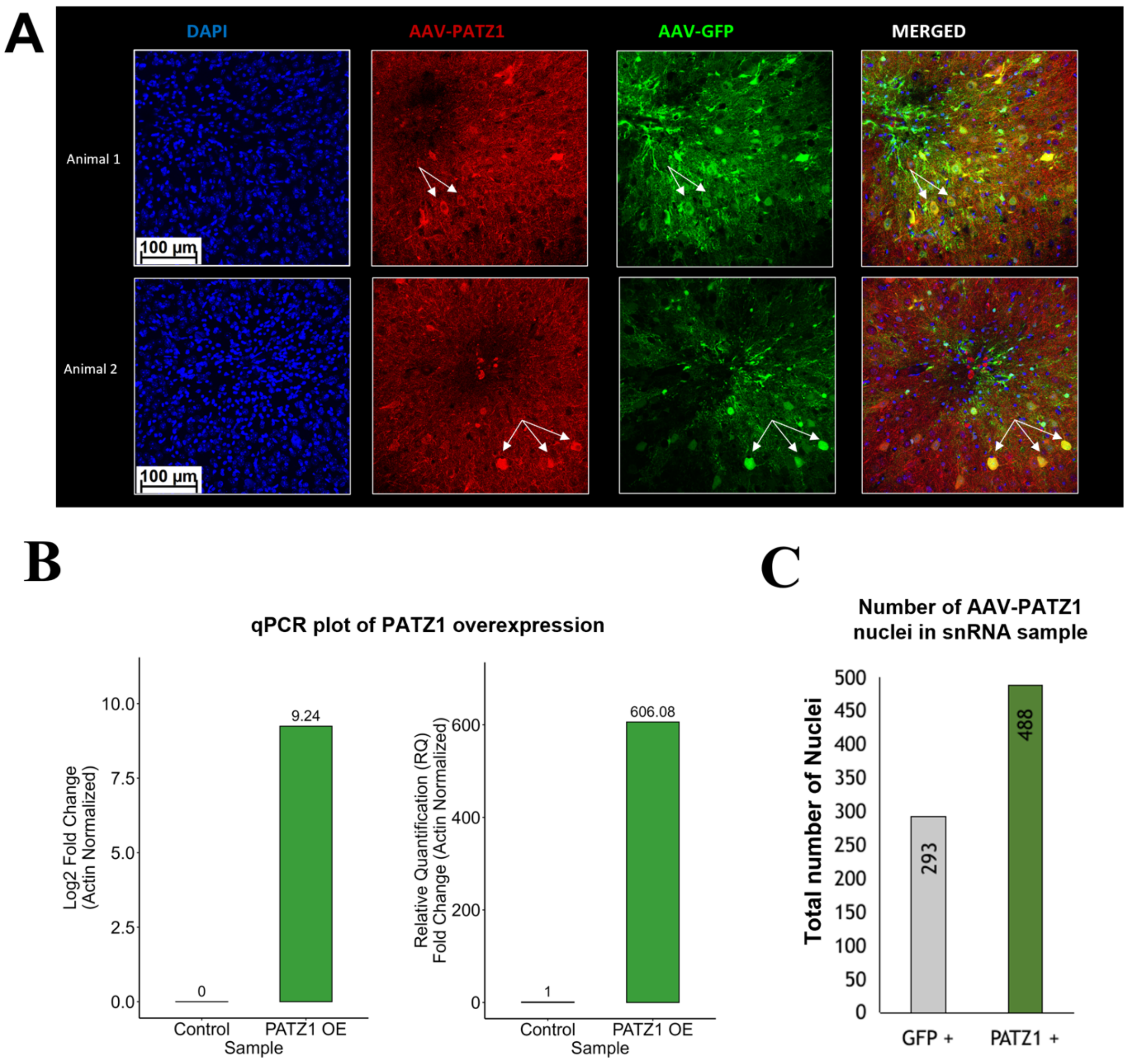


**Supplementary Figure S11 : AAV-PATZ1 transduction efficiency and viral titer quantification in vivo.**

**(A)** Representative immunofluorescence images of motor cortex sections from two animals injected with AAV-PATZ1 and AAV-GFP. DAPI (blue) labels cell nuclei. AAV-PATZ1 (red) indicates against PATZ1 over expression, and AAV-GFP (green) shows reporter gene expression. Merged images demonstrate co-localisation of AAV-PATZ1 and AAV-GFP signals (yellow/orange); white arrows indicate representative co-transduced cells. Scale bar = 100 µm.

**(B)** Quantitative RT-qPCR analysis of PATZ1 mRNA expression in PATZ1-overexpressing (PATZ1 OE) animals relative to control animals. Expression was calculated using the 2−ΔΔCt method, normalised to β-actin (Actb). Left panel: Log$_2$ fold change; right panel: relative quantification (RQ).

**(C)** Bar plot showing the total number of nuclei assigned to each AAV population (GFP+ and PATZ1+) identified in snRNA-seq Seurat clusters from co-injected animals. AAV identity was determined by detection of GFP or PATZ1 transgene transcripts in the single-nucleus dataset. Animals received co-injection of AAV-GFP and AAV-PATZ1.

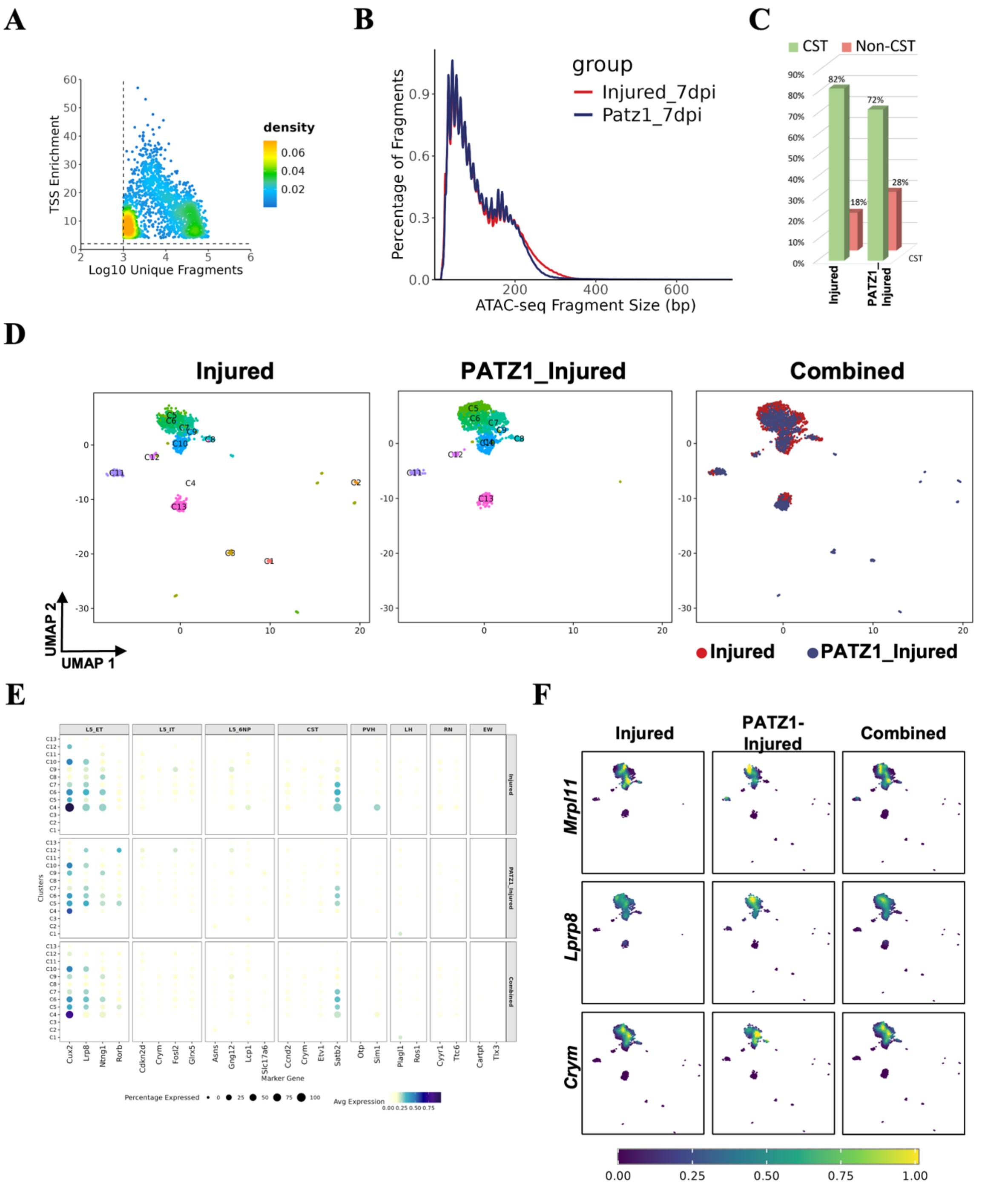


**Supplementary Figure S12: Extended single-nucleus ATAC-seq analysis supporting Figure 5, revealing chromatin accessibility landscape and cell type composition in Injured and PATZ1_Injured spinal cord.**

**(A)** snATAC-seq library quality metrics showing TSS enrichment score versus log10 unique fragments per cell. Density gradient (yellow = high, blue = low) confirms high-quality nuclei passing quality thresholds for downstream analysis.

**(B)** ATAC-seq fragment size distribution for Injured (red) and PATZ1_Injured (blue) samples, demonstrating characteristic nucleosomal periodicity (mono-, di-, and tri-nucleosomal banding pattern), confirming successful snATAC-seq library preparation.

**(C)** Bar graph quantifying the proportion of CST (green) and Non-CST (red) cell populations in Injured and PATZ1_Injured conditions, showing 82% vs 18% and 72% vs 28%, respectively.

**(D)** UMAP visualization of chromatin accessibility profiles from Injured (left), PATZ1_Injured (middle), and Combined (right) datasets. Clusters are color-coded by cell identity and labeled (C0–C15), enabling direct comparison of cluster distribution across conditions.

**(E)** Dot plot displaying cell type-specific chromatin accessibility at marker gene loci across identified clusters in PATZ1_Injured, Injured_7DPI, and Combined conditions. Dot size reflects the percentage of cells with accessible chromatin at each locus; color intensity represents average accessibility (light = low, dark blue = high). Marker genes are grouped by cell lineage: LS_ET, LS_IT, LS_SNP, CST, PSN, LH, RN, and EW populations. (LS_ET — Layer 5 Extra-Telencephalic (corticospinal/corticobulbar projection neurons), LS_IT — Layer 5 Intra-Telencephalic (corticocortical/corticostriatal projection neurons) LS_SNP — Layer 5/6 Near-Projecting neurons, CST — Corticospinal Tract neurons, PVH — Paraventricular Hypothalamic neurons, LH — Lateral Hypothalamic neurons, RN — Red Nucleus neurons, EW — Edinger-Westphal nucleus neurons)

**(F)** Feature plots showing normalized chromatin accessibility of CST marker genes *Mrpl11*, *Lrp8*, and *Crym* across UMAP embeddings in Injured, PATZ1_Injured, and Combined conditions. Color scale ranges from low (dark purple) to high (yellow) accessibility, highlighting CST-specific chromatin remodeling in response to PATZ1 overexpression.

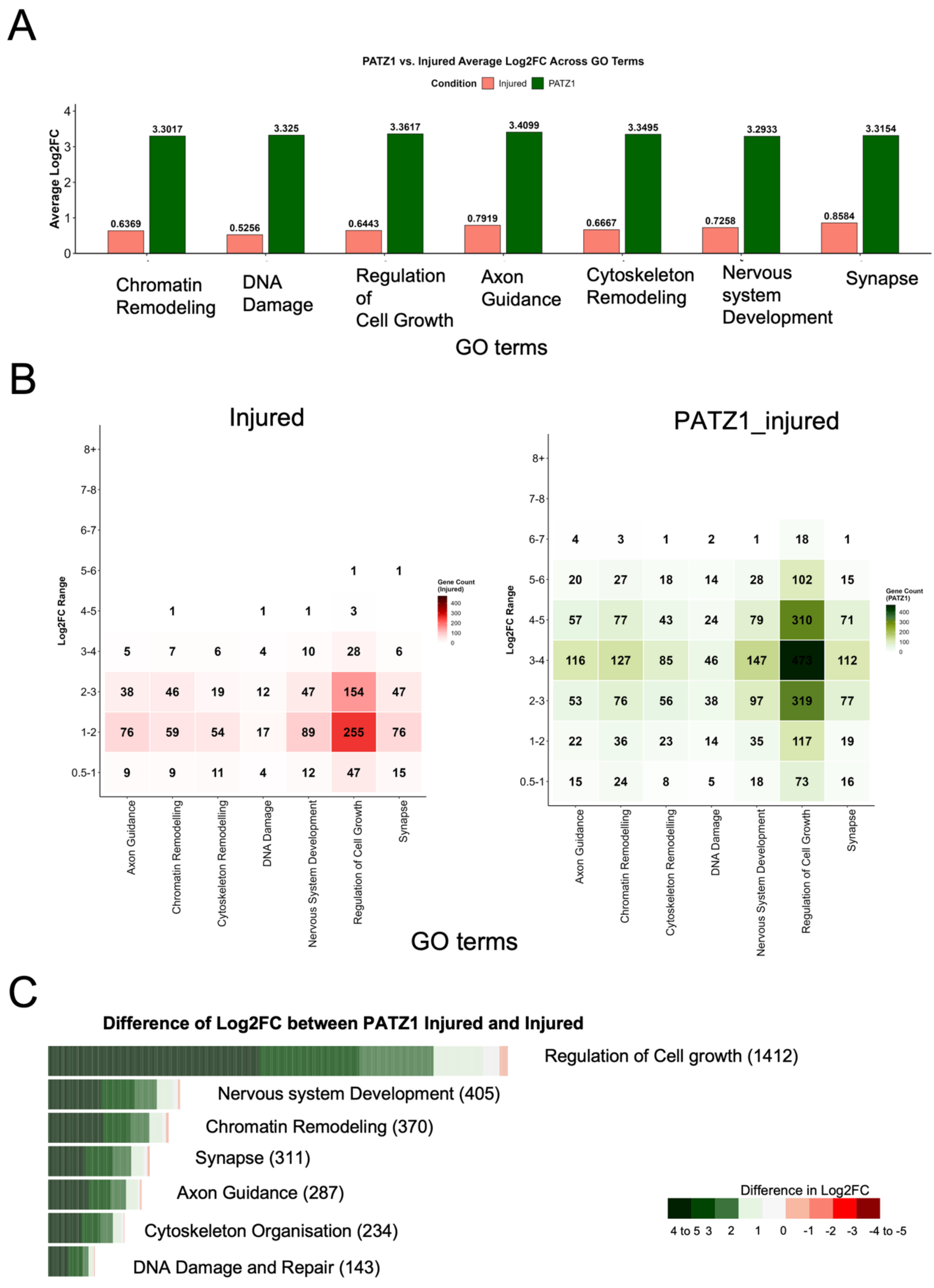


**Supplementary Figure S13. Alternative visualisations of snATAC-seq chromatin accessibility data comparing PATZ1-Injured and Injured control L5 ET neurons.**

**(A)** Grouped bar graph comparing mean chromatin accessibility Log2FC between PATZ1-Injured (dark green) and Injured control (salmon/red) neurons across seven GO-term-defined biological pathways: Chromatin Remodelling, DNA Damage, Regulation of Cell

Growth, Axon Guidance, Cytoskeleton Remodelling, Nervous System Development, and Synapse. Each pair of bars represents one GO term; exact mean Log2FC values are indicated above each bar. Across all pathways, PATZ1-Injured neurons display substantially higher mean Log2FC values (~3.3–3.4) compared to Injured controls (~0.5–0.9), consistent with genome-wide chromatin opening driven by PATZ1 overexpression. This plot serves as a linear alternative to the radar chart shown in Figure 5E.

**(B)** Paired heatmap showing the number of differentially accessible peaks binned by Log2FC magnitude for PATZ1-Injured (green gradient, left) and Injured control (red gradient, right) neurons across six GO-term-defined biological pathways. Rows represent Log2FC bins (0.5–1 at bottom to 8+ at top); columns represent GO term pathways. Colour intensity and numerical values within each tile reflect gene count, with darker colour indicating higher counts (colour scales shown right). Both heatmaps share the same gene count scale to allow direct comparison between conditions. PATZ1-Injured neurons show markedly higher gene counts across mid-to-high Log2FC bins (3–4 and above), particularly for Regulation of Cell Growth and Nervous System Development, reflecting the broader and stronger chromatin opening induced by PATZ1 overexpression. This plot serves as an alternative to the circular bar plots shown in Figure 5F.

**(C)** Diverging heatmap showing the per-gene difference in Log2FC between PATZ1-Injured and Injured control neurons across seven GO-term-defined biological pathways. Each horizontal bar represents an individual gene; bar length reflects the magnitude of the Log2FC difference, and colour encodes the direction and magnitude (dark green = PATZ1-Injured greater by 4–5 Log2FC units; dark red = Injured greater by 4–5 Log2FC units; white = no difference), as indicated in the shared colour scale (right). GO terms are ordered by total gene count (shown in parentheses): Regulation of Cell Growth (1,412), Nervous System Development (405), Chromatin Remodelling (370), Synapse (311), Axon Guidance (287), Cytoskeleton Organisation (234), and DNA Damage and Repair (143). The predominance of green across all pathways confirms that PATZ1 overexpression drives substantially greater chromatin accessibility genome-wide.

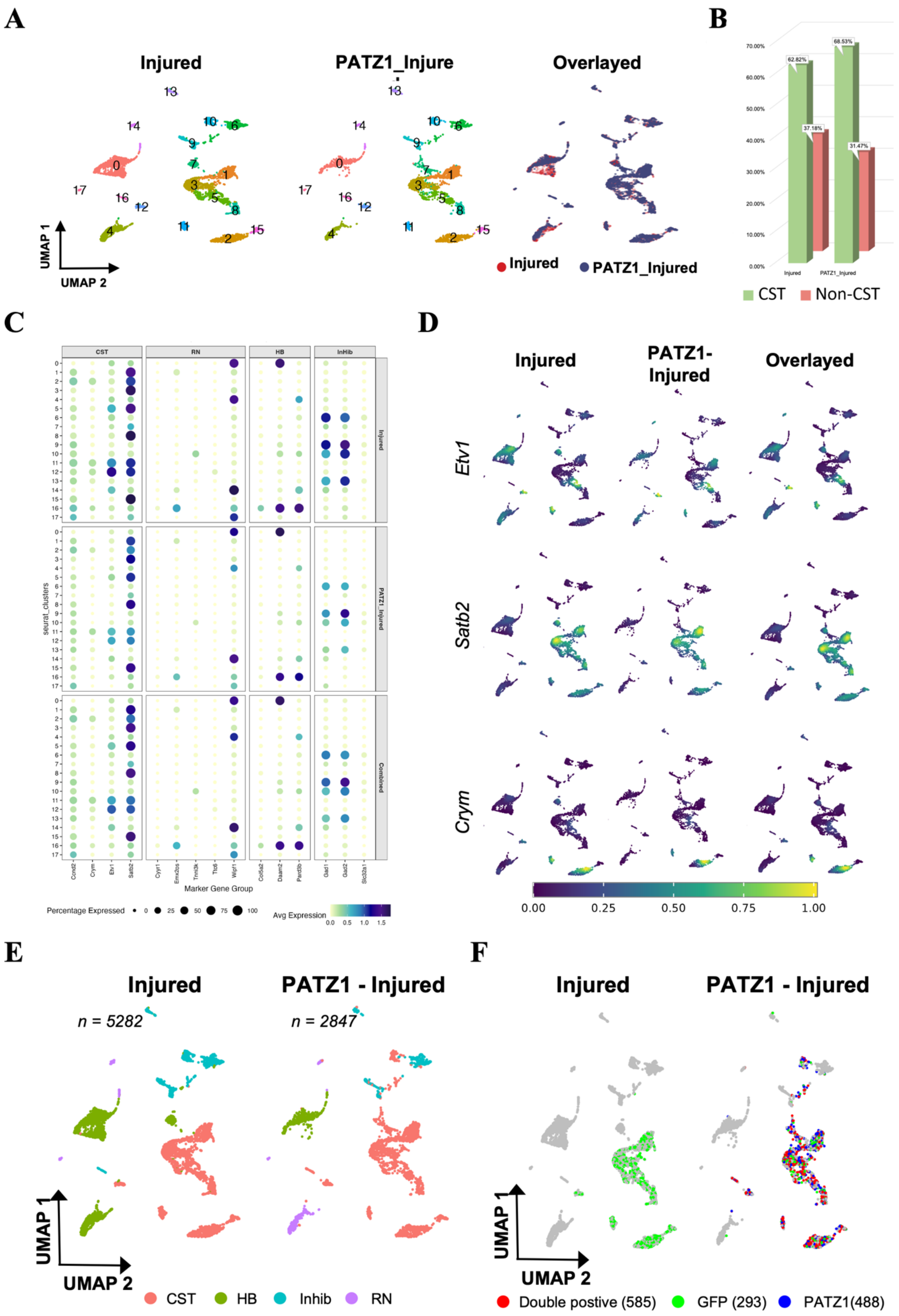
A
Injured
PATZ1_Injure
Overlayed
UMAP 1
UMAP 2
Injured
PATZ1_Injured
B
CST
Non-CST
C
CST
RN
HB
InHib
seurat_clusters
Marker Gene Group
Percentage Expressed
Avg Expression
D
Injured
PATZ1-Injured
Overlayed
Etv1
Satb2
Crym
E
Injured
PATZ1 - Injured
n = 5282
n = 2847
UMAP 1
UMAP 2
CST
HB
Inhib
RN
F
Injured
PATZ1 - Injured
UMAP 1
UMAP 2
Double postive (585)
GFP (293)
PATZ1(488)

**Supplementary Figure S14: Extended single-nucleus RNA-seq analysis supporting Figure 6, revealing detailed cellular composition, marker gene profiles, and PATZ1 overexpression effects in the injured spinal cord**

**(A)** UMAP visualization of snRNA-seq data showing cell clusters in Injured (left), PATZ1_Injured (middle), and overlayed (right) conditions. Individual clusters are numbered (0–18) and color-coded, enabling comparison of cluster distribution and composition between conditions.

**(B)** Bar graph comparing the proportion of CST (green) and Non-CST (red) neurons between Injured and PATZ1_Injured conditions. Percentages are indicated above each bar.

**(C)** Dot plot displaying expression of cell type-specific marker genes across Seurat clusters in Injured, PATZ1_Injured, and Combined conditions. Dot size reflects the percentage of cells expressing each marker gene; color intensity represents average expression level (light = low, dark blue = high). Marker genes are grouped by cell identity: CST, RN, HB, and Inhib populations (CST – Corticobrospinal Tract, RN= Red Nuclei, HB- Hind Brai, Inhib- Inhiboitory Neuron).

**(D)** Feature plots showing normalized expression of CST marker genes Etv1, Satb2, and Crym across UMAP embeddings in Injured, PATZ1_Injured, and Overlayed conditions. Color scale ranges from low (dark purple) to high (yellow) expression.

**(E)** UMAP plots for Injured and PATZ1_Injured conditions with cells colored by major cell type identity: CST (pink), HB (green), Inhib (teal), and RN (purple), illustrating the broad cellular composition in each condition.(F) UMAP plots highlighting the distribution of virally labeled cell populations in Injured and PATZ1_Injured conditions. Cells are colored by labeling identity: Double positive (red; n=1255), GFP (green; n=1255), and PATZ1 (blue; n=1046), with unlabeled cells shown in grey.

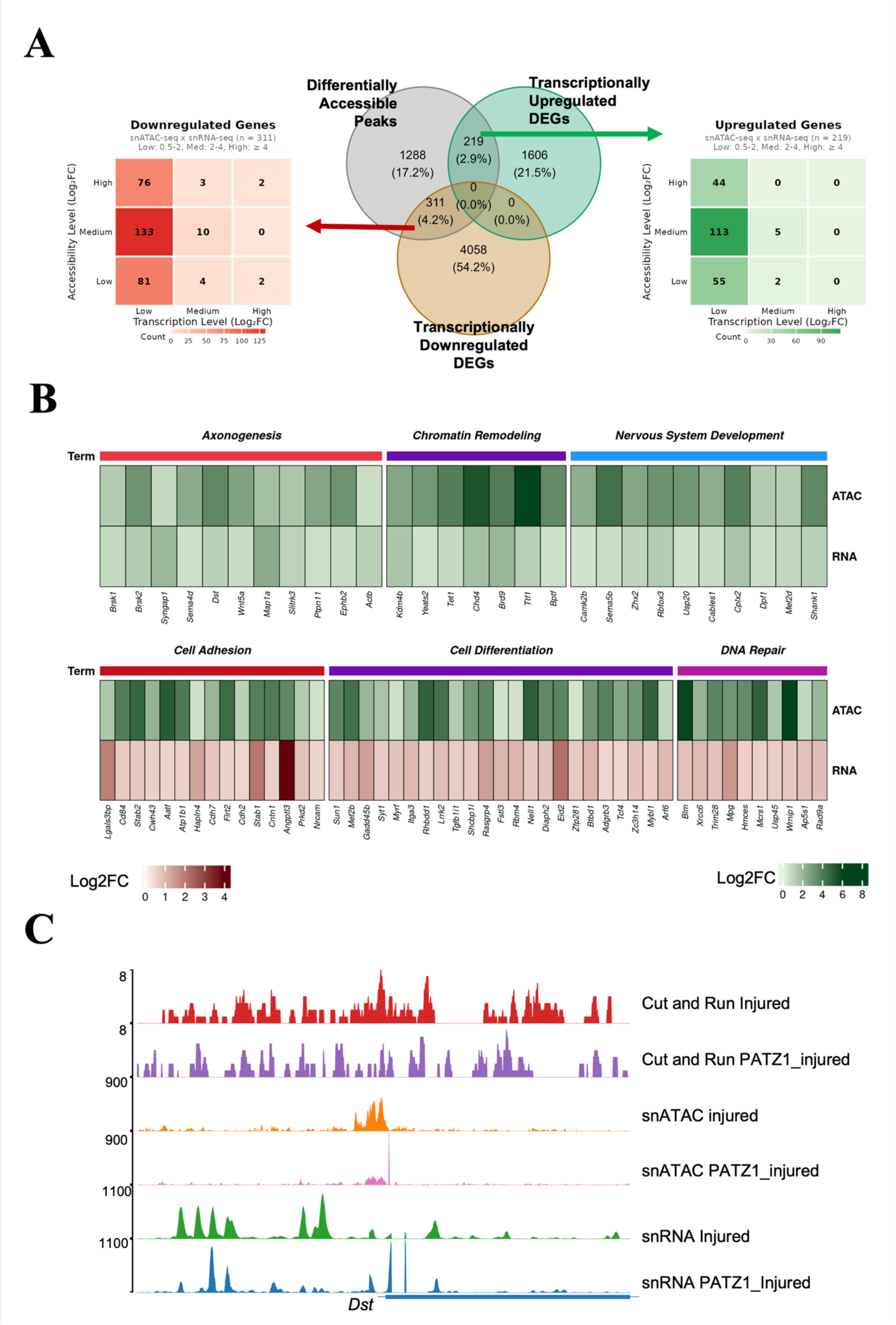

A
Differentially Accessible Peaks
Transcriptionally Upregulated DEGs
Transcriptionally Downregulated DEGs
1288 (17.2%)
219 (2.9%)
1606 (21.5%)
0 (0.0%)
311 (4.2%)
0 (0.0%)
4058 (54.2%)
Downregulated Genes
snATAC-seq x snRNA-seq (n = 311)
Low: 0.5-2, Med: 2-4, High: ≥ 4
Accessibility Level (Log₂FC)
High 76 3 2
Medium 133 10 0
Low 81 4 2
Low Medium High
Transcription Level (Log₂FC)
Count 0 25 50 75 100 125
Upregulated Genes
snATAC-seq x snRNA-seq (n = 219)
Low: 0.5-2, Med: 2-4, High: ≥ 4
Accessibility Level (Log₂FC)
High 44 0 0
Medium 113 5 0
Low 55 2 0
Low Medium High
Transcription Level (Log₂FC)
Count 0 30 60 90
B
Term
Axonogenesis
Chromatin Remodeling
Nervous System Development
ATAC
RNA
Brsk1 Brsk2 Syngap1 Sema4d Dst Wnt5a Map1a Slitrk3 Ptpn11 Ephb2 Actb
Kdm4b Yeats2 Tet1 Chd4 Brd9 Tti1 Bptf
Camk2b Sema5b Zhx2 Rblox3 Usp20 Cables1 Cplx2 Dpf1 Mef2d Shank1
Cell Adhesion
Cell Differentiation
DNA Repair
Lgals3bp Cd84 Stab2 Cwh43 Aatf Atp1b1 Hapln4 Cdh7 Flrt2 Cdh2 Stab1 Cntn1 Angptl3 Prkd2 Nrcam
Sun1 Mef2b Gadd45b Syt1 Myrf Itga3 Rhbdd1 Lrrk2 Tgfb1i1 Shcbp1l Rasgrp4 Fstl3 Rbm4 Nell1 Diaph2 Eid2 Zfp281 Btbd1 Adgrb3 Tcf4 Zc3h14 Mybl1 Arl6
Blm Xrcc6 Trim28 Mpg Hmces Mcrs1 Usp45 Wrnip1 Ap5s1 Rad9a
Log2FC
0 1 2 3 4
Log2FC
0 2 4 6 8
C
8
8
900
900
1100
1100
Cut and Run Injured
Cut and Run PATZ1_injured
snATAC injured
snATAC PATZ1_injured
snRNA Injured
snRNA PATZ1_Injured
Dst

**Supplementary Figure S15: Extended integrative analysis of chromatin accessibility and transcriptional changes induced by PATZ1 overexpression, supporting Figure 6.**

**(A)** Venn diagram illustrating the overlap between differentially accessible chromatin peaks (snATAC-seq) and transcriptionally upregulated (top right) and downregulated (bottom) differentially expressed genes (DEGs) in PATZ1_Injured versus Injured conditions. Flanking heatmaps quantify the distribution of overlapping genes categorized by chromatin accessibility level (snATAC-seq log2FC, y-axis) and transcription level (snRNA-seq log2FC, x-axis), with color intensity representing gene count. The majority of differentially accessible peaks (54.2%) do not overlap with DEGs, while 2.9% show concordant upregulation and 4.2% show concordant downregulation.

**(B)** Heatmap displaying paired ATAC log2FC (top row, green) and RNA log2FC (bottom row, red-green) for genes showing concordant chromatin and transcriptional changes. Upper panel shows genes with increased chromatin accessibility and transcriptional upregulation, grouped by GO biological process terms: Axonogenesis, Nervous System Development, and Chromatin Remodeling. Lower panel shows genes with increased chromatin accessibility but transcriptional downregulation, grouped by GO terms: DNA Repair, Cell Differentiation, and Cell Adhesion. Color intensity reflects the magnitude of log2FC for ATAC (green scale, 1–5) and RNA (red scale, 1–3) changes respectively.

**(C)** Genome browser tracks displaying multi-modal genomic data at the *Dst* locus across six conditions: CUT&RUN Injured (red), CUT&RUN PATZ1 (purple), snATAC Injured (orange), snATAC PATZ1_Injured (pink), snRNA Injured (green), and snRNA PATZ1_Injured (blue). Tracks demonstrate concordant PATZ1 binding, chromatin accessibility, and transcriptional changes at this CST-enriched locus, illustrating the multi-layered regulatory effect of PATZ1 overexpression.

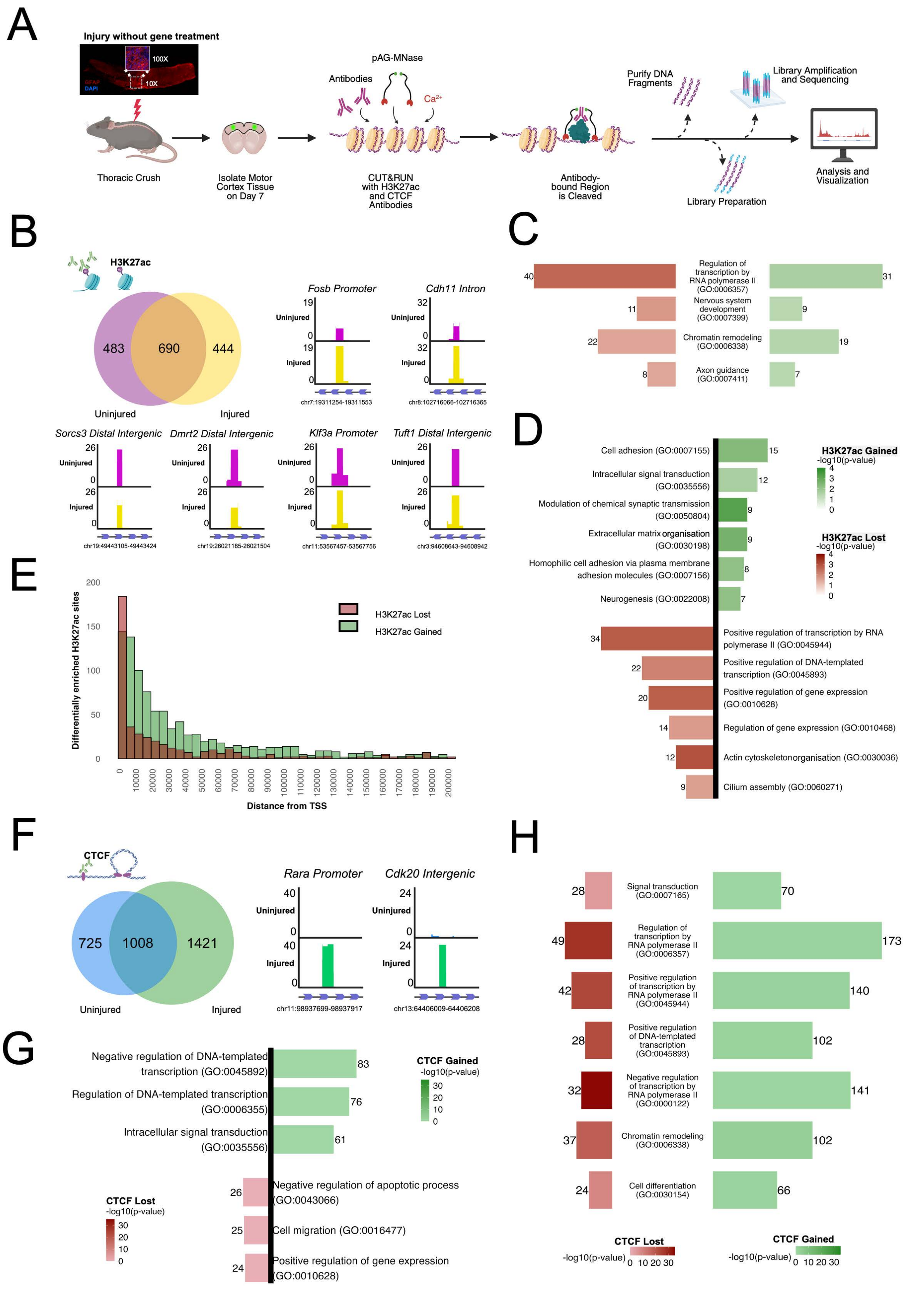


**Supplementary Figure S16: Genome-wide profiling of H3K27ac and CTCF binding using CUT&RUN.**

**(A)** Schematic overview of the CUT&RUN experimental strategy.

**(B)** Venn diagram comparing H3K27ac regions between uninjured (purple) and injured (yellow) conditions, showing 483 unique, 690 overlapping, and 444 unique regions, along with a representative genomic locus displayed using the UCSC genome browser.

**(C)** Gene Ontology (GO) pathways commonly enriched in H3K27ac-marked regions following injury, showing H3K27ac lost (red) and H3K27ac gained (green) categories.

**(D)** GO pathways uniquely enriched in H327ac regions in the injured, where unique H3k27ac is lost upon injury (red), and unique H3K27ac terms are gained upon Injury (green).

**(E)** Genomic distribution of distance of H3K27ac peaks relative to transcription start sites (TSS) between H3k27ac gained(green) and lost (red).

**(F)** Venn diagram comparing CTCF regions between uninjured (blue) and injured (green) conditions, showing 725 unique, 1008 overlapping, and 1421 unique regions with representative genome browser tracks.

**(G)** GO pathways uniquely enriched in CTCF binding sites upon injury. Left side shows pathways for CTCF Lost (red bars) Right side shows pathways for CTCF Gained (green bars).

**(H)** GO pathways commonly enriched in CTCF binding sites across both conditions. Horizontal bar chart of GO pathways commonly enriched in CTCF binding sites, with red bars (CTCF Lost) and green bars (CTCF Gained)

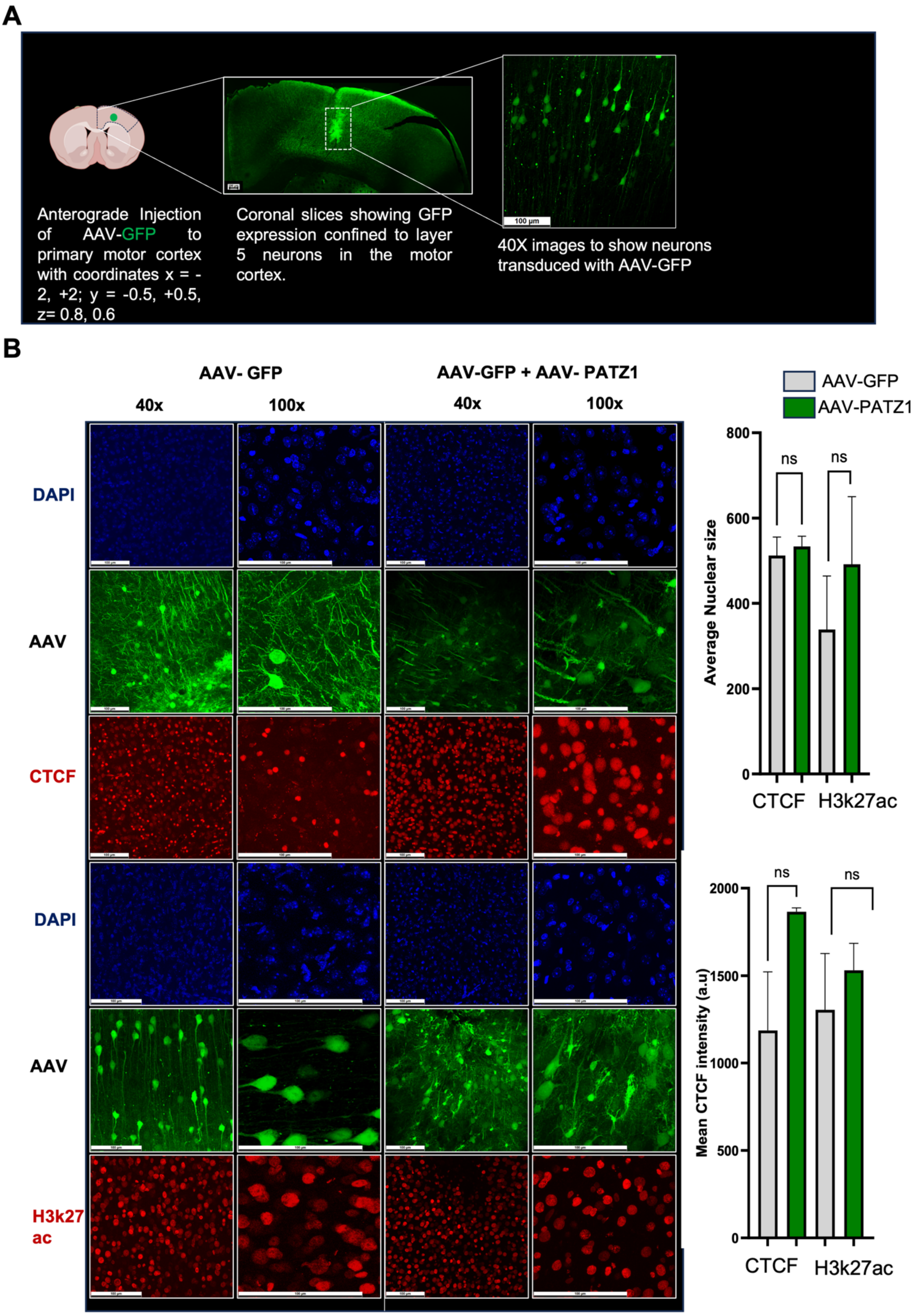


**Supplementary Figure S17: Validation of AAV-PAT21 delivery to motor cortex layer 5 and chromatin marker antibody specificity.**

**(A)** AAV-PATZ1-GFP injection targeting and transduction in layer 5 (L5) motor cortex neurons. Schematic illustrates the stereotaxic injection site (left) and representative image of GFP-positive transduced neurons within L5 (right).

**(B)** Validation of CTCF and H3K27ac antibody specificity by immunofluorescence. Upper panels: Representative immunofluorescence images showing DAPI (blue), AAV-GFP (green), and CTCF (red) staining in control (AAV-GFP) and PATZ1 overexpression (AAV-PATZ1) conditions at 40× and 100× magnification, confirming expected nuclear localisation of CTCF. Lower panels: Representative immunofluorescence images showing DAPI (blue), H3K27ac (red), and AAV-GFP or AAV-PATZ1 (green) in control and PATZ1 overexpression conditions at 40× and 100× magnification, confirming a nuclear H3K27ac distribution pattern consistent with active chromatin marking. **Right:** Quantification of nuclear size between AAV-GFP and AAV-PATZ1 conditions revealed no significant difference for either CTCF-stained samples ($p = 0.333$, two-tailed non-parametric Mann–Whitney U test, $U = 0$) or H3K27ac-stained samples ($p = 0.6667$, two-tailed non-parametric Mann–Whitney U test, $U = 1$). **Bottom:** Quantification of fluorescence intensity between AAV-GFP and AAV-PATZ1 conditions revealed no significant difference in either CTCF-stained samples ($p = 0.6667$, two-tailed non-parametric Mann–Whitney U test, $U = 1$) or H3K27ac-stained samples ($p = 0.6667$, two-tailed non-parametric Mann–Whitney U test, $U = 1$).

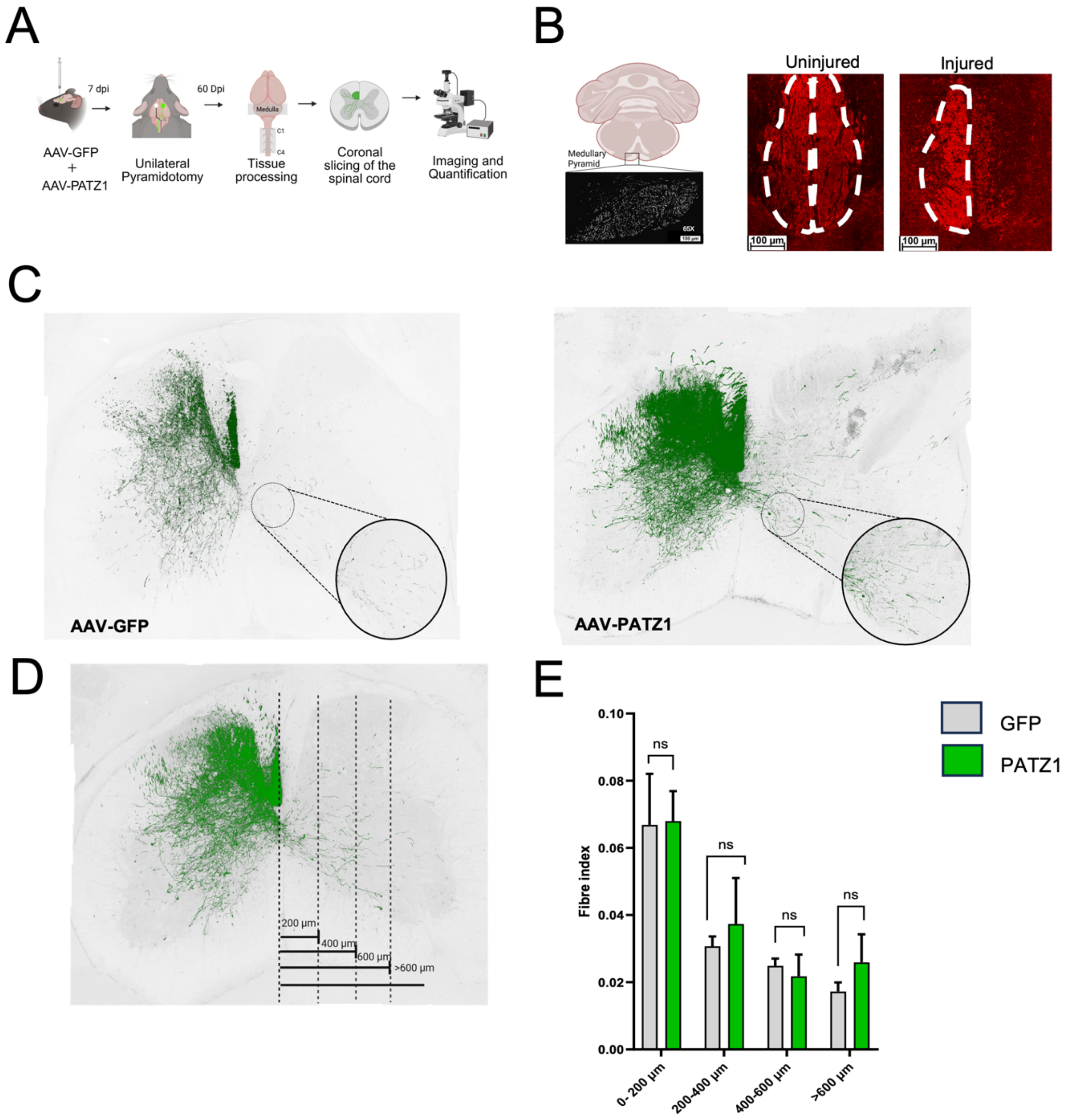


**Supplementary Figure S18. Pyramidotomy model reveals limited axonal sprouting in response to PATZ1 treatment**

**(A)** Experimental design showing AAV injection, pyramidotomy injury, and analysis timeline.

**(B)** Left: CST labelling is confirmed with medullary pyramids. Right: Pyramidotomy is confirmed with PKCγ IHC (dashed outlines); uninjured tissue shows both pyramids, and injured tissue shows only one side pyramid.

**(C)** GFP-labelled corticospinal axons in AAV-GFP control and AAV-PATZ1-treated spinal cords after pyramidotomy. Insets show higher magnification of sprouting patterns. We inverted the original image and then adjusted the selective colour (green) in Photoshop.

**(D)** Distance measurement schematic showing analysis bins from the lesion border.

**(E)** Quantification of fibre index across distances from the lesion site. Bar graph showing fibre index (density of GFP+ axons normalised to total tissue area) in control (AAV-GFP, white bars) and PATZ1-treated (AAV-PATZ1, green bars) animals at increasing distances from the pyramidotomy lesion (0-200 µm, 200-400 µm, 400-600 µm, >600 µm). Two-way ANOVA revealed a significant main effect of distance (F(3,9) = 9.689, **P = 0.0035), indicating a progressive decrease in fibre density with increasing distance from the lesion. No significant main effect of PATZ1 (P = 0.4497) or interaction between PATZ1 and distance (P = 0.8538) was observed. Sidak's post-hoc comparisons showed no significant differences between groups at individual distances (all adjusted P > 0.5). Data are presented as mean ± SEM; n = 3-4 animals per group. **P < 0.01; n.s. = not significant.

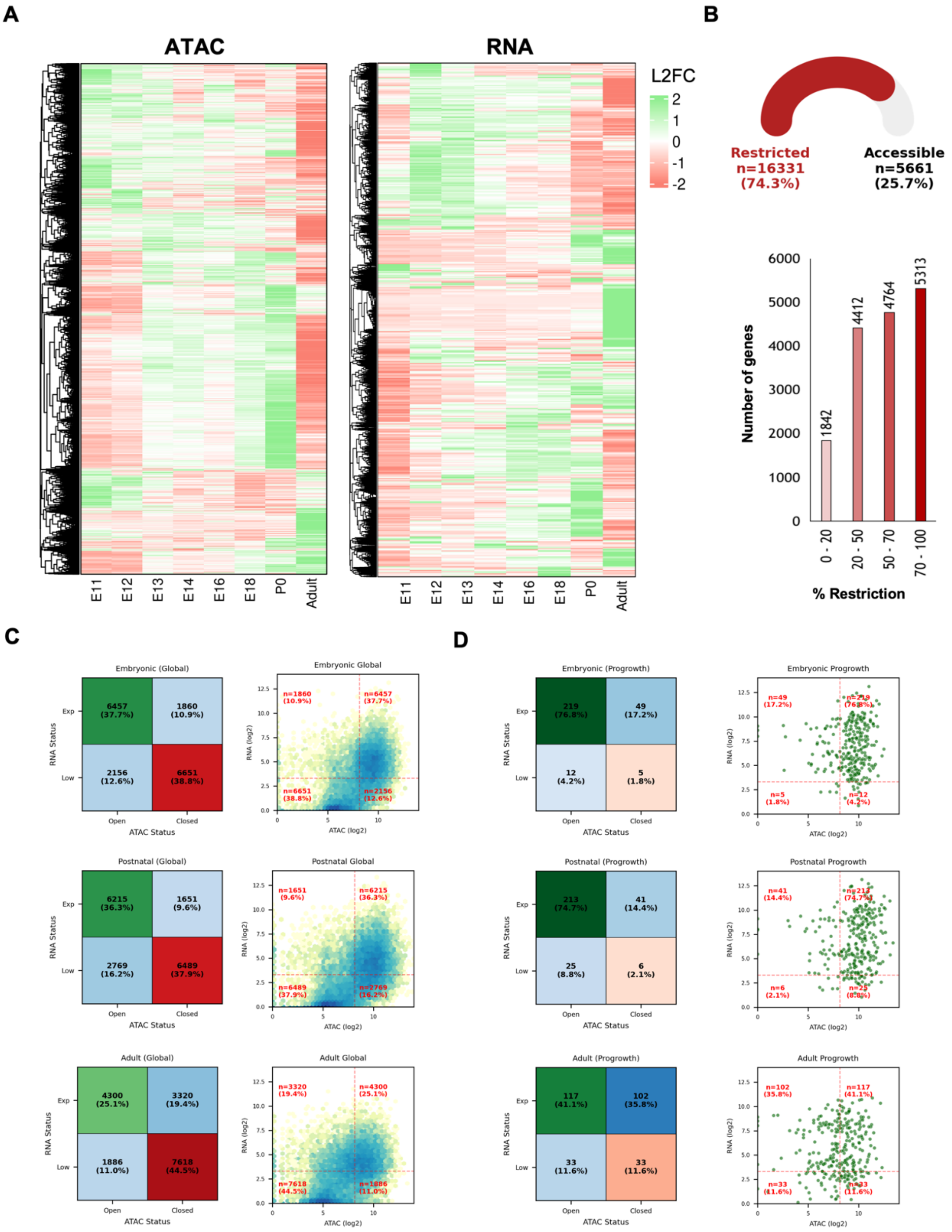


**Supplementary Figure S19: Chromatin Accessibility and Transcriptomic Landscapes Across Development and Injury.**

**(A)** Hierarchical clustering heatmap of ATAC-seq signals at gene promoter regions across developmental stages (E11, E12, E13, E14, E16, E18, P0) and the adult brain. Color scale represents log2 fold change (L2FC), where green indicates accessible (open) and red indicates inaccessible (closed) chromatin, revealing dynamic remodeling of the chromatin landscape during development.

**(B)** Top: Gauge plot representing the number of genes ( entire mouse genes, which are represented as global)  which are accessible and restricted between P0 and Adult. Red represents the total genes restricted (74.3%) , and grey represents accessible (25.7%). Bottom: Bar graph quantifying the number of genes undergoing chromatin restriction from the neonatal (P0) to adult stage, categorized by severity of accessibility loss (20–0%, 50–20%, 70–50%, 100–70%). Restriction was calculated as ((P0 − Adult) / P0) × 100, with the majority of genes showing greater than 70% reduction in chromatin accessibility.

**(C)** Confusion matrices (left) and scatter plots (right) correlating RNA-seq expression status (Expressed vs. Low) with ATAC-seq chromatin accessibility (Open vs. Closed) across global gene sets during Embryonic, Postnatal, and Adult stages. Each quadrant displays gene counts and percentages, highlighting the strong correspondence between open chromatin and active transcription across developmental stages.

**(D)** Confusion matrices (left) and scatter plots (right) showing the relationship between chromatin accessibility and transcriptional status specifically for pro-growth genes across Embryonic, Postnatal, and Adult stages. Compared to the global gene set in panel C, pro-growth genes show a progressively stronger bias toward open chromatin and active expression during embryonic stages, with this correspondence diminishing in the adult, reflecting developmental restriction of pro-growth gene accessibility.

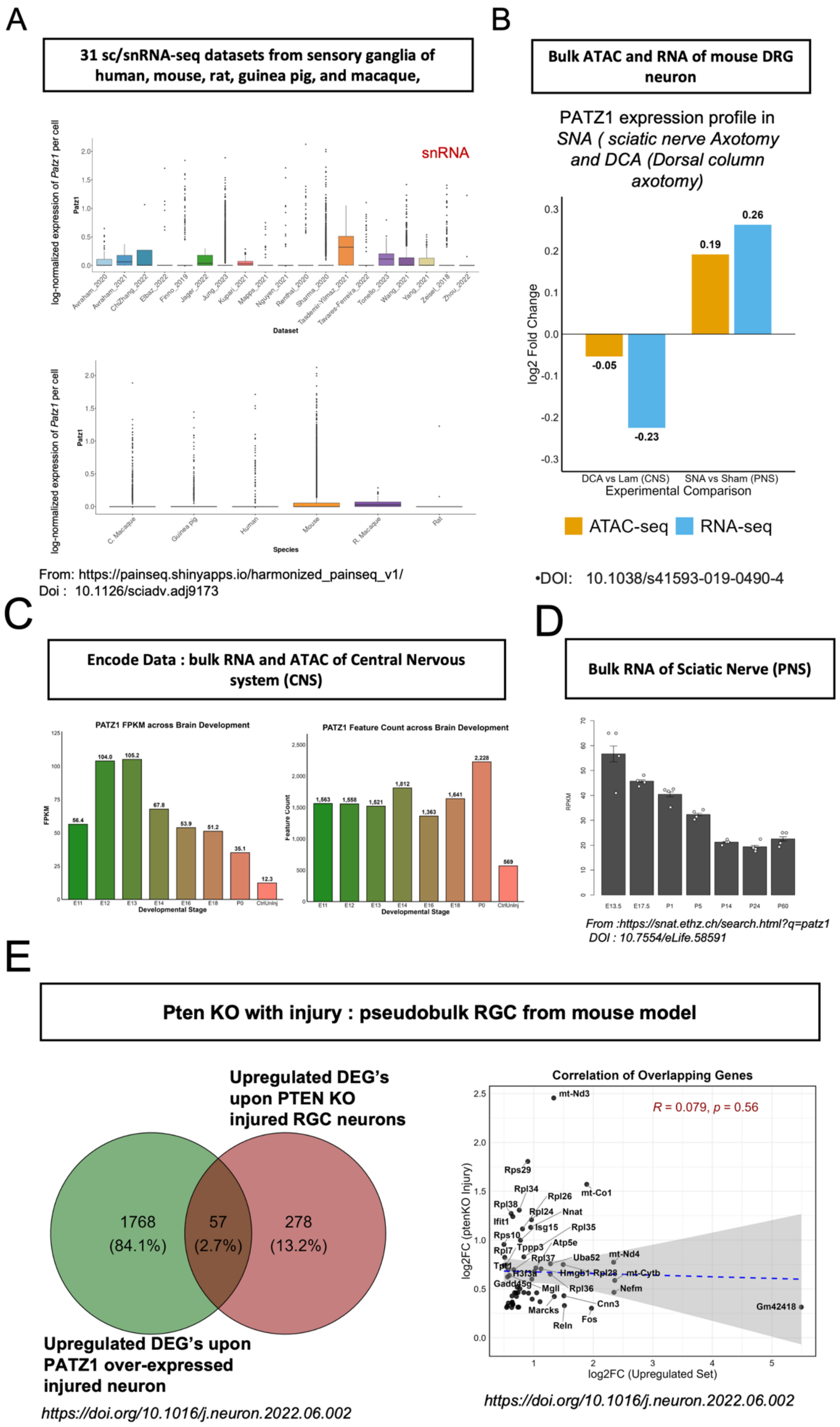


**Supplementary Figure S20: PATZ1 expression across species, tissue types, and injury contexts, and comparison with PTEN KO transcriptional programs.**

**(A)** Box plots showing log-normalized expression of *Patz1* per cell across 31 sc/snRNA-seq datasets derived from sensory ganglia of human, mouse, rat, guinea pig, and macaque (data sourced from the Harmonized Pain Sequencing resource). Upper panel displays expression stratified by individual dataset; lower panel summarizes expression grouped by species. Notably, *Patz1* expression is largely absent or minimal across sensory ganglia datasets, consistent with its restricted expression in CNS neurons.

**(B)** Bar graph showing log2 fold change (log2FC) of *Patz1* chromatin accessibility (ATAC-seq, orange) and transcript levels (RNA-seq, blue) in mouse dorsal root ganglion (DRG) neurons under two injury paradigms: dorsal column axotomy versus laminectomy (DCA vs Lam, CNS injury) and sciatic nerve axotomy versus sham (SNA vs Sham, PNS injury). *Patz1* shows increased accessibility and transcription following peripheral nerve injury (SNA), but not following CNS injury (DCA).

**(C)** Bar graphs showing *Patz1* expression levels across mouse brain developmental stages (E11–P0 and adult) from ENCODE bulk datasets. Left: FPKM values from bulk RNA-seq, showing high embryonic expression that declines postnatally. Right: ATAC-seq feature counts at the *Patz1* locus across the same stages, confirming concordant reduction in chromatin accessibility, consistent with developmental restriction of *Patz1* in the CNS.

**(D)** Bar graph showing *Patz1* mRNA expression (RPKM) in sciatic nerve at postnatal developmental timepoints (E13.5, E17.5, P1, P5, P14, P24, P60) from bulk RNA-seq data. Expression is highest at embryonic stages and progressively declines postnatally, suggesting a role for PATZ1 during peripheral nervous system development.

**(E)** Left: Venn diagram comparing DEGs upregulated upon *Patz1* overexpression in injured neurons (green; n=1,825) with DEGs upregulated in *Pten* knockout injured retinal ganglion cells (RGCs; red; n=335; Jacobi et al., 2022). A total of 57 genes (2.7%) are shared between the two datasets. Right: Scatter plot of log2FC values for the 57 overlapping genes between the PATZ1-upregulated set (x-axis) and the PTEN KO injury dataset (y-axis). The absence of significant correlation (Pearson R = 0.079, p = 0.56) indicates that despite partial gene overlap, PATZ1 overexpression and PTEN KO drive largely distinct transcriptional programs in injured neurons.

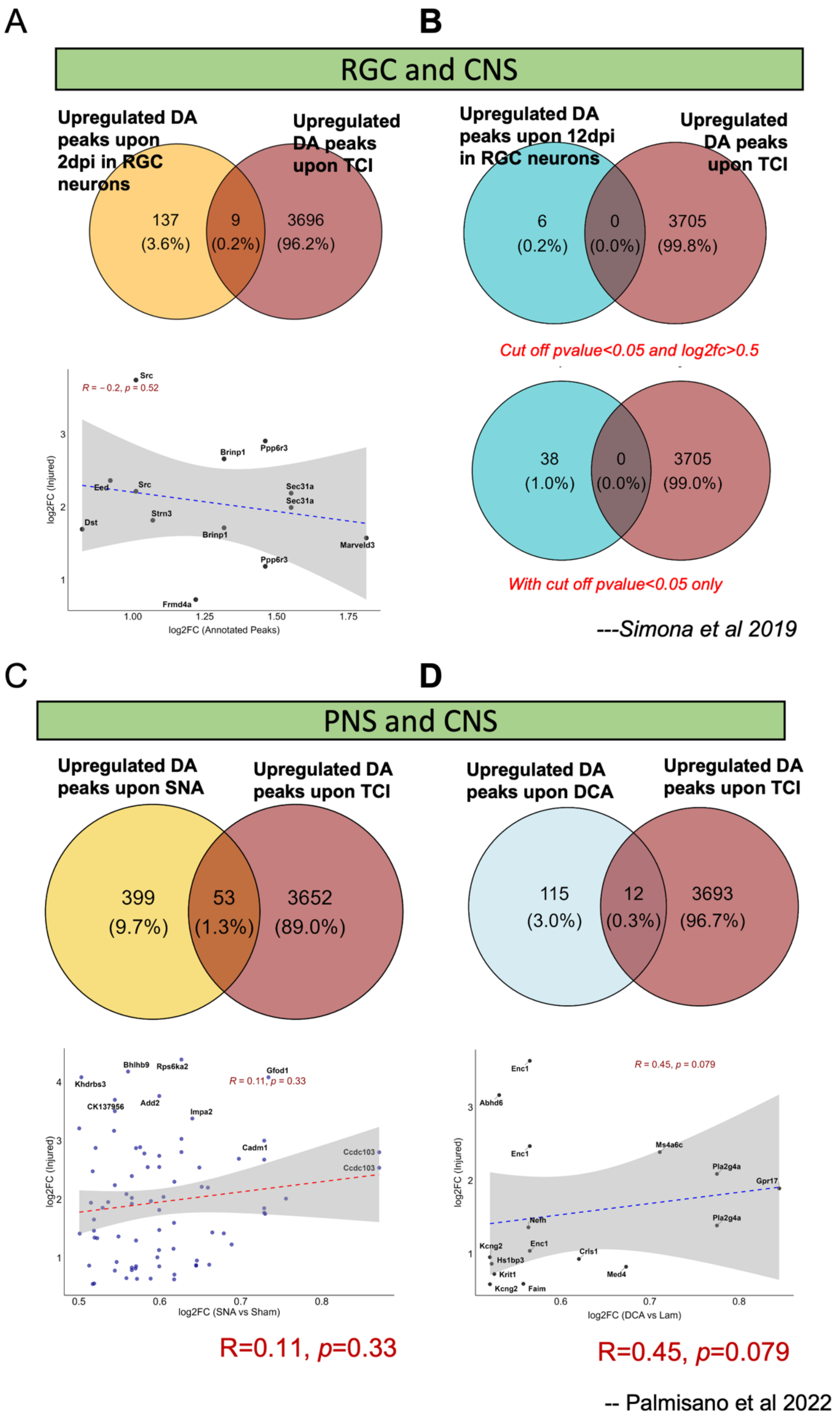


**Supplementary Figure S21: Cross-injury-model comparison of chromatin accessibility changes reveals limited overlap with spinal cord injury-induced chromatin remodeling in corticospinal neurons.**

**RGC and CNS comparison (Panels A–B; dataset from Simona et al., 2019)**

**(A)** TThe 5,482 TCI-upregulated accessibility peaks were first deduplicated by gene symbol (SYMBOL), yielding 3,705 unique genes, which were used for all Venn diagram comparisons below. Multiple peaks mapping to the same gene were collapsed to a single entry prior to analysis.Top: Venn diagram showing overlap between upregulated differentially accessible (DA) peaks in retinal ganglion cells (RGCs) at 2 days post-injury (2dpi; n=146) and Thoracic crush injury (TCI) -upregulated DA peaks in corticospinal neurons (n=3,705). Only 9 peaks (0.2%) are shared, with 137 (3.6%) unique to RGC 2dpi and 3,696 (96.2%) unique to SCI, indicating minimal chromatin accessibility overlap between these two injury models. Bottom: Scatter plot showing the correlation of log2FC values for the 9 shared peaks between RGC 2dpi (x-axis) and TCI (y-axis). Pearson correlation is minimal and non-significant ($R = -0.2$, $p = 0.52$), indicating no meaningful linear relationship between the magnitude of chromatin opening across the two injury contexts.

**(B)** Top: Venn diagram showing the same comparison at 12dpi in RGCs under stringent thresholds ($p < 0.05$ and log2FC > 0.5), revealing 0 shared peaks (0.0%), with 6 unique to RGC 12dpi (0.2%) and 3,705 unique to TCI (99.8%). Bottom: Applying a relaxed threshold ($p < 0.05$ only), the overlap remains negligible at 0 shared peaks (0.0%), with 38 unique to RGC (1.0%) and 3,705 unique to TCI (99.0%), collectively demonstrating that injury-induced chromatin remodeling in RGC neurons does not recapitulate thoracic crush injury-associated accessibility changes in corticospinal neurons.

***PNS and CNS comparison*** *(Panels C–D; dataset from Ilaria et al., 2022)*

**(C)** Top: Venn diagram comparing upregulated DA peaks following sciatic nerve axotomy (SNA; a regeneration-permissive peripheral nervous system injury model; n=452) with TCI (Thoracic crush Injury) -upregulated DA peaks (n=3,705). Only 53 peaks (1.3%) are shared, with 399 (9.7%) unique to SNA and 3,652 (89.0%) unique to TCI. Bottom: Scatter plot of log2FC values for the 53 shared peaks in SNA vs Sham (x-axis) versus TCI-Injured (y-axis). Pearson correlation is weak and non-significant ($R = 0.11$, $p = 0.33$), indicating that the chromatin remodeling response in regenerating PNS neurons does not predict the magnitude of TCI-induced changes in corticospinal neurons.

**(D)** Top: Venn diagram comparing upregulated DA peaks following dorsal column axotomy (DCA; a non-regenerative CNS injury model; n=127) with TCI-upregulated DA peaks (n=3,705). Only 12 peaks (0.3%) are shared, with 115 (3.0%) unique to DCA and 3,693 (96.7%) unique to TCI. Bottom: Scatter plot of log2FC values for the 12 shared peaks in DCA vs Lam (x-axis) versus TCI-Injured (y-axis), with labeled loci including *Enc1*, *Abhd6*, *Ms4a6c*, *Pla2g4a*, *Nefh*, *Gpr17*, *Kcng2*, and *Med4*. Pearson correlation shows a positive trend but remains non-significant ($R = 0.45$, $p = 0.079$), suggesting a modest but statistically unreliable tendency for loci opened in DCA to also show increased accessibility following TCI.

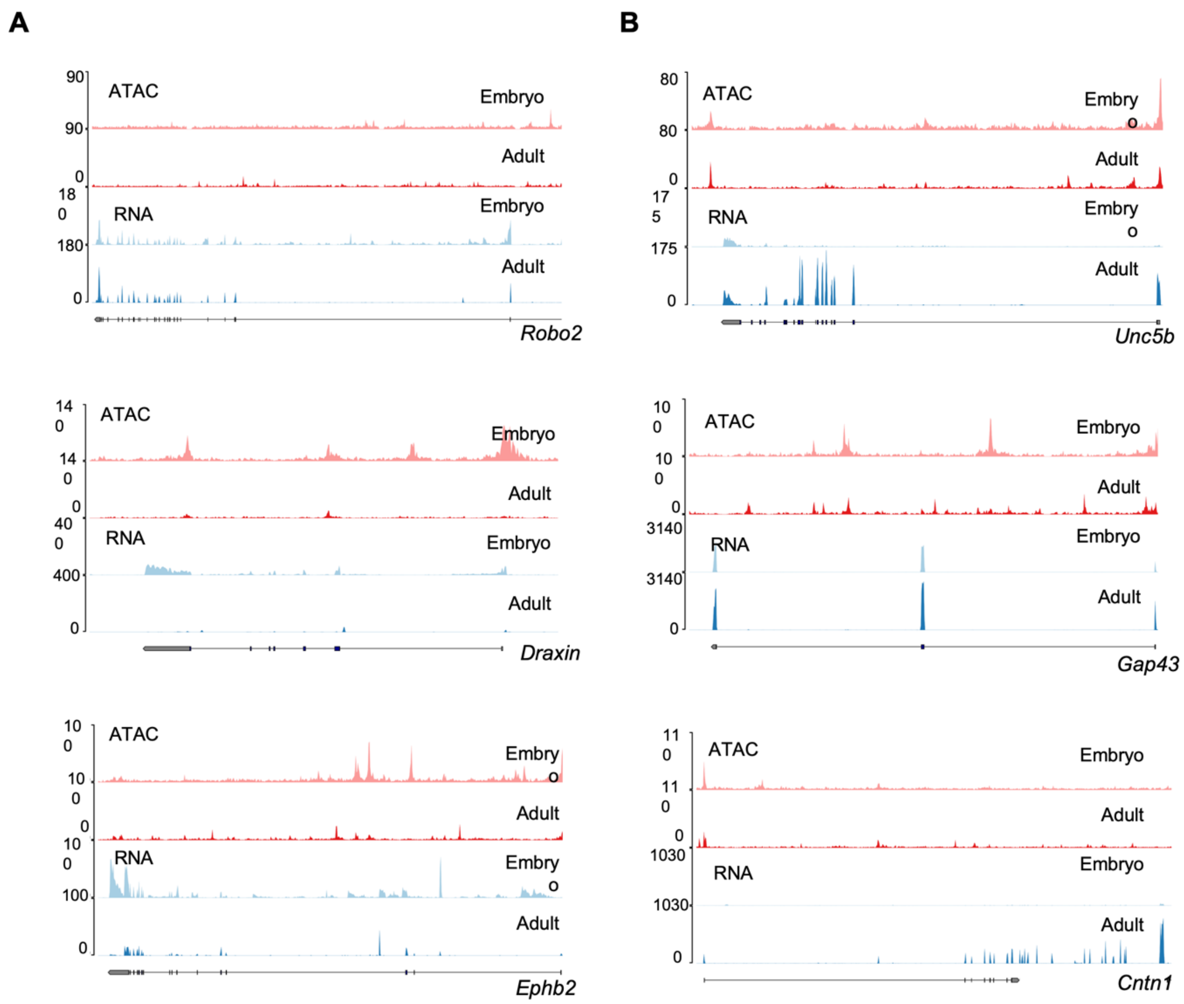


**Supplementary Figure 22. Chromatin accessibility and gene expression at selected axon guidance loci in embryonic and adult cortex.**

**(A)** Adult-retained loci. *Robo2* and *Ephb2* maintain accessible chromatin in adult despite reduced RNA. *Draxin* retains promoter accessibility but adult RNA is near-absent, indicating a permissive but transcriptionally silent state.

**(B)** Divergent chromatin–RNA loci. *Unc5b* shows concordant adult promoter accessibility and adult RNA, consistent with ongoing adult function. *Gap43* exhibits near-complete loss of adult ATAC signal yet retains substantial adult RNA, suggesting non-promoter-driven adult expression. *Cntn1* maintains adult chromatin accessibility with the highest adult RNA among all genes examined.

*Note : IGV browser tracks showing bulk ATAC-seq (red) and RNA-seq (blue) signal across embryonic (pooled E11–E18) and adult mouse cortex. Each locus displays four tracks: ATAC Embryo, ATAC Adult, RNA Embryo, RNA Adult. Y-axis scales are identical between embryonic and adult tracks within each gene.*

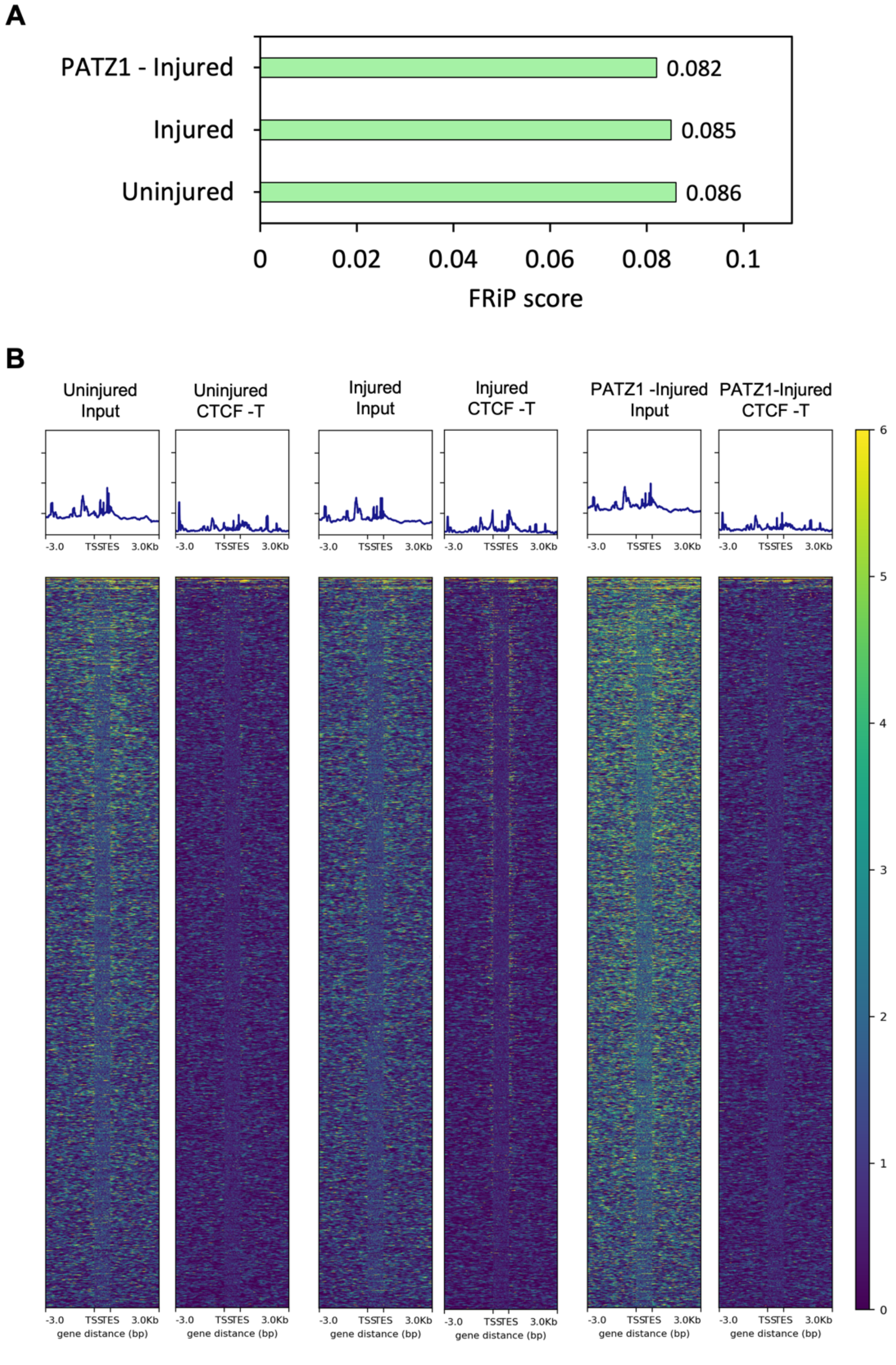


**Figure S23. CUT&RUN library quality metrics for CTCF profiling.**

**(A)** Fraction of reads in peaks (FRiP) scores for CTCF CUT&RUN libraries across uninjured (0.086), injured (0.085), and PATZ1-injured (0.082) conditions, confirming consistent and comparable enrichment across all three experimental groups.

**(B)** TSS enrichment profiles (top) and genome-wide signal heatmaps (bottom) for Input and CTCF CUT&RUN libraries (CTCF-T) in uninjured, injured, and PATZ1-injured conditions, plotted ±3 kb around transcription start sites. CTCF antibody libraries show enrichment at TSS-proximal regions relative to matched Input controls, confirming antibody specificity and library quality across all conditions.

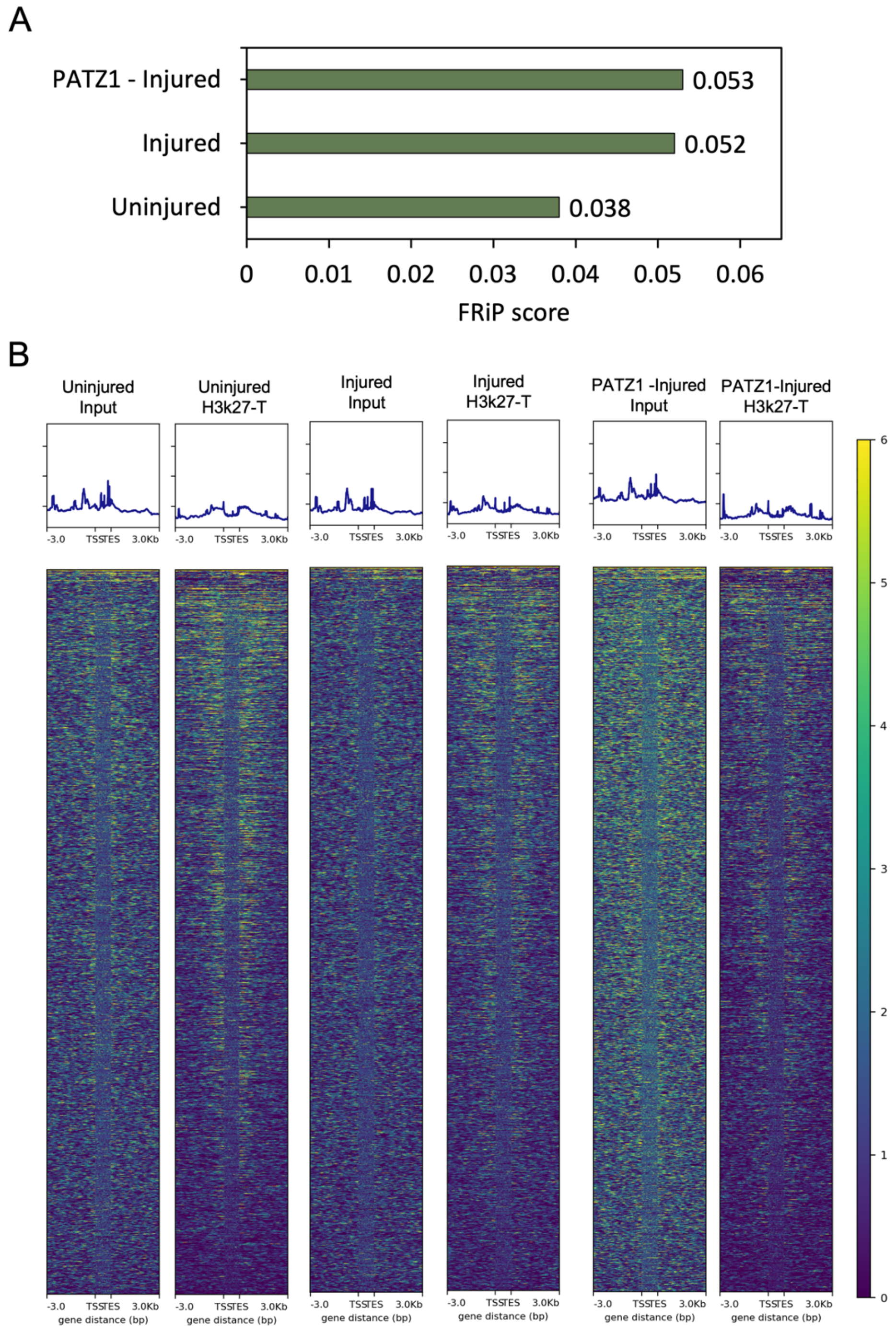


**Figure S24. CUT&RUN library quality metrics for H3K27ac profiling.**

**(A)** Fraction of reads in peaks (FRiP) scores for H3K27ac CUT&RUN libraries across uninjured (0.038), injured (0.052), and PATZ1-injured (0.053) conditions. The increase in FRiP score in injured and PATZ1-injured samples relative to uninjured cortex is consistent with increased genome-wide H3K27ac deposition following injury and PATZ1 overexpression.

**(B)** TSS enrichment profiles (top) and genome-wide signal heatmaps (bottom) for Input and H3K27ac CUT&RUN libraries (H3K27ac-T) in uninjured, injured, and PATZ1-injured conditions, plotted ±3 kb around transcription start sites. H3K27ac antibody libraries show characteristic enrichment at transcription start sites relative to matched Input controls, confirming antibody specificity and successful library preparation across all conditions.
